# Native mass spectrometry of membrane protein-lipid interactions in different detergent environments

**DOI:** 10.1101/2024.06.27.601044

**Authors:** Smriti Kumar, Lauren Stover, Lie Wang, Hanieh Bahramimoghaddam, Ming Zhou, David H. Russell, Arthur Laganowsky

## Abstract

Native mass spectrometry (MS) is revealing the role of specific lipids in modulating membrane protein structure and function. Membrane proteins solubilized in detergents are often introduced into the mass spectrometer; however, commonly used detergents for structural studies, such as dodecylmaltoside, tend to generate highly charged ions, leading to protein unfolding, thereby diminishing their utility for characterizing protein-lipid interactions. Thus, there is a critical need to develop approaches to investigate protein-lipid interactions in different detergents. Here, we demonstrate how charge-reducing molecules, such as spermine and trimethylamine-N-oxide, enable characterization of lipid binding to the bacterial water channel (AqpZ) and ammonia channel (AmtB) in complex with regulatory protein GlnK in different detergent environments. We find protein-lipid interactions are not only protein-dependent but can also be influenced by the detergent and type of charge-reducing molecule. AqpZ-lipid interactions are enhanced in LDAO (n-dodecyl-*N*,*N*-dimethylamine-N-oxide), whereas the interaction of AmtB-GlnK with lipids is comparable among different detergents. A fluorescent lipid binding assay also shows detergent dependence for AqpZ-lipid interactions, consistent with results from native MS. Taken together, native MS will play a pivotal role in establishing optimal experimental parameters that will be invaluable for various applications, such as drug discovery, as well as biochemical and structural investigations.

## Introduction

Native MS is a widely employed method for investigating the structure and function of biomolecular assemblies.^1-3^ By carefully adjusting the experimental conditions, it can preserve and probe non-covalent interactions, enabling quantitative analysis of binding events and determining stoichiometry of protein complexes.^4-6^ Native MS has proven invaluable in studying biomolecule interactions with small molecules like lipids, drugs, and nucleotides, which are essential in biomedical research. Remarkably, examining membrane protein complexes that transport ions and drugs across the cellular membrane has benefited from native MS analysis.^7-9^ Within the membrane environment, lipids have been recognized as critical regulators of membrane protein structure and function.^10-12^

Detergent micelles are typically employed to solubilize membrane proteins for native MS studies.^13^ Minimal collisional energy is applied to release the membrane protein from the detergent micelle.^14^ However, detergents commonly used for structural studies of membrane proteins, such as decylmaltoside (DM) and dodecylmaltoside (DDM), require higher collisional energies to dissociate from the protein.^14^ Consequently, membrane proteins acquire high charge states, resulting in the complex acquiring substantial internal energy, that often leads to protein unfolding.^14,15^ Charge-reducing detergents, such as C8E4 (tetraethylene glycol monooctyl ether) and LDAO (n-dodecyl-*N*,*N*-dimethylamine-N-oxide), have been discovered to circumvent this issue.^13,14,16^ Charge-reducing detergents facilitate the production of low-charged ions (with respect to non-charge-reducing detergents), preserving the protein’s native-like structure and non-covalent interactions.^17^ Additionally, More importantly, charge reduction increases the peak spacing, reducing the likelihood of mass spectral peak overlap and improving resolution for higher-order ligand-bound states.^18,19^

Various native MS approaches have been developed to generate charge-reduced ions using nanoelectrospray ionization (nanoESI).^20-22^ One strategy involves the addition of charge-reducing molecules.^18,23^ Trimethylamine-N-oxide (TMAO), a natural osmolyte, has been shown to effectively lower the average charge of proteins, enabling the analysis of higher-order lipid-bound states.^19,24,25^ Polyamines, such as spermine (SPM) and spermidine, are more potent and can effectively reduce protein charge at a much lower concentration than TMAO.^26^ SPM-derived detergents have recently been tailored for native MS studies, and they can significantly lower the charge state of membrane protein complexes.^27^ In contrast, other charge-reducing molecules like imidazole have exhibited only minor charge reduction and suffer from significant adduction, leading to poorly resolved mass spectra.^23,28^

An open question is how are membrane protein-lipid interactions influenced in different detergent environments? To this end, we conducted native MS experiments on the bacterial Aquaporin Z (AqpZ) and AmtB-GlnK (a complex between the ammonia channel AmtB and soluble regulatory protein GlnK) in various detergent environments: DM (n-decyl-ß-maltoside), OGNG (octyl glucose neopentyl glycol), NG (n-nonyl-ß-D-glucopyranoside), C8E4 (tetraethylene glycol monooctyl ether), and LDAO (lauryl dimethylamine N-oxide). To preserve non-covalent interactions, we used two charge-reducing molecules, SPM and TMAO, which have been shown to reduce charge states for membrane proteins in non-charge-reducing detergents significantly. We systematically characterized the interaction of five different lipids with both membrane proteins in various detergent environments. We also performed lipid titrations in different detergent environments to determine equilibrium dissociation constants (K_d_s) to provide a critical assessment of protein-lipid interactions. These studies are complemented by a fluorescent lipid binding assay.^29^ This work provides critical insight into the impact of detergents on membrane protein-lipid interactions.

## Materials and Methods

### Expression and Purification of AqpZ

Aquaporin Z (AqpZ, UniProt P60844) from *Escherichia coli* was expressed and purified as previously described.^18^ AqpZ containing a C-terminal Strep-tag II (AqpZ-STII) was expressed in *E. coli* BL21-A1 (Invitrogen). A single colony was used to inoculate 100 mL of LB (IBI Scientific) and grown overnight at 37°C while shaking. The overnight culture was used to inoculate TB (IBI Scientific) and allowed to grow at 37°C until the culture reached an OD_600_ of 0.6. Arabinose was added to a final concentration of 0.2%, and the culture was grown overnight at 20°C while shaking. Cells were harvested by centrifugation at 5,000g for 10 min, resuspended in lysis buffer (150 mM sodium chloride, 50 mM Tris pH 7.4 at room temperature), lysed with 4-5 passes through an M-110P microfluidizer (Microfluidics) operating at 25,000 PSI, and the lysate clarified by centrifugation at 20,000g for 25 min at 4°C. Crude membranes were pelleted by centrifugation for 2 hrs at 100,000g at 4°C. The membrane pellet was resuspended in extraction buffer (100 mM sodium chloride, 20% glycerol, 20 mM Tris pH 7.4 at room temperature) and extracted overnight with 5% octyl glucoside (OG). The supernatant was filtered and loaded onto a StrepTrap HP 5mL column (Cytiva) pre-equilibrated with SPNHA-DDM buffer (100 mM sodium chloride, 10% glycerol, 0.025% DDM, and 20 mM Tris pH 7.4 at room temperature). After loading, the column was washed with 20 mL of SPNHA-DDM supplemented with 2% OG, followed by 25 mL SPNHA-DDM until a steady baseline was reached. AqpZ was eluted with SPNHB-DDM (100 mM sodium chloride, 10% glycerol, 3 mM D-desthiobiotin, 0.025% DDM, and 20 mM Tris pH 7.4 at room temperature), concentrated using a 50 kDa MWCO concentrator (Millipore), and loaded onto a Superdex 200 Increase 10/300 GL column (GE Healthcare) equilibrated with SPNHC-C8E4 buffer (100 mM sodium chloride, 10% glycerol, 0.5% C8E4, and 50 mM Tris pH 7.4 at room temperature). The peak fractions containing C8E4-solubilized AqpZ were aliquoted, flash-frozen in the liquid nitrogen, and stored at -80°C before use.

### Expression, Purification, and Cy3 Labelling of AqpZ-STII-GC

The AqpZ-STII expression construct was modified to include a Gly-Cys following the C-terminal Strep-tag II sequence (AqpZ-STII-GC). The protein was expressed and purified as described for AqpZ-STII. The only exception was the wash buffer was modified to include 1 mM dithiothreitol (DTT). The eluted protein was mixed with DDM to a final concentration of 0.1% and mixed with a ten-fold excess of Cy3 Maleimide (Click Chemistry Tools, stock dissolved in DMSO). The labeling reaction proceeded for 2 hours at room temperature. The labelling reaction was quenched by the addition of 1mM DTT, and DDM was added to a final concentration of 0.5% prior to loading onto a HiPrep 26/10 Desalting column (Cytiva) pre-equilibrated with SPNHA-DDM buffer. Peak fractions containing DDM-solubilized Cy3 Maleimide labeled AqpZ-STII-GC were aliquoted, flash-frozen in the liquid nitrogen, and stored at -80°C.

### AmtB-GlnK Complex Purification

AmtB-GlnK complex was purified as previously described.^5^ In brief, GlnK containing an N-terminal HRV3C protease cleavable Strep-tag II and maltose binding protein (STII-TEV-MBP-HRV3C-GlnK) was expressed from pCOLA in *E. coli* BL21(DE3) (New England Biolabs). Protein expression was induced with 0.1 mM IPTG and grown overnight at 20°C. 8×His-HRV3C-AmtB was expressed from pCDF in *E. coli* BL21-A1 (Invitrogen), induced with 0.2% arabinose and grown overnight at 20°C. Cell harvesting, resuspension, and lysis for both proteins were identical to that described for the purification of AqpZ. Post lysis, the lysate containing GlnK was clarified by centrifugation at 40,000g for 20 mins, filtered, and loaded onto a 5 mL MBPTrap HP column (Cytiva) equilibrated with MBP-loading buffer (100 mM sodium chloride, 10% glycerol, and 20 mM Tris; pH 7.4 at room temperature). The protein was eluted with MBP-elution buffer (100 mM sodium chloride, 10% glycerol, and 20 mM Tris, 10 mM maltose; pH 7.4 at room temperature) concentrated and loaded onto a HiLoad 16/600 Superdex 200 pg column (GE Healthcare) equilibrated in MBP-loading buffer. Membrane isolation and detergent extraction steps for AmtB were identical to AqpZ’s. AmtB was first purified using HisTrap HP 5 mL column (Cytiva) pre-equilibrated with NHA-DDM-1 buffer (200 mM sodium chloride, 10% glycerol, 0.025% DDM, 20 mM imidazole, and 20 mM Tris; pH 7.4 at room temperature). After elution with NHB-DDM buffer (100 mM sodium chloride, 10% glycerol, 0.025% DDM, 500 mM imidazole, and 20 mM Tris; pH 7.4 at room temperature), the pooled sample was loaded onto a HiPrep 26/10 Desalting column pre-equilibrated with NHA-DDM-2 buffer (100 mM sodium chloride, 10% glycerol, 0.025% DDM, 20 mM imidazole, and 20 mM Tris; pH 7.4 at room temperature). Purified tagged AmtB and GlnK were mixed at a molar ratio 1:3 supplemented with 1 mM adenosine diphosphate (ADP) and HRV3C protease (50:1 protein to protease ratio). The mixture was incubated overnight at 4°C and loaded onto a drip column packed with Ni-NTA (IBA Biosciences) pre-equilibrated with NHA-DDM-2 buffer. The flow through was collected and loaded onto a Superdex 200 Increase 10/300 GL column equilibrated with SPNHC-C8E4 buffer supplemented with 1 mM ADP. Peak fractions containing AmtB-GlnK were collected, flash-frozen in liquid nitrogen, and stored at -80°C.

### Detergent Exchange

AqpZ-STII was loaded onto a drip column packed with Streptactin Sepharose agarose (IBA Biosciences) pre-equilibrated with SPNHC-C8E4 buffer. After loading, the column was washed with 10 column volumes (CV) of SPNHC-C8E4, followed by a 10 CV wash of SPNHC-wash buffer (100 mM sodium chloride, 10% glycerol, and 50 mM Tris pH 7.4 at room temperature) supplemented with 2x CMC (critical micelle concentration) of the desired detergent. The protein was eluted with 3 mM D-desthiobiotin in SPNHC-wash buffer containing 2x CMC of the desired detergent. AqpZ-STII-GC was detergent-exchanged similarly, except that SPNHC-DDM was used instead of SPNHC-C8E4. The AmtB-GlnK complex was detergent-exchanged using a Superdex 200 Increase 10/300 GL column equilibrated with SPNHC-wash buffer supplemented with 2x CMC of the desired detergent and 1 mM ADP.

### Sample Preparation for Native Mass Spectrometry (MS) Analysis

Purified membrane proteins were buffer exchanged into 200 mM ammonium acetate (pH 7.4 at room temperature adjusted with ammonium hydroxide) supplemented with 2x CMC of the desired detergent using a centrifugal desalting column (Micro Bio-Spin 6 columns, BioRad). For the AmtB-GlnK complex, 100 µM ADP was added to the MS buffer. All the phospholipids, including 18:1 Cy5 Cardiolipin (1,1’,2,2’-tetraoleoyl cardiolipin-N-(Cyanine 5)) were purchased from Avanti Polar Lipids and prepared as previously described.^19,30^ Charge-reducing reagents, spermine, and trimethylamine N-oxide (TMAO) were purchased from Alfa Aesar and Cayman Chemical, respectively. Lipids, charge-reducing reagents, and buffer-exchanged protein were prepared in aqueous ammonium acetate and incubated for two to five minutes. The optimized concentrations of AqpZ, AmtB-GlnK complex, spermine, and TMAO were 1 µM, 2 µM, 5 mM, and 60 mM, respectively. For samples without charge-reducing reagents, the same volume of aqueous ammonium acetate was used instead of charge-reducing reagents as previously described.^19^

### Native Mass Spectrometry (MS)

Samples were loaded into a gold-coated borosilicate nanoelectrospray ionization emitters prepared in-house^13^ at room temperature and introduced into an Exactive Plus EMR Orbitrap Mass Spectrometer (Thermo Scientific). The instrument was optimized for each sample, and detailed instrument settings can be found in Tables S1 and S3. Native MS spectra were processed using UniDec^31^ with the following settings: m/z range 2000 to 20000, charge range 5-30, mass sampling every 1 Da, and peak FWHM of 0.85. The weighted average states (Z_avg_) were computed using UniDec.

### Fluorescence Resonance Energy Transfer (FRET) Lipid Binding Assays

AqpZ-STII-GC labelled with Cy3 served as the donor (530 nm excitation, 580 nm emission) and 18:1 Cy5 Cardiolipin (620 nm excitation, 675 nm emission) as the acceptor. FRET (530 nm excitation, 675 nm emission) measurements and correction factors were calculated as previously described.^32,33^ Protein and lipid were both at a final concentration of 0.5 µM. SPNHC-wash buffer with a high concentration of detergent was mixed to make samples containing 10x CMC of detergent. The total protein, lipid, and buffer volume was 50 µL, and the experiments were performed at room temperature in a black 384-well plate (NUNC). Measurements were recorded on a CLARIOstar microplate reader (BMG LABTECH).

## Results

### AqpZ in different detergent environments

We first investigated AqpZ in different detergent environments **(Figures 1 and S1-S2)**. For AqpZ solubilized in DM, only signals corresponding to those of DM micelles were observed under various instrument settings **(Figure S1 and Table S1)**. In OGNG, peaks were observed for detergent micelles along with monomer, trimer, and tetramer distributions of AqpZ **(Figure 1C)**. The presence of monomer and trimer signals results from the activation of the complex under these conditions, leading to the dissociation of the intact tetrameric complex. The mass spectrum of the channel in NG had signals for the tetrameric complex **(Figure S2C)**. Despite comparatively mild instrument conditions, signals for dissociated products (monomer and trimer) were also observed but to a lesser extent than that observed for the channel in OGNG **(Figure 1C, S2C, and Table S1)**. The presence of dissociated species presents challenges to studying membrane protein-lipid interactions since conditions that favor subunit dissociation also support the dissociation of non-covalently bound lipids. In contrast, AqpZ in either C8E4 or LDAO, both of which are charge-reducing detergents,^14,16^ showed only signal for the intact tetrameric complex with a reduction (up to four) in the average charge state **(Figures S2 and Table S2)**. These results illustrate how charge-reducing detergents preserve the integrity of non-covalent interactions.

**Figure 1.**
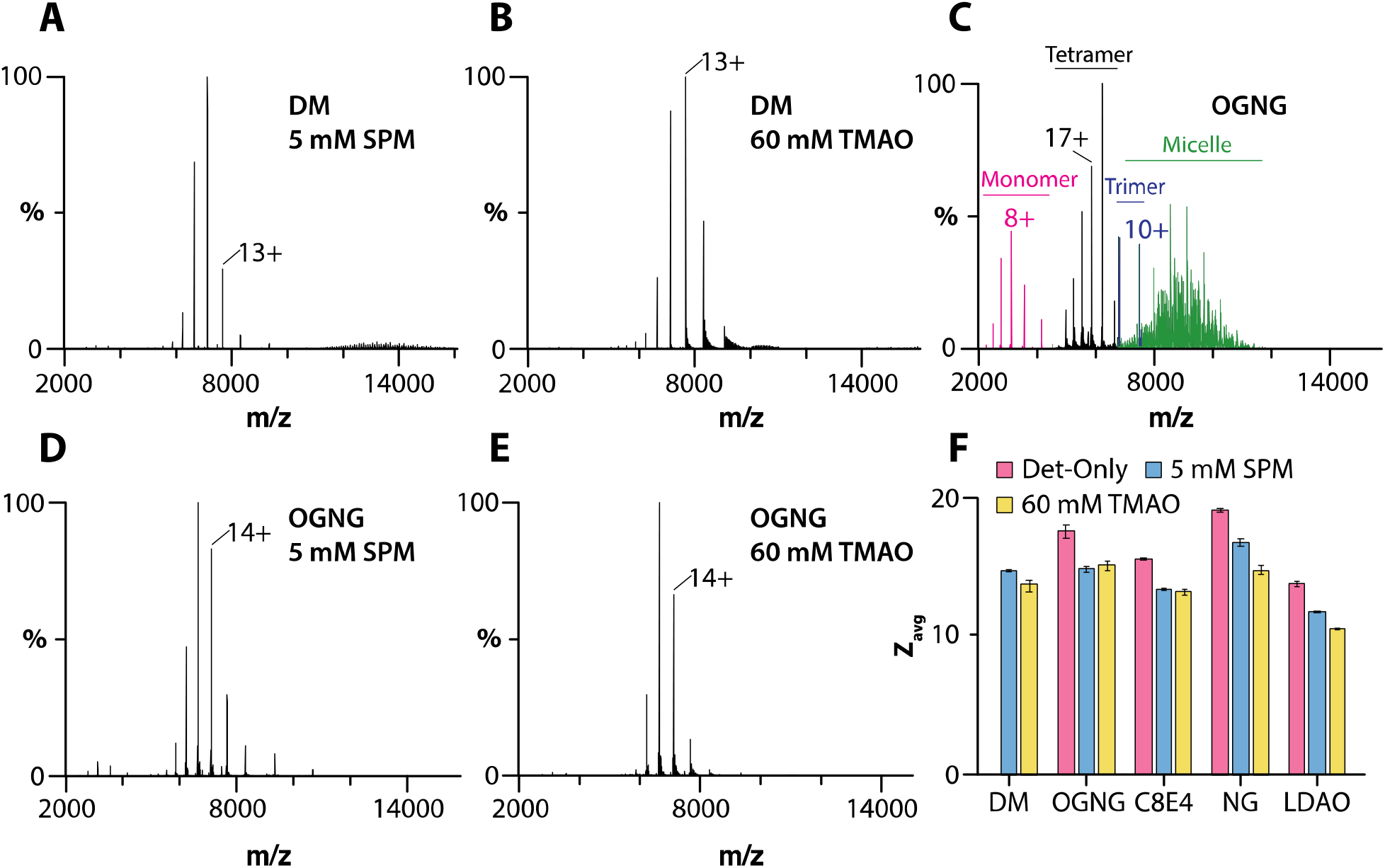
SPM and TMAO preserve the tetramer of AqpZ in different detergent environments. A-B) Mass spectra of 1 μM AqpZ in DM in the presence of A) 5mM spermine and B) 60 mM TMAO. C-E) Mass spectra of 1 μM AqpZ in OGNG and in the presence of D) 5mM spermine and E) 60 mM TMAO. F) The average charge state (Z_avg_) of AqpZ in different detergent environments and the presence of charge-reducing molecules. Reported are the mean and standard deviation (*n*=3).

We next explored the utility of charge-reducing molecules, such as spermine and TMAO, to preserve the AqpZ complex in various detergents. For the studies presented herein, spermine and TMAO were added to a final concentration of 5 mM and 60 mM, respectively. These concentrations were found to be optimal in terms of balancing charge reduction and signal intensity. Adding either of the charge-reducing molecules to AqpZ in DM displayed a well-resolved mass spectrum of the tetrameric complex with an average charge state (Z_avg_) of ∼14 **(Figures 1A-B, 1F, and Table S2)**. Likewise, in the case of AqpZ in OGNG and NG, the presence of both spermine and TMAO demonstrated lower charged states (Z_avg_ ranging from 15 to 16) and enhanced protein complex stability **(Figures 1D-E, S2F, S2I, and Table S2)**. Adding spermine and TMAO to AqpZ in charge-reducing detergents C8E4 and LDAO reduced Z_avg_ by approximately three charges, with Z_avg_ ranging from 11 to 13 **(Figures 1F and S2D-E, S2G-H)**. In short, the addition of SPM and TMAO enables the opportunity to characterize AqpZ-lipid interactions in non-charge-reducing detergents.

### AmtB-GlnK in different detergent environments

Analogous experiments were performed for the AmtB-GlnK in different detergent environments and instrumental conditions **(Figures S3-S4 and Table S3)**. Dissociated species of the AmtB-GlnK complex were observed in DM and OGNG **(Figure S3)**. In the case of NG, the intact AmtB-GlnK complex was observed **(Figure S4B)**. In general, the addition of TMAO to AmtB-GlnK in different detergents led to the dissociation of the complex **(Figures S3B-D)**. Mass spectra for the intact AmtB-GlnK complex were obtained in most detergents when spermine was present **(Figures S4 and Table S4)**. In several instances, we observed the adduction of charge-reducing molecules and detergents to the protein complex. TMAO formed a significant number of adducts with the protein in OGNG **(Figure S3G)**. These TMAO adducts could be dissociated at higher collision energies, but this also resulted in the dissociation of the complex **(Figures S3B-D)**. Another interesting observation was the case of AmtB-GlnK in C8E4 and TMAO, where the appearance of detergent adducts was noted **(Figure S3H)**. The detergent environments that displayed adduction were omitted from the studies that followed. In the studies we follow, the intact complex was obtained notably in the charge-reducing detergents C8E4 and LDAO, even without charge-reducing molecules **(Figures S4D and S4F)**.

### Cardiolipin-protein interactions

The ability to preserve the intact membrane protein complexes in different detergent environments (described above) inspired us to investigate membrane protein-lipid interactions **(Figures 2-3 and S5)**. We first focused on cardiolipin, which has been reported to modulate the water transport activity of AqpZ.^16^ The native mass spectrum of AqpZ solubilized in DM with 25 equivalents of 18:1 cardiolipin (TOCDL, 1,1’,2,2’-tetraoleoyl-cardiolipin) and in the presence of spermine or TMAO displayed a similar distribution of TOCDL-bound states **(Figures 2A-C)**. In the case of AqpZ solubilized in OGNG or NG, the presence of TMAO gave rise to an increase in TOCDL-bound states compared to SPM, despite both conditions displaying a similar Z_avg_ **(Figures 2D-G and S5G-H)**. In the case of charge-reducing detergents, the addition of spermine or TMAO did not significantly influence the number of TOCDL molecules bound to AqpZ **(Figures 2H-I and S5A-F)**. For the AmtB-GlnK complex (2 µM) with TOCDL (25 µM) in different detergent environments, TOCDL binding was more prominent in LDAO and DM **(Figures 3C-D)**. The addition of spermine did not change the abundance of TOCDL bound to AmtB-GlnK in C8E4, LDAO, and NG (**Figures 3A-B and 3D**). These results show that the binding of TOCDL to both membrane protein complexes is directly influenced by the detergent environment.

**Figure 2.**
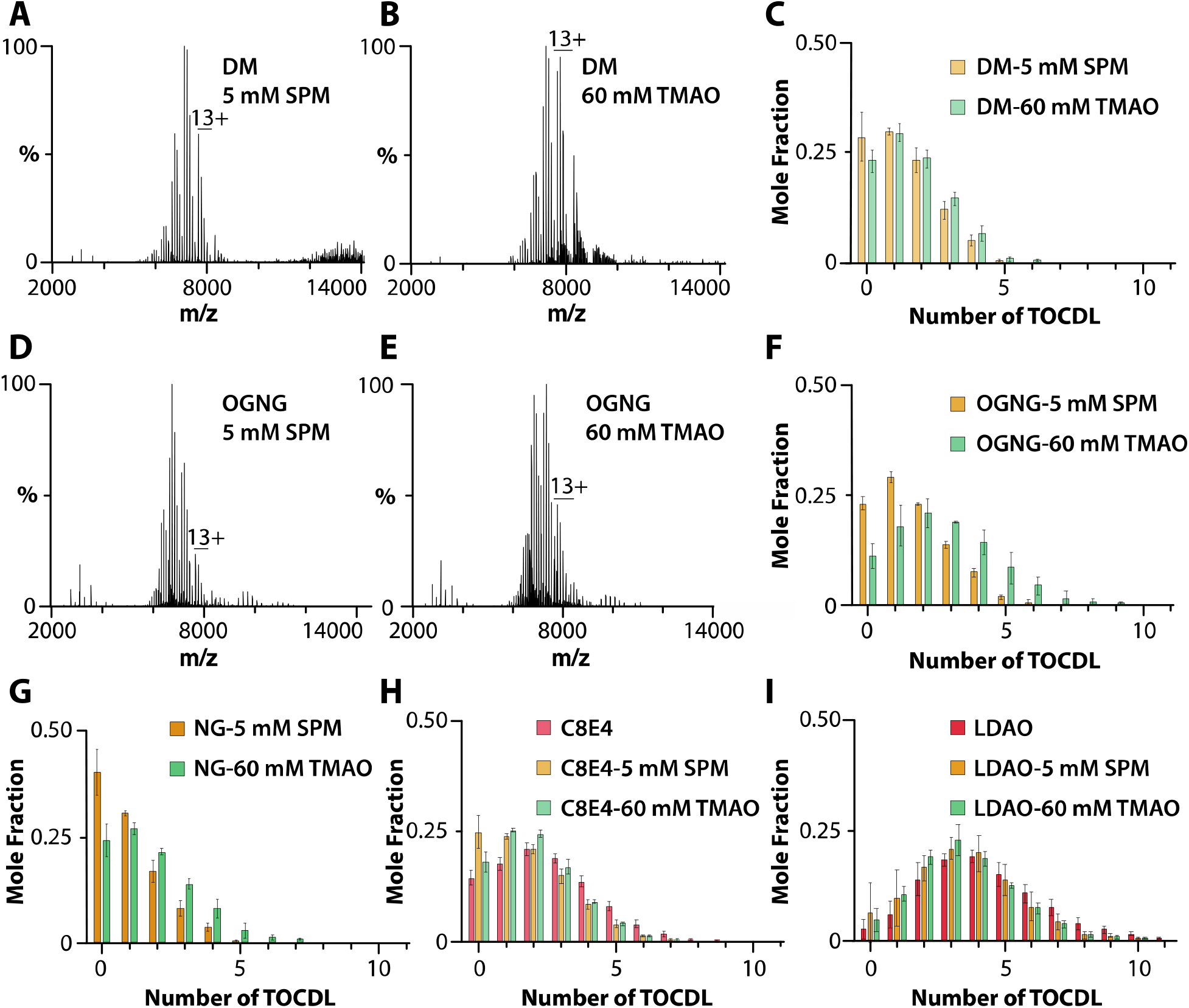
Characterization of AqpZ-TOCDL interactions in different detergent environments. (A-B) AqpZ (1 μM) in DM mixed with 25 μM TOCDL and in the presence of A) 5 mM spermine and B) 60 mM TMAO. C) Mole fraction plot of AqpZ-TOCDL species determined from the deconvolution of the mass spectra shown in A and B. (D-E) AqpZ in OGNG mixed with 25 equivalents of TOCDL in the presence of D) 5 mM spermine and E) 60 mM TMAO. F-I) Mole fraction plots for AqpZ and in complex with TOCDL in F) OGNG, G) NG, H) C8E4, and I) LDAO. Reported are the mean and standard deviation (*n*=3).

**Figure 3.**
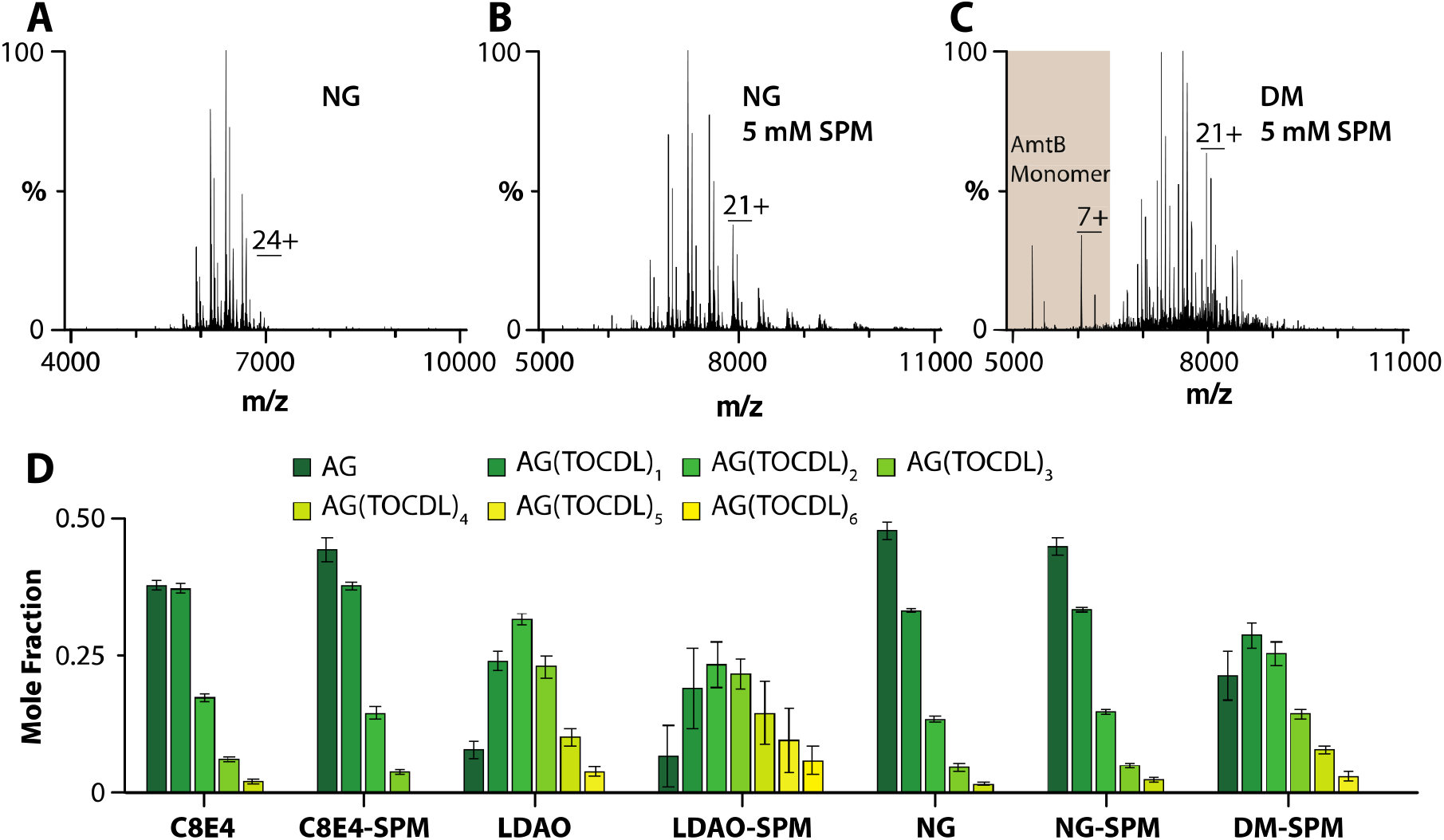
TOCDL binding to AmtB-GlnK in different detergents. A-B) Mass spectra of 2 μM AmtB-GlnK mixed with 25 μM TOCDL in NG and in the presence of B) 5 mM spermine. C) AmtB-GlnK solubilized in DM with TOCDL and 5 mM spermine. D) Mole fraction plots of AmtB-GlnK in complex with TOCDL in different detergent environments. Reported are the mean and standard deviation (*n*=3).

### Phosphatidylethanolamine-protein interactions

We next investigated the binding of 1-palmitoyl-2-oleyl phosphatidylethanolamine (POPE, 16:0-18:1), a zwitterionic phospholipid, to AqpZ and AmtB-GlnK in different detergent environments **(Figure 4 and S6)**. Similar to the experiments for TOCDL, the membrane protein, and POPE concentrations were held at a fixed concentration. In DM, up to ten POPE molecules bound to AqpZ with similar abundances were observed in the presence of either spermine or TMAO **(Figures S6A-B and S6M)**. In the case of AqpZ solubilized in OGNG and NG, the addition of spermine or TMAO resulted in a similar number of POPE molecules bound to the membrane protein complex **(Figures S6C-F and S6N-O)**. However, the mole fraction of higher POPE-bound states of AqpZ was pronounced for the detergent conditions containing TMAO. In C8E4, up to ten POPE molecules bound to AqpZ were observed, and this detergent with TMAO slightly skewed the abundance for a subset of AqpZ-POPE stoichiometries **(Figures S6G-H, S6K, and S6P)**. Interestingly, AqpZ in LDAO bound up to17 POPE molecules and was independent of the presence or absence of charge-reducing molecules **(Figures S6I-J, S6L, and S6Q)**. The variation in the POPE bound states of AmtB-GlnK in different detergent environments was less pronounced **(Figure 4)**. For LDAO and DM, higher POPE-bound states were more abundant, although the total number of POPE-bound states varied slightly **(Figures 4C-D)**. AmtB-GlnK in DM with spermine bound up to nine POPE molecules **(Figure 4D)**. Adding spermine to NG increased the number of POPE bound to AmtB-GlnK **(Figure 4D)**. Like AqpZ, POPE binding to AmtB-GlnK was independent of spermine in charge-reducing detergents C8E4 and LDAO **(Figure 4D)**.

**Figure 4.**
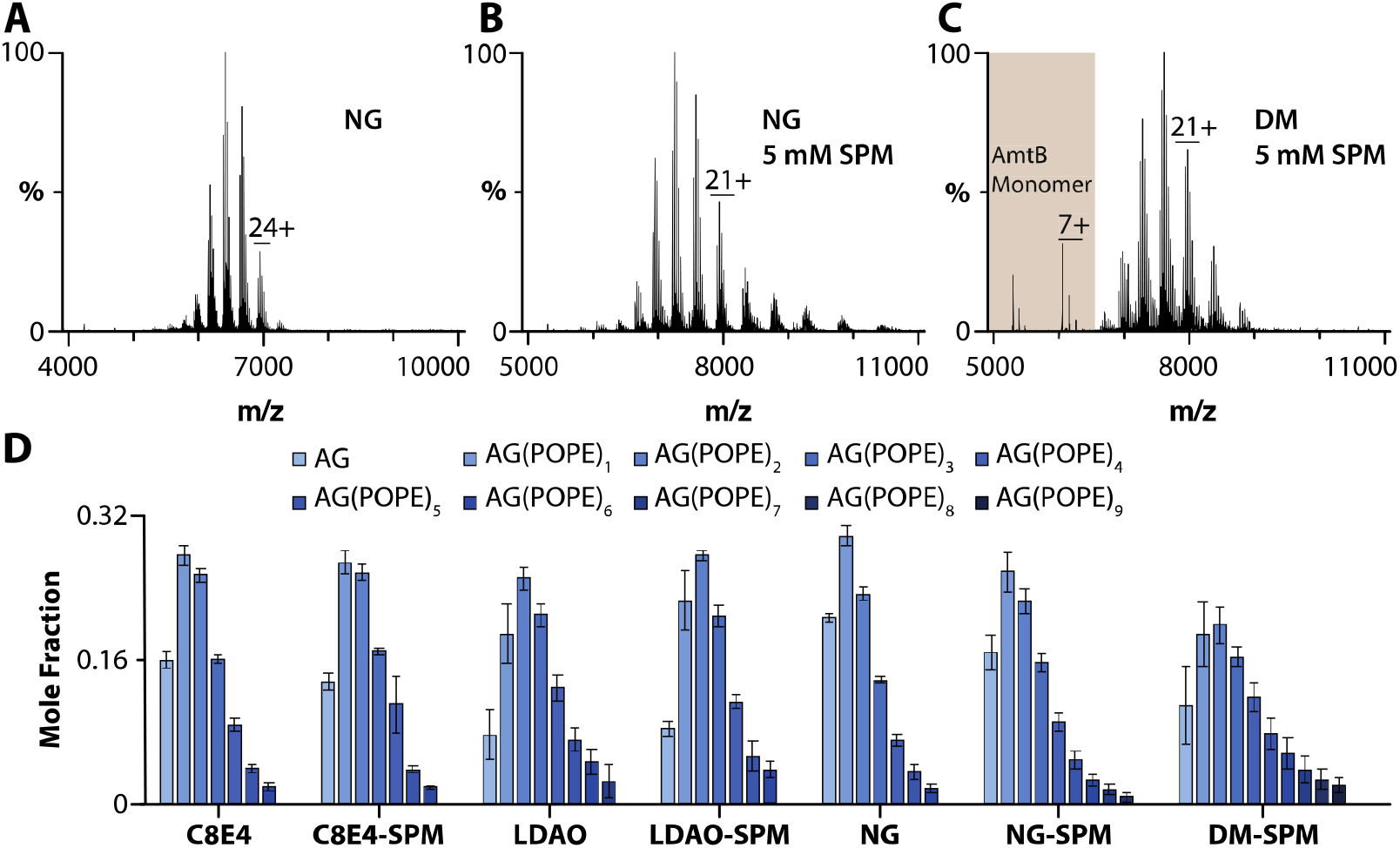
Characterization of POPE binding to AmtB-GlnK in different detergent environments. Mass spectra of 2 μM AmtB-GlnK mixed with 50 μM POPE in A) NG, B) NG with 5 mM spermine, C) DM with 5 mM spermine, D) Mole fractions determined from the deconvolution of the mass spectra of AmtB-GlnK with 50 μM POPE in different environments. Reported are the mean and standard deviation (*n*=3).

### Phosphatidylglycerol-protein interactions

The binding of the anionic phospholipid, 1-palmitoyl-2-oleyl phosphatidylglycerol (POPG, 16:0-18:1) to membrane proteins was also investigated **(Figure S7-S8)**. AqpZ in DM supplemented with spermine or TMAO showed up to ten POPG molecules binding the channel **(Figures S7A-B and S7M)**. Unlike the other lipids, the presence of TMAO promoted a higher abundance of AqpZ-POPG bound states in DM compared to the same condition with SPM. Similarly, AqpZ in OGNG promoted POPG binding to AqpZ in the presence of TMAO over SPM **(Figures S7C-D and S7N)**. AqpZ solubilized in NG bound up to 8 lipids, and the abundance of the different states between the two charge-reducing molecules was comparable **(Figures S7E-F and S7O)**. POPG binding to AqpZ in C8E4 was comparable to that in NG and independent of charge-reducing molecules **(Figures S7G-H, S7K, S7P)**. Like C8E4, AqpZ-POPG interactions were independent of the charge-reducing molecule, but a significant increase in POPG binding (up to 20) was observed **(Figures S7I-J, S7L, and S7Q)**. Unlike AqpZ, AmtB-GlnK in different detergent environments showed comparable binding of POPG, and the addition of charge-reducing molecules did not significantly impact the abundance of lipid-bound states **(Figures S8)**.

### Other lipid-protein interactions

The other two lipids found in *Ecoli* membrane^34^ we investigated were 1-palmitoyl-2-oleyl phosphatidic acid (POPA, 16:0-18:1) and 1-palmitoyl-2-oleoyl-sn-glycero-3-phospho-L-serine (POPS, 16:0-18:1) **(Figure S9-S12)**. In general, AqpZ showed a comparable number of POPA and POPS bound to the channel in DM, OGNG, and NG **(FiguresS9-S10)**. In most cases, the mole fraction of higher AqpZ-lipid bound states in DM, OGNG, and NG was enhanced in the presence of TMAO. AqpZ solubilized in C8E4 showed a similar number of POPAs bound to the complex that was comparable to the protein in non-charge-reducing detergents and independent of charge-reduction **(Figures S9G-H, S9K, and S9P)**. In LDAO, AqpZ bound nearly two-fold the number of POPA molecules **(Figures S9I-J, S9L, and S9Q)**. An appreciable reduction in POPS binding to AqpZ in C8E4 was observed **(Figure S10G-H, S10K, and S10P)**. However, AqpZ in LDAO had a similar number of POPS-bound states to those obtained in non-charge-reducing detergents **(Figures S10I-J, S10L, and S10Q)**. For both C8E4 and LDAO, introducing charge-reducing molecules resulted in no significant change in the abundance of AqpZ-lipid bound states. POPA and POPS binding to AmtB-GlnK was broadly comparable for the different detergents **(Figures S11-12)**. A notable exception was enhanced lipid binding to AmtB-GlnK in LDAO. In the case of POPA, the addition of SPM to AmtB-GlnK in LDAO led to the dissociation of AmtB-GlnK to AmtB **(Figure S13)** and a reduction in the mole fraction of the higher order of POPS bound states **(Figure S12D)**.

### Determination of equilibrium binding dissociation constants

To gain additional insight into the influence of detergent on protein-lipid interactions, we determined equilibrium dissociation constants (K_d_s) for AqpZ-lipid interactions in different detergents containing either SPM or TMAO **(Figure 5 and S14-S42 and Table S5-S7)**. For example, mass spectra from a titration series of TOCDL were deconvoluted^31^ to determine the mole fraction of AqpZ(TOCDL) (Figure 5A-C). Subsequently, a sequential lipid binding model was used to determine K_d_s **(Figure 5C-D)**.^35^ Of the different detergent environments, AqpZ displayed the highest binding affinity for TOCDL in LDAO, with the K_d_ for binding the first TOCDL (K_d1_) ranging from 0.3 to 1.4 µM **(Figure 5D)**. The binding affinity for TOCDL decreased for AqpZ in the other detergents. The highest K_d_s for TOCDL were observed for AqpZ in C8E4 and NG with SPM. In the case of POPE and POPG, AqpZ in LDAO displayed the highest lipid binding affinity **(Figure 5E-F)**. Like TOCDL, POPE and POPG interactions with AqpZ showed a decrease in binding affinity, in which the K_d_s for AqpZ in C8E4 and NG displayed the weakest binding.

**Figure 5.**
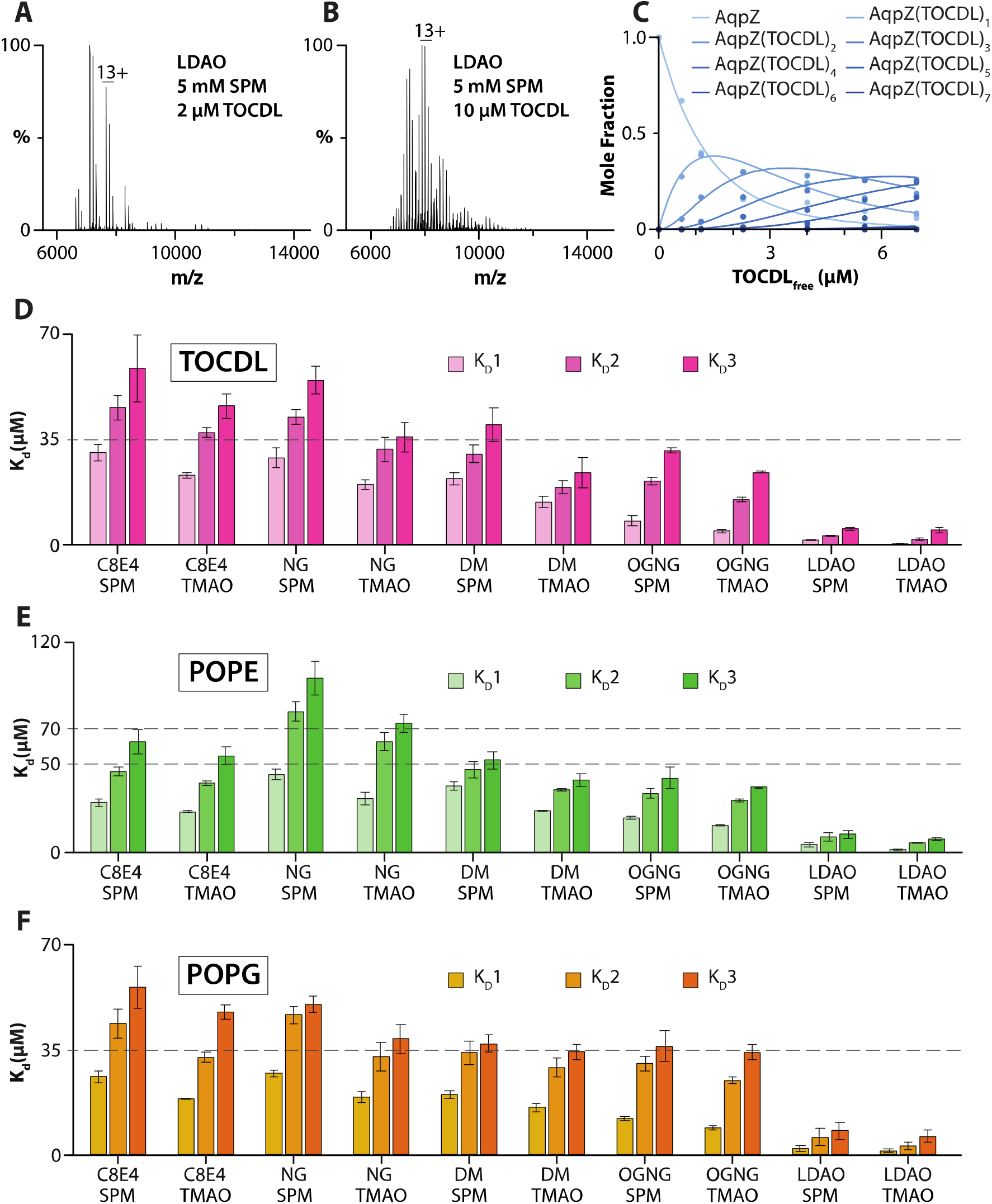
Determination of AqpZ-TOCDL equilibrium dissociation binding constants (K_d_) in different detergents. A-B) Representative mass spectra of AqpZ (1 μM) in LDAO mixed with SPM and different concentrations of TOCDL. The concentration of TOCDL is denoted in the inset. C) Plot of mole fraction data (dots) for AqpZ(TOCDL)0-7 determined from a titration series of TOCDL and subsequent fit (*R*^2^ = 0.99) of a sequential lipid binding model (lines). D-F) Plot of Kds for AqpZ binding, D) TOCDL, E) POPE, and F) POPG in different detergent environments. Reported are the mean and standard deviation (*n*=3).

### FRET-Based Lipid Binding Assay

To complement the native MS studies, we employed a soluble fluorescent lipid binding assay **(Figure 6)**.^29^ In these studies, the binding of TOCDL modified with a cyanine 5 fluorophore (Cy5CDL) to AqpZ labelled with cyanine 3 (AqpZCy3) is monitored by Förster Resonance Energy Transfer (FRET) measurements. An equimolar mixture of Cy5CDL and AqpZCy3 was evaluated in two different concentrations - two and three times the critical micelle concentration (CMC) - of the selected detergent **(Figure 5A)**. The largest FRET signal was observed for the sample in OGNG. In contrast, AqpZ in NG and C8E4 displayed the lowest FRET signals, indicating reduced binding of the fluorophore-modified lipid. In all cases, a five-fold increase in detergent concentration resulted in a significant reduction in the FRET signal, consistent with the higher concentration of detergent competing with the binding of Cy5CDL to AqpZCy3. We also assessed the impact of SPM and TMAO on lipid binding to AqpZ in OGNG. No statistical difference was observed among the different conditions **(Figure 5B**)

**Figure 6.**
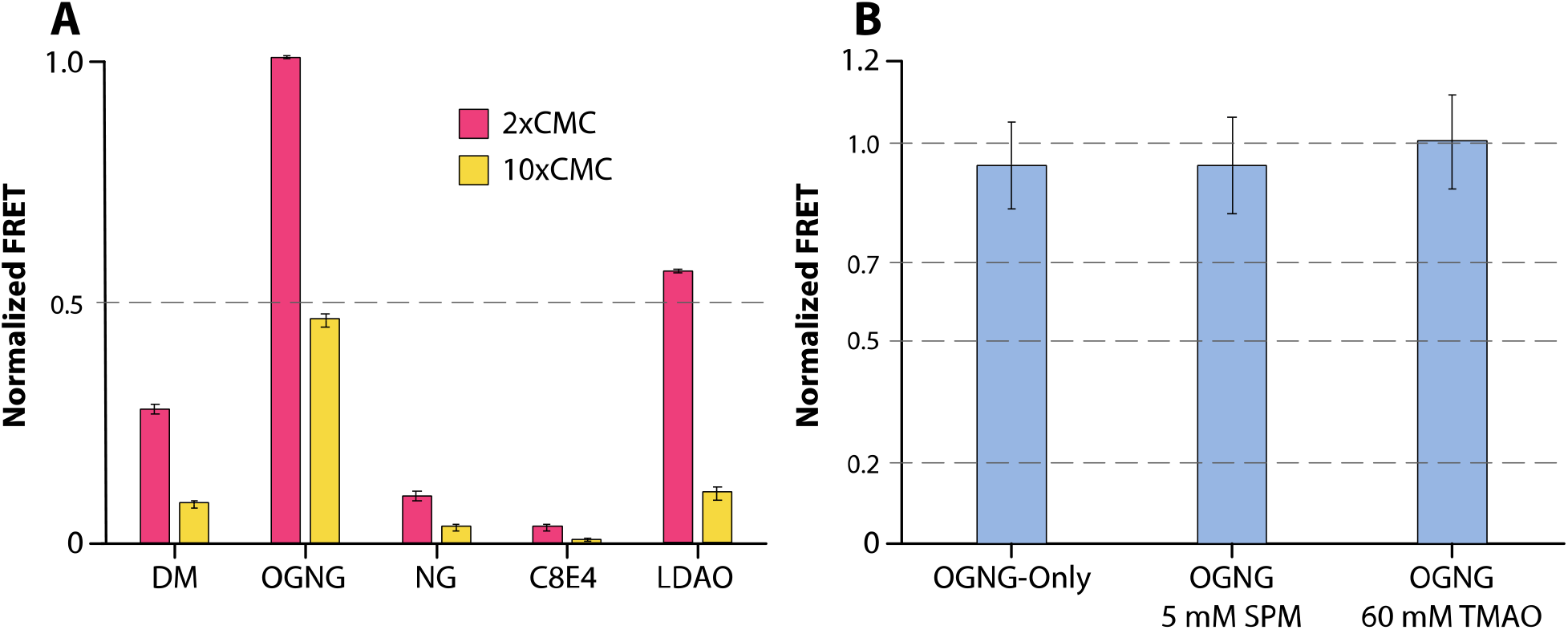
AqpZ binding to TOCDL modified with a fluorophore monitored by FRET. AqpZ modified with cyanine 3 (AqpZCy3) and fluorophore-modified TOCDL (Cy5CDL) were both held at a concentration of 0.5 μM. A) Plot of FRET for a mixture of AqpZCy3 and Cy5CDL in different detergents at two and ten times the critical micelle concentration (CMC). FRET data is normalized to the largest response. B) The influence of SPM and TMAO on AqpZCy3 and Cy5CDL interactions. Reported are the mean and standard deviation (*n*=3).

## Discussion

Membrane proteins are often purified using detergents, and the choice of detergent is usually determined by several criteria, such as biochemical stability and specific activity of the purified complex. As a result, the selected detergent will be dependent on the target membrane protein complex. Detergents commonly used for structural studies are often not particularly useful for native MS where the goal is to preserve non-covalent interactions.^14,16,36^ Here, mass spectra of AqpZ and AmtB-GlnK in DM, OGNG, and NG illustrate these environments result in conditions that require considerable collisional activation to liberate the membrane protein from the detergent micelle, which can also promote dissociation of the intact protein complexes. The discovery of charge-reducing detergents,^16,36^ such as C8E4 and LDAO used herein, provide conditions that support mass measurements of intact membrane protein complexes. While these detergents have proved useful for native MS studies, such as those characterizing the binding of lipids and other molecules, a potential problem is that these detergents may not be ideal for purifying various membrane protein complexes that support biochemical stability and activity.

Charge-reducing molecules have been found to be beneficial for promoting the stabilization of membrane proteins and the preservation of non-covalent interactions by promoting the production of ions with lower charge states. ^16,20-22,36^ The addition of SPM and TMAO to AqpZ and AmtB-GlnK in different detergents resulted in the reduction of Z_avg_ and mass measurements of intact protein complexes. The reduction in Z_avg_ depends on each component in the system **(Figure S43)**. The concentration of SPM and TMAO can be optimized to obtain the desired charge-reduction and signal, as done here. It is worth noting that in some cases, we observed the adduction of molecules that are dependent on the protein and detergent. For example, the AmtB-GlnK complex in OGNG with TMAO resulted in a poorly resolved mass spectrum due to the adduction of TMAO and OGNG. Another example is the AmtB-GlnK complex in C8E4 with TMAO, which resulted in the adduction of C8E4. Notably, AqpZ in these detergent environments did not suffer from adducts. We have previously reported that membrane protein solubilized in DDM displayed a broad mass spectrum in the presence of SPM and TMAO. However, the use of SPM-detergents can produce charge-reduced ions of membrane proteins in DDM.^27^ In short, the growing arsenal of charge-reducing molecules affords the opportunity to charge-reduce membrane proteins in different detergent environments.

Native MS reveals membrane protein-lipid interactions are not only dependent on the protein, but they can be influenced by the detergent environment. In the case of AqpZ, LDAO supports an environment that promotes the binding of various lipids. Moreover, lipid binding is largely independent of the addition of SPM and TMAO. On the other hand, AqpZ in the other detergents showed a similar number of lipids bound to the water channel. There are some notable exceptions where the addition of a charge-reducing molecule results in enhanced binding, such as for the binding of POPS to AqpZ in OGNG with TMAO. These results are consistent with the fluorescent lipid binding assay, in which OGNG and LDAO showed the most binding. Most interestingly, the binding of lipids to AmtB-GlnK in the different detergent environments displayed less variation as compared to AqpZ. More specifically, the binding of POPA was comparable for the complex in C8E4, LDAO, and DM. These results illustrate the detergent environment can influence lipid binding.

While the results from experiments using a fixed molar ratio of protein to lipid are illuminating, the determination and evaluation of K_d_s for AqpZ-lipid interactions provide a more quantitative assessment. Interestingly, AqpZ in LDAO displays the highest lipid binding affinity. AqpZ in NG and C8E4 consistently gave the highest K_d_ values, implying these detergents are more effective at competing with AqpZ-lipid interactions. The K_d_s for AqpZ-lipid interactions in different detergents will be a composite of lipid binding to AqpZ and the competition of detergent binding at the specific lipid binding site. Evaluation of the fold change in subsequent K_d_s, for example, K_d_2/K_d_1, shows that nearly all the detergent environments are comparable **(Figure S44)**. In contrast, LDAO, which supported higher lipid binding affinity to AqpZ, displayed the largest fold change in binding affinity for the binding of subsequent lipids. This result may suggest the LDAO may promote specific binding of the first lipid compared to other detergents.

Detergents are commonly classified into “harsh” and “mild” categories, depending on their capacity to preserve the integrity of membrane structures,^37-41^ which may be the result of removing or preserving essential protein-lipid interactions. Interestingly, LDAO, a zwitterionic detergent considered to be “harsh”, promotes the interaction of AqpZ with various lipids. For example, AqpZ-TOCDL interactions are enhanced by more than 20-fold compared to that in DM, a detergent considered to be “mild”. In contrast, the interaction of lipids with AmtB-GlnK was largely independent of the detergent environment. While the categorization of detergents is largely anecdotal, this report demonstrates how different combinations of detergents with varying CMC **(Table S8)** and charge-reducing molecules can be exploited for native MS studies of membrane proteins. It is also important to emphasize that these different detergent environments open up an exciting opportunity to study a broader range of membrane proteins, especially those that are not biochemically stable or active in charge-reducing detergents. Taken together, native MS will be instrumental in defining experimental conditions that support high-affinity binding of ligands, such as lipids and other molecules, to membrane proteins that will be of critical importance for drug discovery and biochemical and structural studies.

## Author Contributions

S.K. and A.L. designed the research. S.K. and L.S. expressed and purified the protein. S.K. performed the experiments. S.K., L.S., D.R., and A.L. analyzed the data. S.K. and A.L. wrote the manuscript with input from other authors.

## Acknowledgments

The work was supported by National Institutes of Health (NIH) under grant numbers (R01GM121751, R01GM139876, R01GM138863, RM1GM145416, and RM1GM149374).

## Notes

The authors declare no competing financial interest.

## TOC

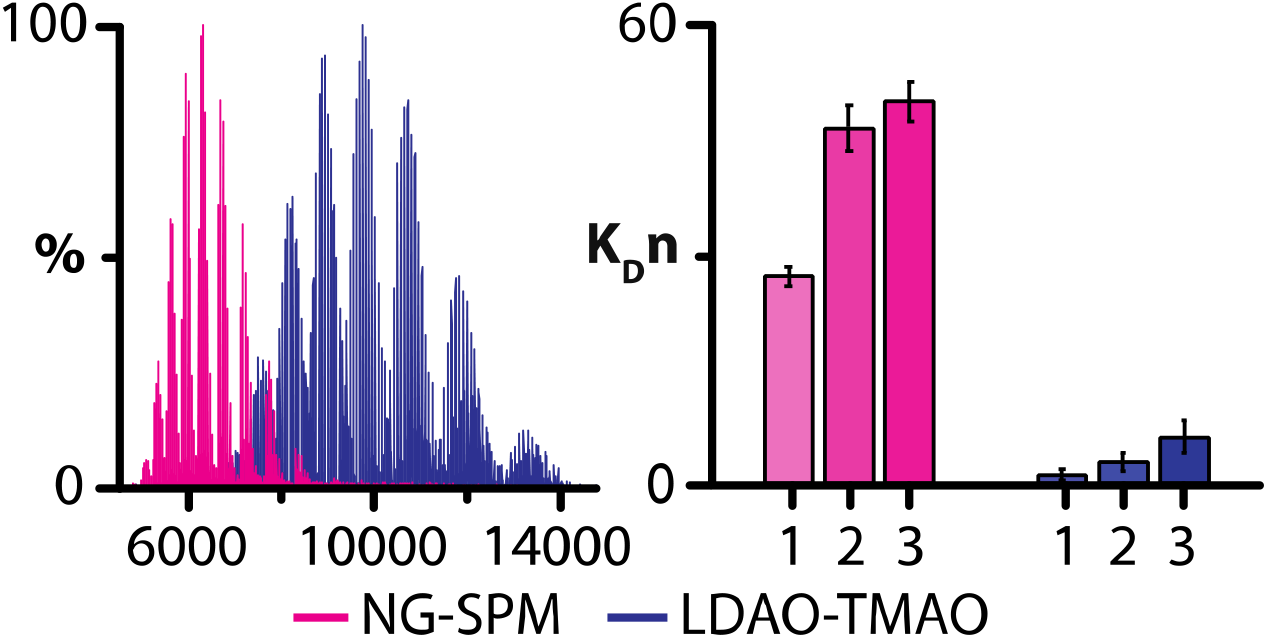

