## Supplemental Information for "Native mass spectrometry of membrane protein-lipid interactions in different detergent environments"

**Supporting Figures**

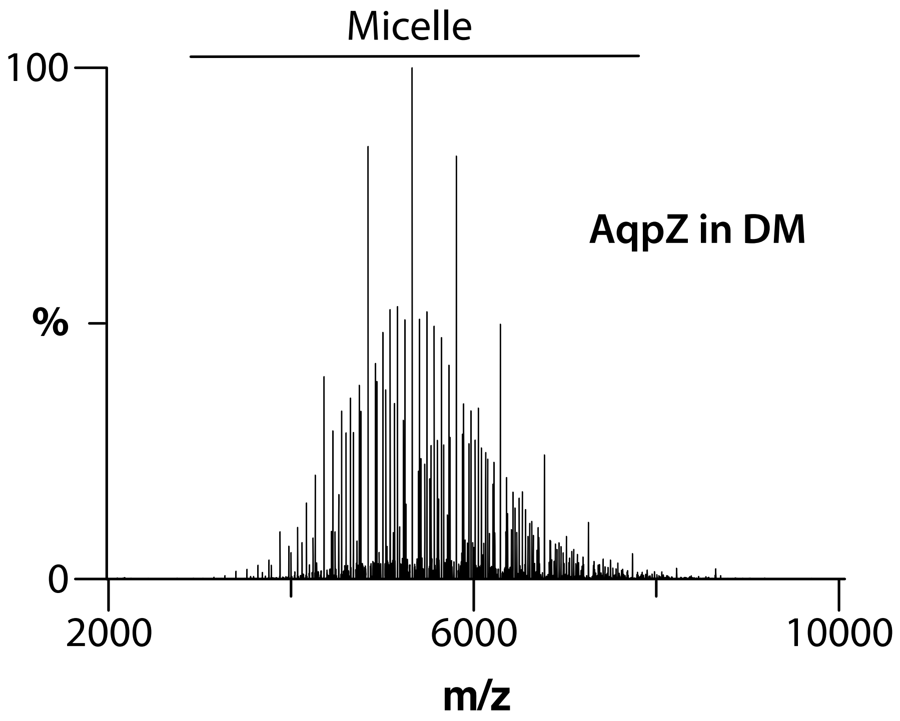


**Figure S1. Mass spectrum of AqpZ (1 µM) in DM**. The mass spectrum corresponds to DM micelles with no signal for AqpZ.

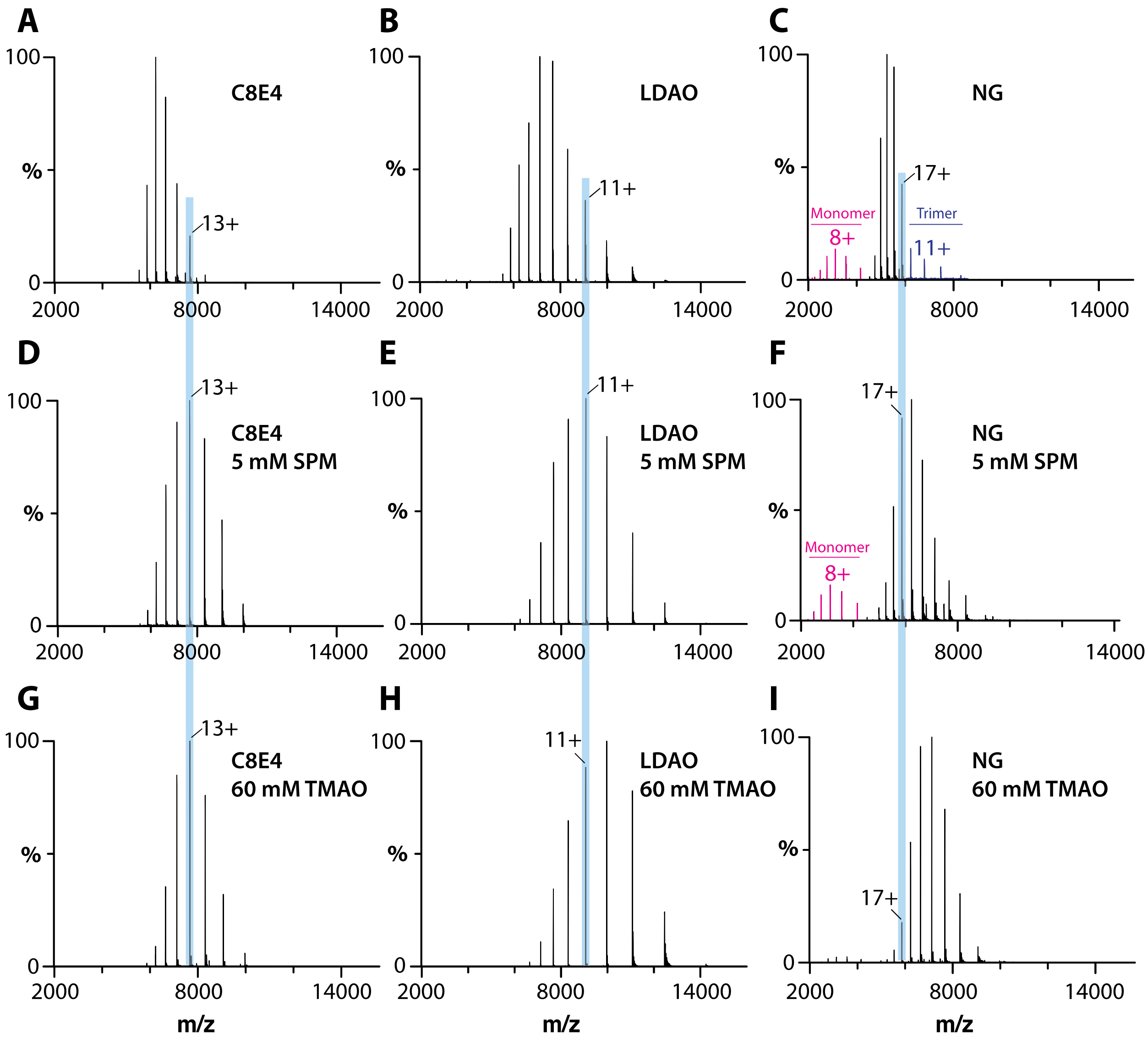


**Figure S2. Mass spectra of AqpZ in different detergents with SPM and TMAO**. Mass spectra of 1 μM AqpZ in various detergent environments. The detergent and charge-reducing molecule (if added) are labelled.

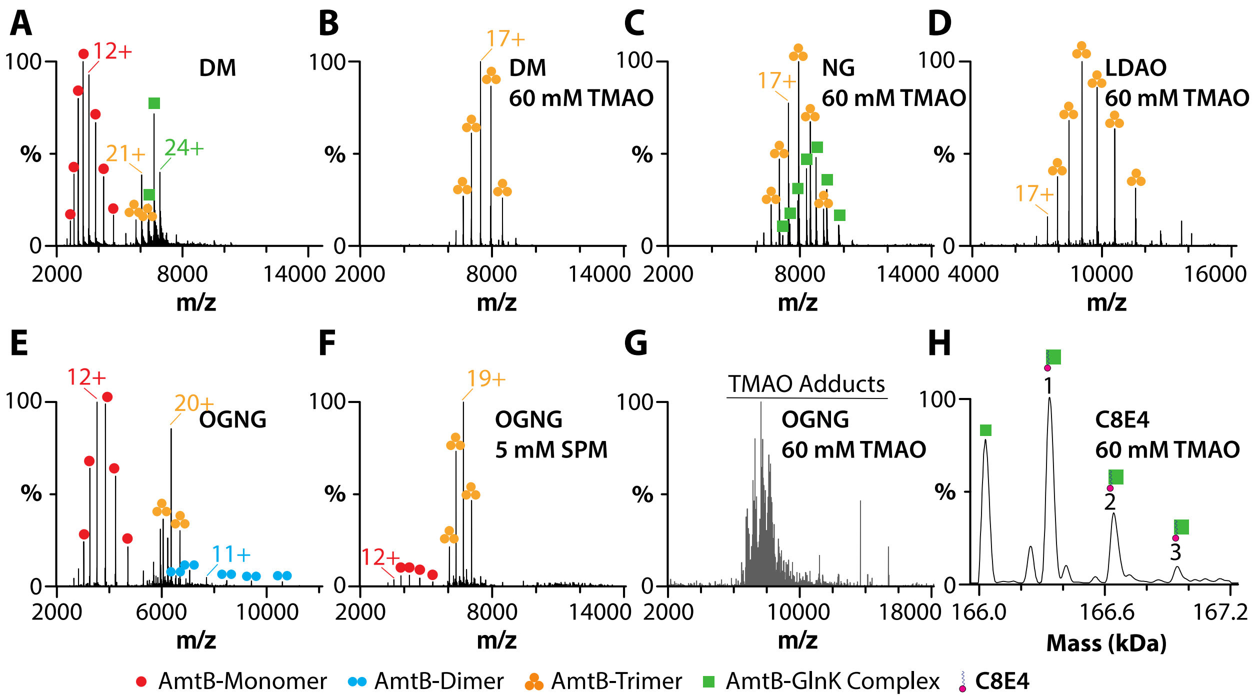


**Figure S3. AmtB-GlnK in different detergent environments**. The concentration of the AmtB-GlnK complex was 2 µM. Shown as described in Figure S2.

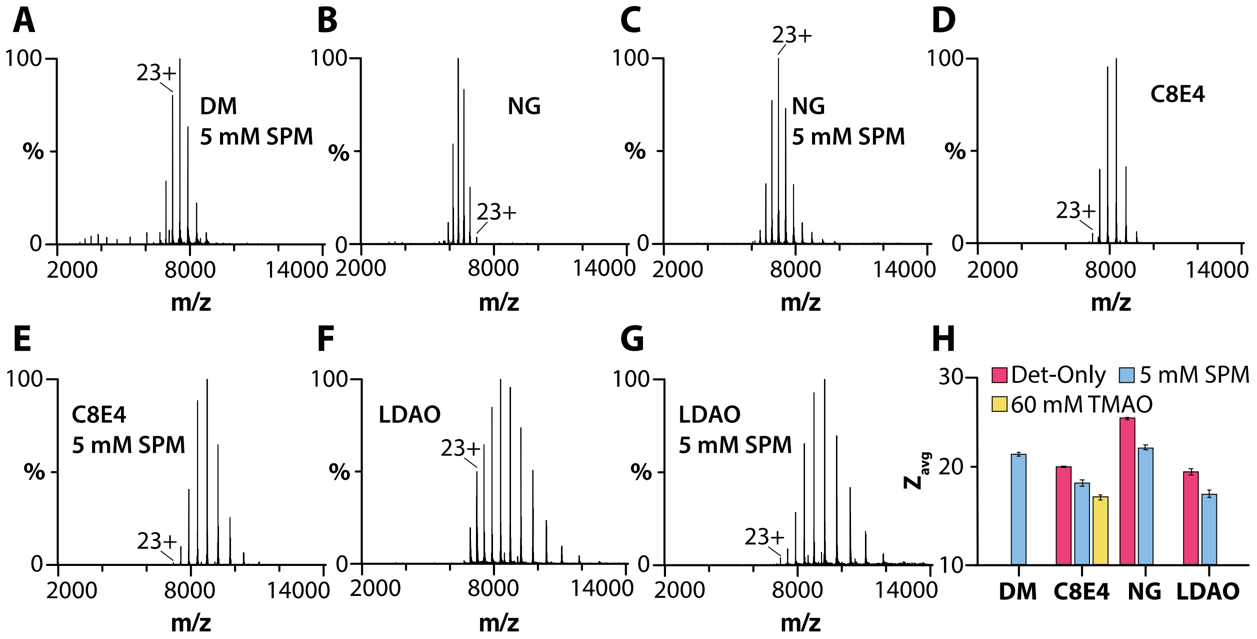


**Figure S4. AmtB-GlnK complex in different environments**. A-G) Mass spectra of 2 µM AmtB-GlnK in various detergent environments. Shown as described in Figure S1. H) Plot of $Z_{avg}$ for AqpZ in different detergent environments in the presence or absence of charge-reducing molecules. Reported are the mean and standard deviation (*n*=3).

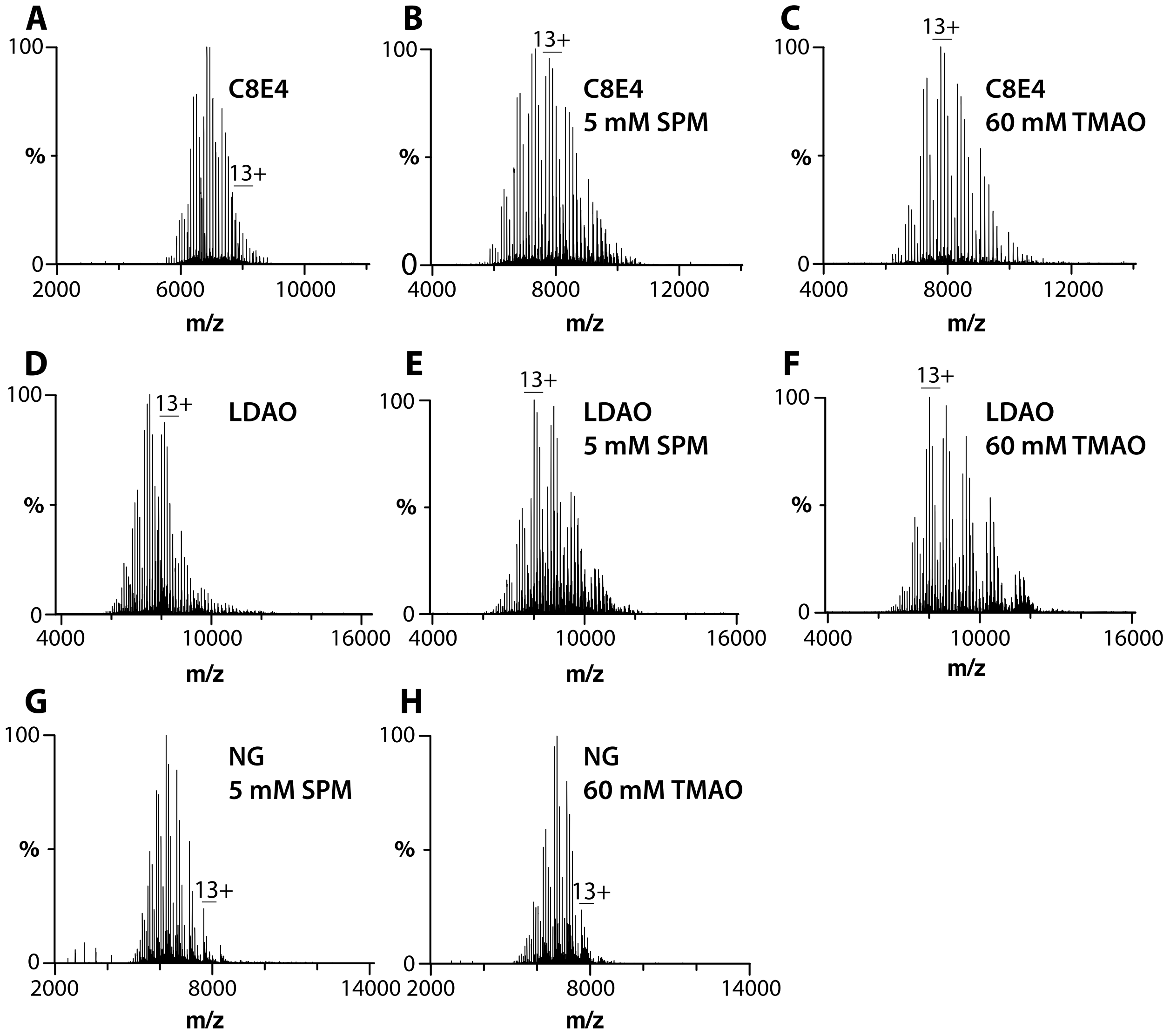


**Figure S5. TOCDL binding to AqpZ in different detergents.** Mass spectra of 1 µM AqpZ in different detergents mixed with 25 µM TOCDL. The detergent and charge-reducing molecule (if added) are labelled.

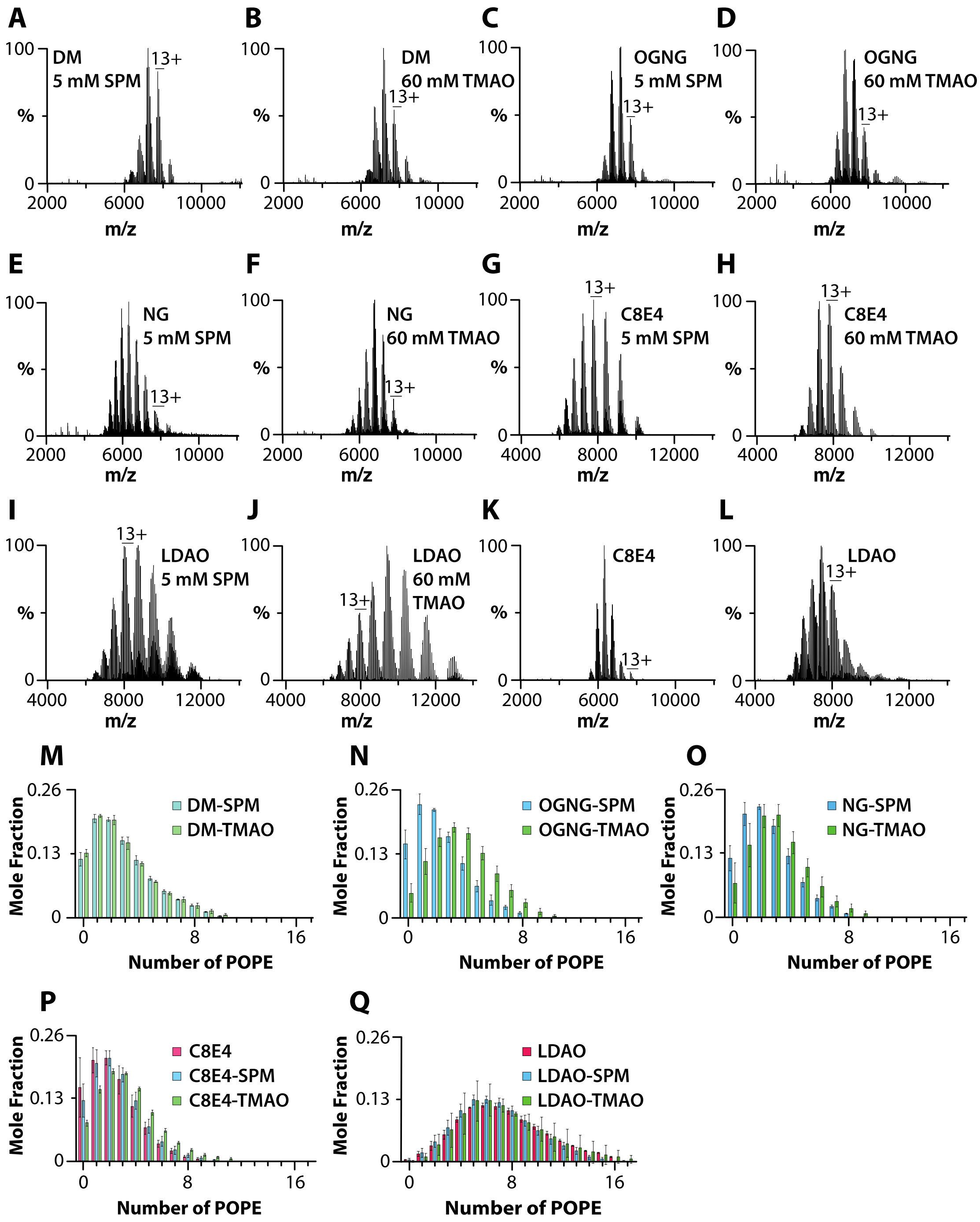


**Figure S6. AqpZ-POPE interactions in different detergents.** A-L) AqpZ (1 µM) mixed with 50 µM POPE. Shown as described in Figure S5. M-Q) Plot of the mole fraction for different species determined from the deconvolution of the mass spectra shown in A-L. Reported are the mean and standard deviation (*n*=3).

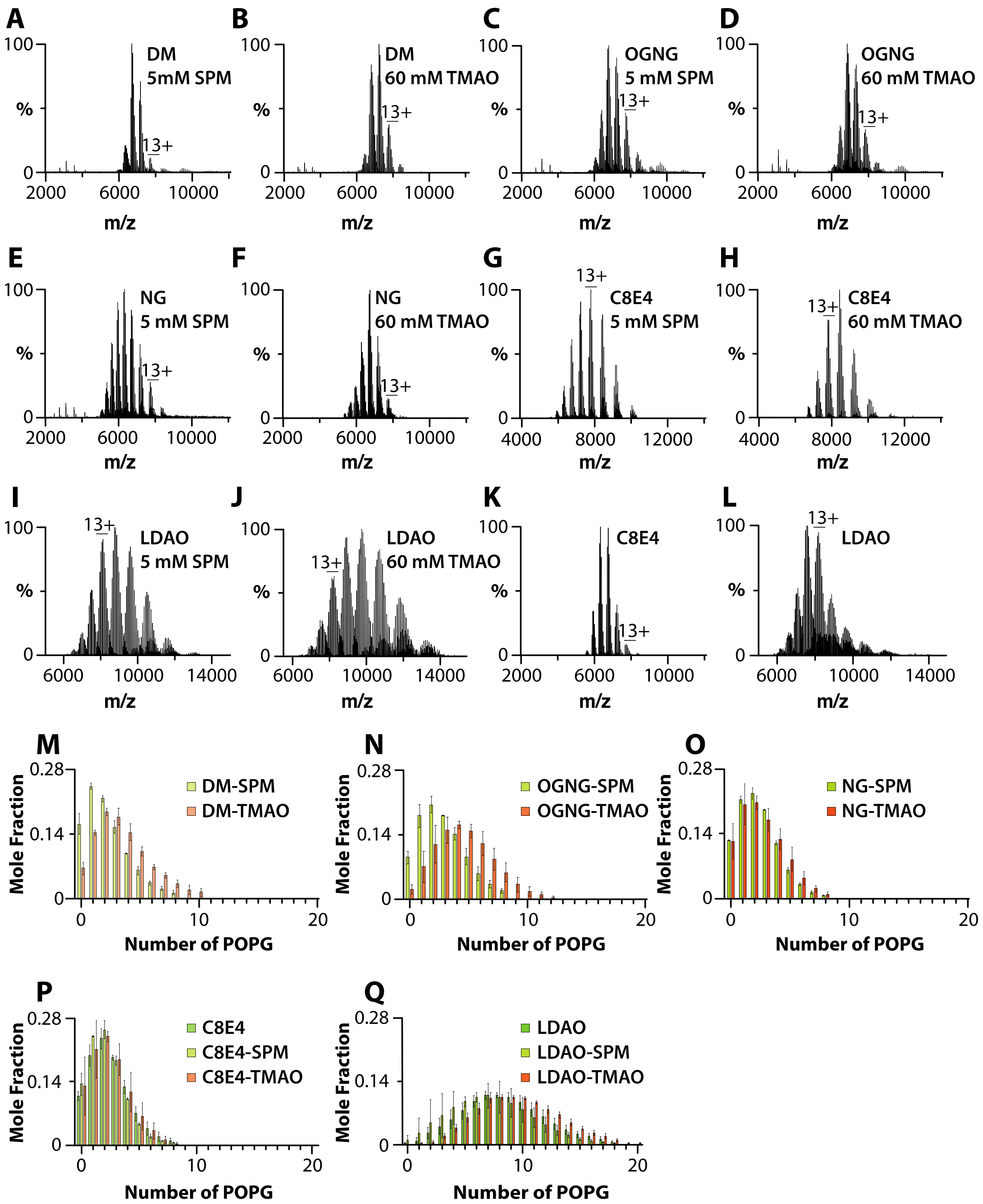


**Figure S7. AqpZ binding POPG in different detergents.** A-L) AqpZ (1 µM) mixed with 50 µM POPG. Shown as described in Figure S6. M-Q) Plot of the mole fraction for different species determined from the deconvolution of the mass spectra shown in A-L. Reported are the mean and standard deviation (*n*=3).

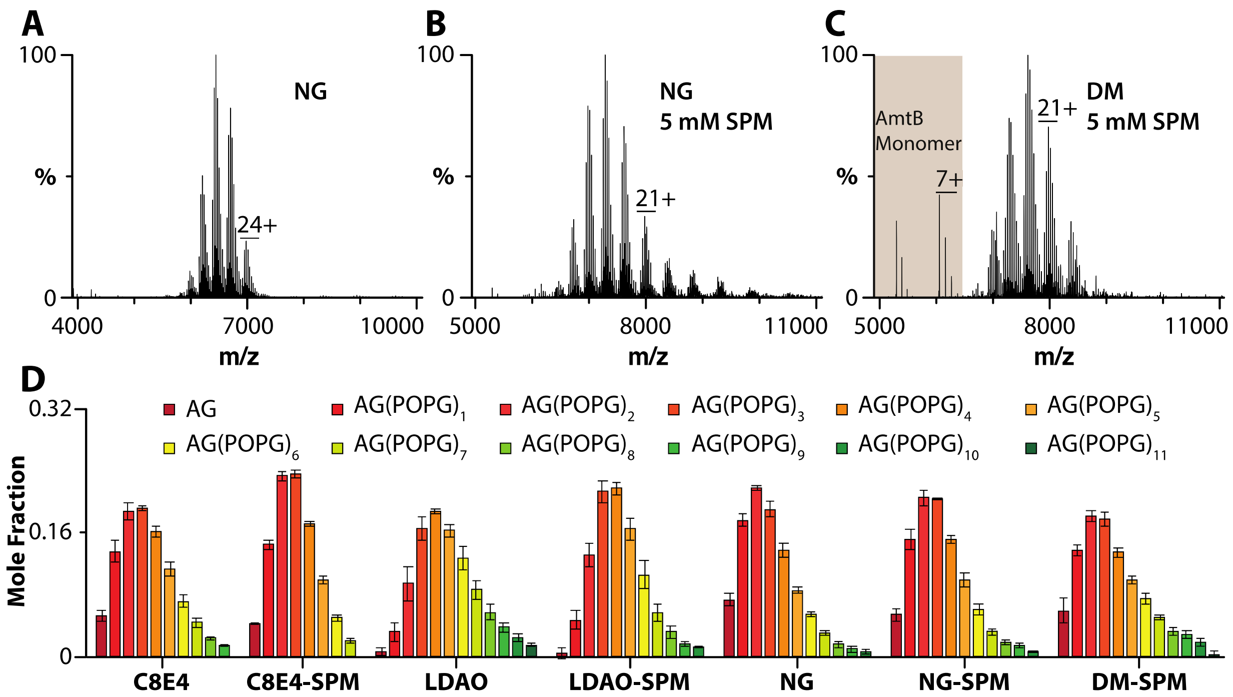


**Figure S8. POPG binding to AmtB-GlnK in different detergents.** A-C) AmtB-GlnK (2 μM) mixed with 50 μM POPG. Shown as described in Figure 4. D) Plot of the mole fraction for different species determined from the deconvolution of the mass spectra in different environments. Reported are the mean and standard deviation (*n*=3).

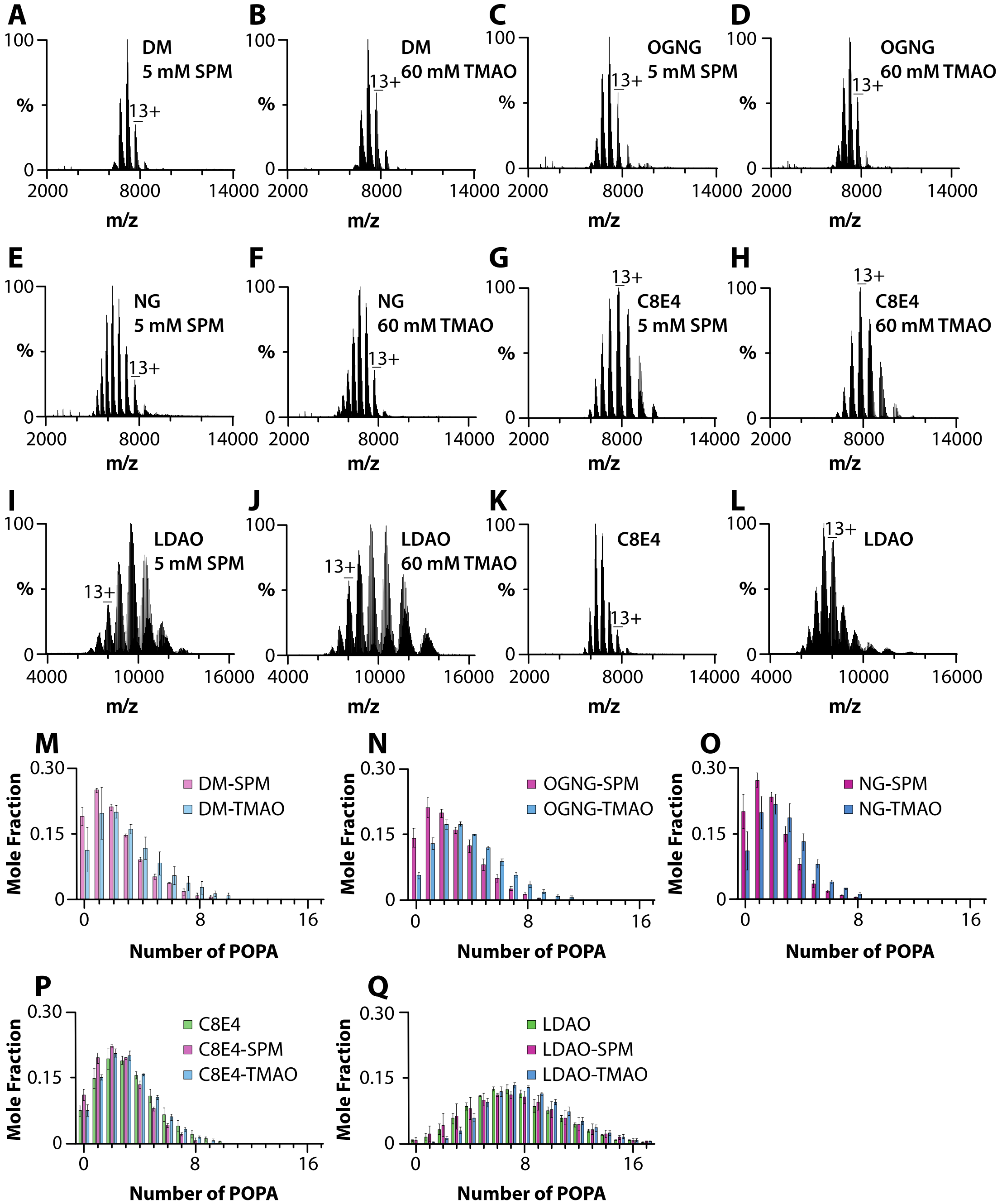


**Figure S9. POPA binding to AqpZ in different detergents.** A-L) AqpZ (1 µM) mixed with 50 µM POPA. Shown as described in Figure S6. M-Q) Plot of the mole fraction for different species determined from the deconvolution of the mass spectra shown in A-L. Reported are the mean and standard deviation (*n*=3).

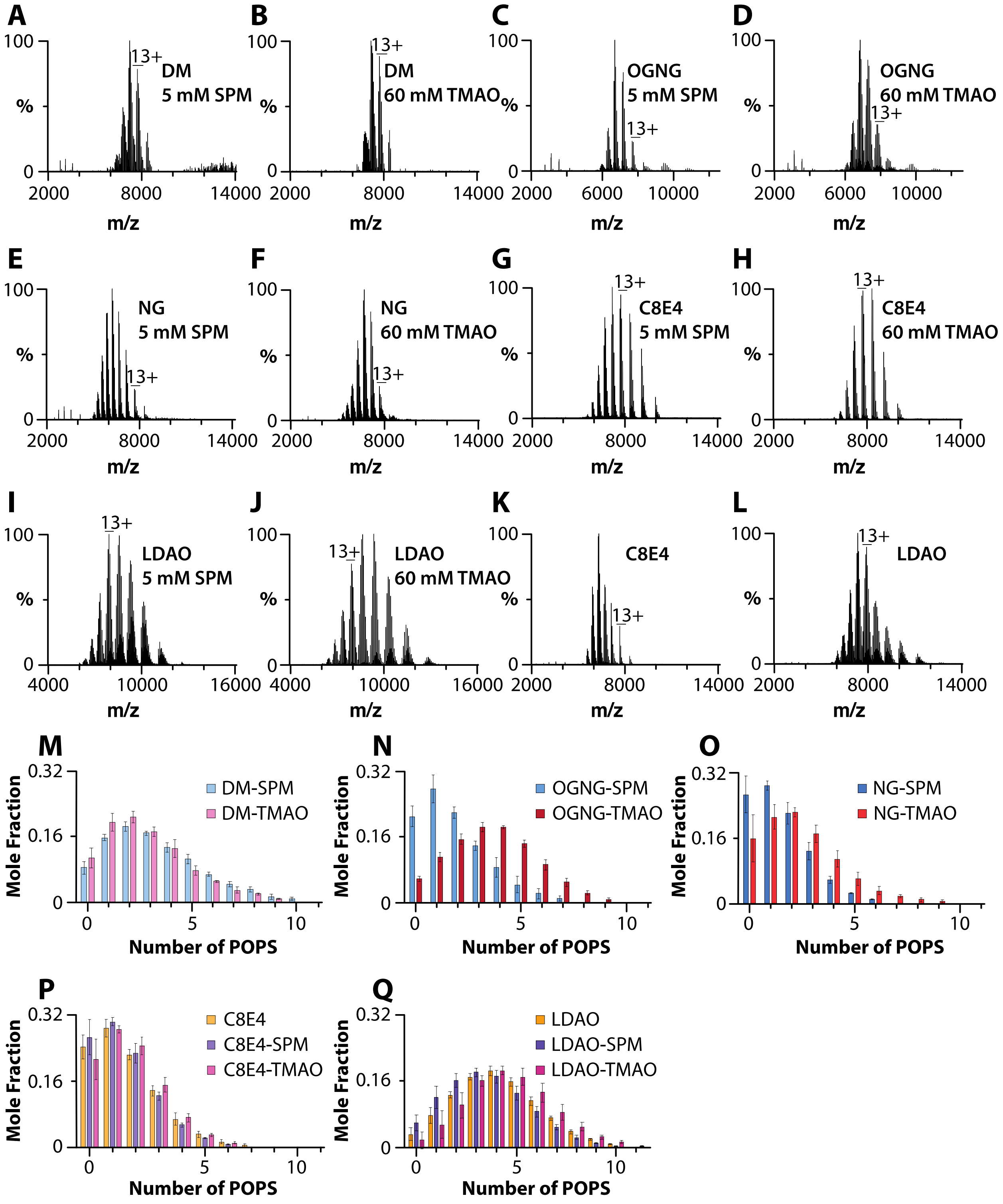


**Figure S10. POPS binding to AqpZ in different detergents.** A-L) AqpZ (1 µM) was mixed with 50 µM POPS. Shown as described in Figure S6. M-Q) Plot of the mole fraction for different species determined from the deconvolution of the mass spectra shown in A-L. Reported are the mean and standard deviation (*n*=3).

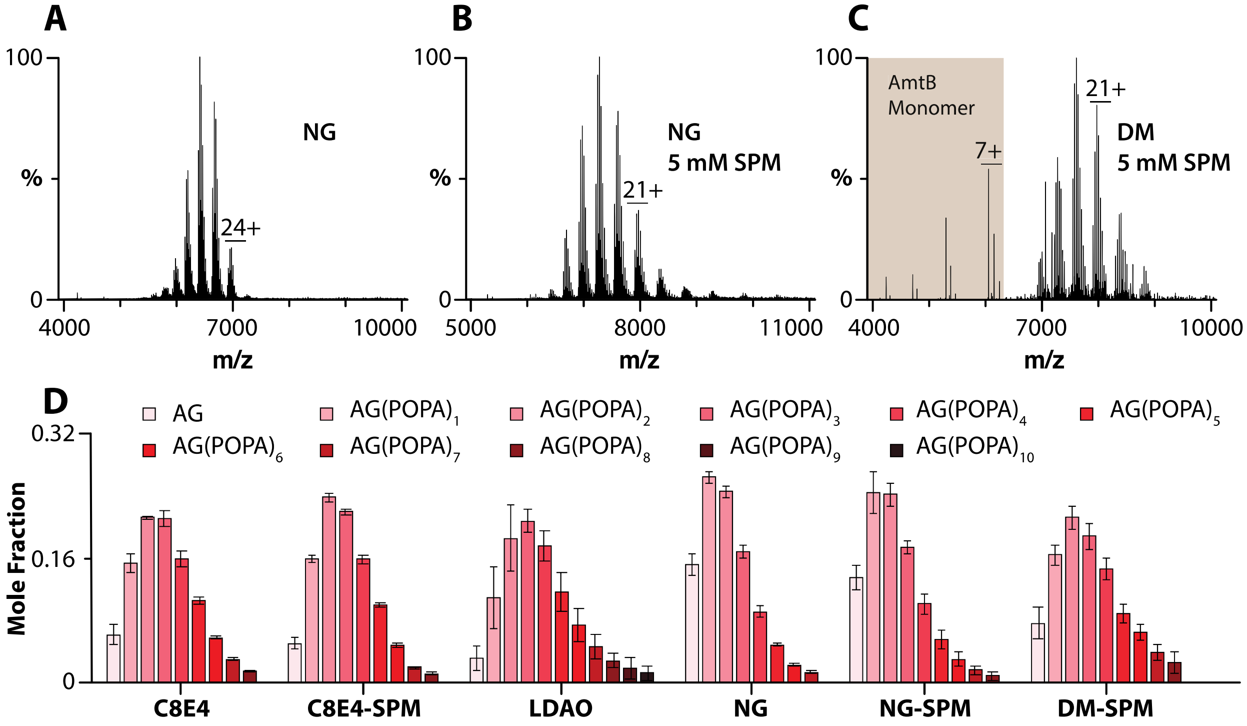


**Figure 11. AmtB-GlnK binding POPA in different detergents.** A-C) AmtB-GlnK (2 µM) mixed with 50 µM POPA. Shown as described in Figure 4. D) Plot of the mole fraction for different species determined from the deconvolution of the mass spectra shown in different environments. Reported are the mean and standard deviation (*n*=3).

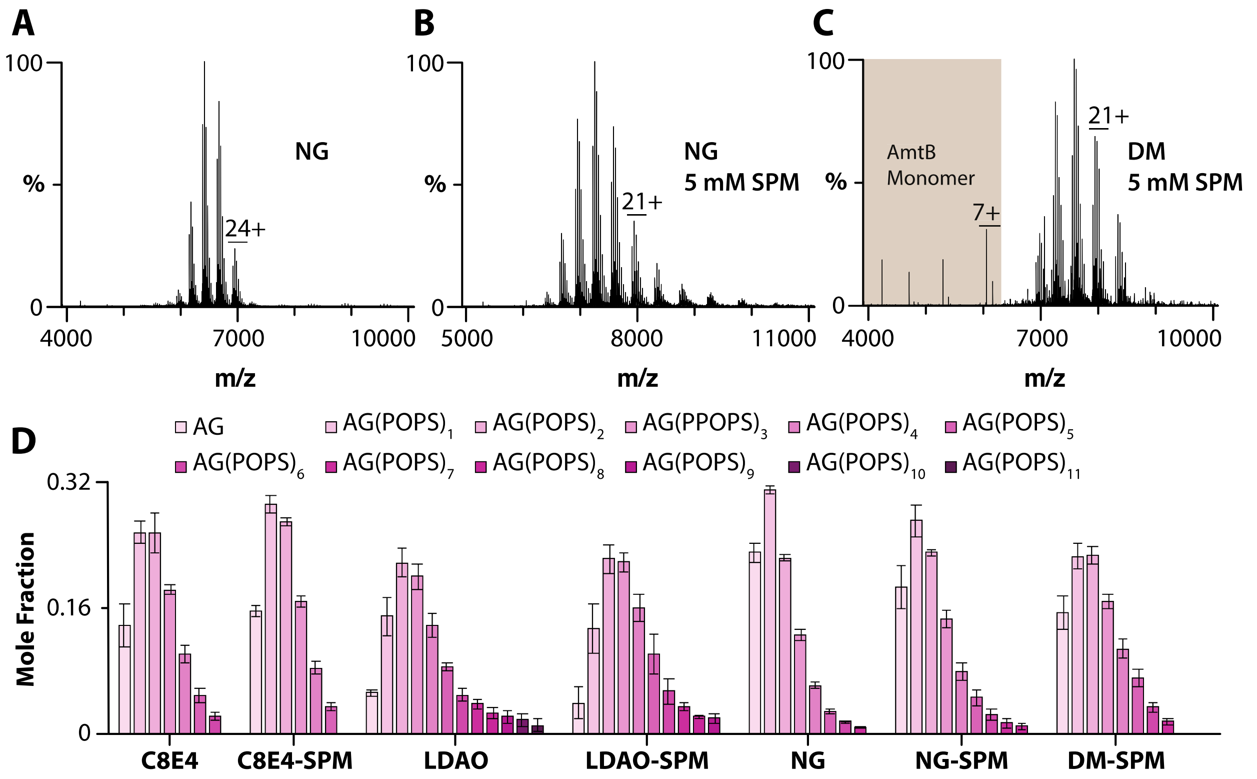


**Figure 12. POPS binding to AmtB-GlnK in different detergents.** A-C) AmtB-GlnK (2 μM) mixed with 50 μM POPS. Shown as described in Figure 4. D) Plot of the mole fraction for different species determined from the deconvolution of the mass spectra shown in A-C. Reported are the mean and standard deviation (*n*=3).

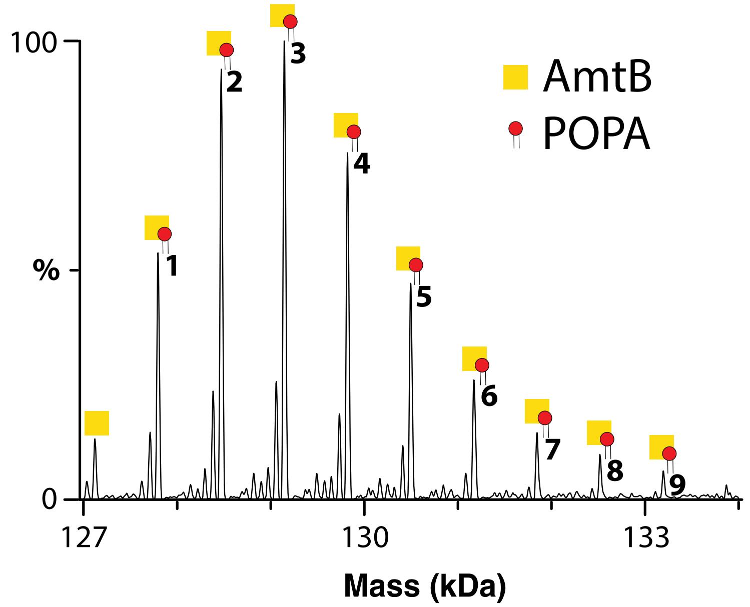


**Figure S13. POPA binding to AmtB in LDAO with spermine.** Deconvoluted mass spectra of 2 µM AmtB-GlnK in LDAO and 5 mM SPM mixed with 50 µM POPA.

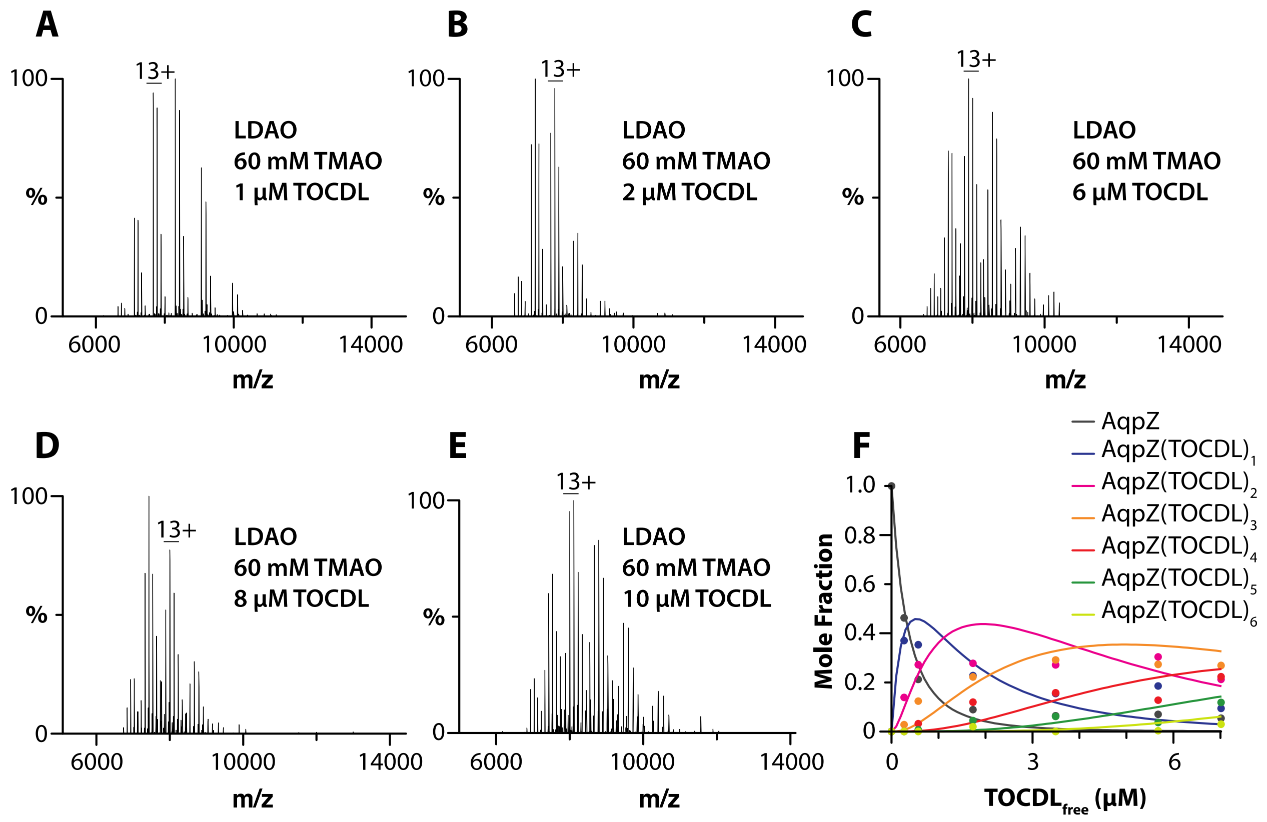


**Figure S14. Determination of AqpZ-TOCDL equilibrium binding constants.** A-E) AqpZ (1 µM) in LDAO and 60 mM TMAO mixed with different concentrations of TOCDL. F) Plot of mole fraction data (dots) determined from a titration series of TOCDL and subsequent fit of a sequential lipid binding model (lines).

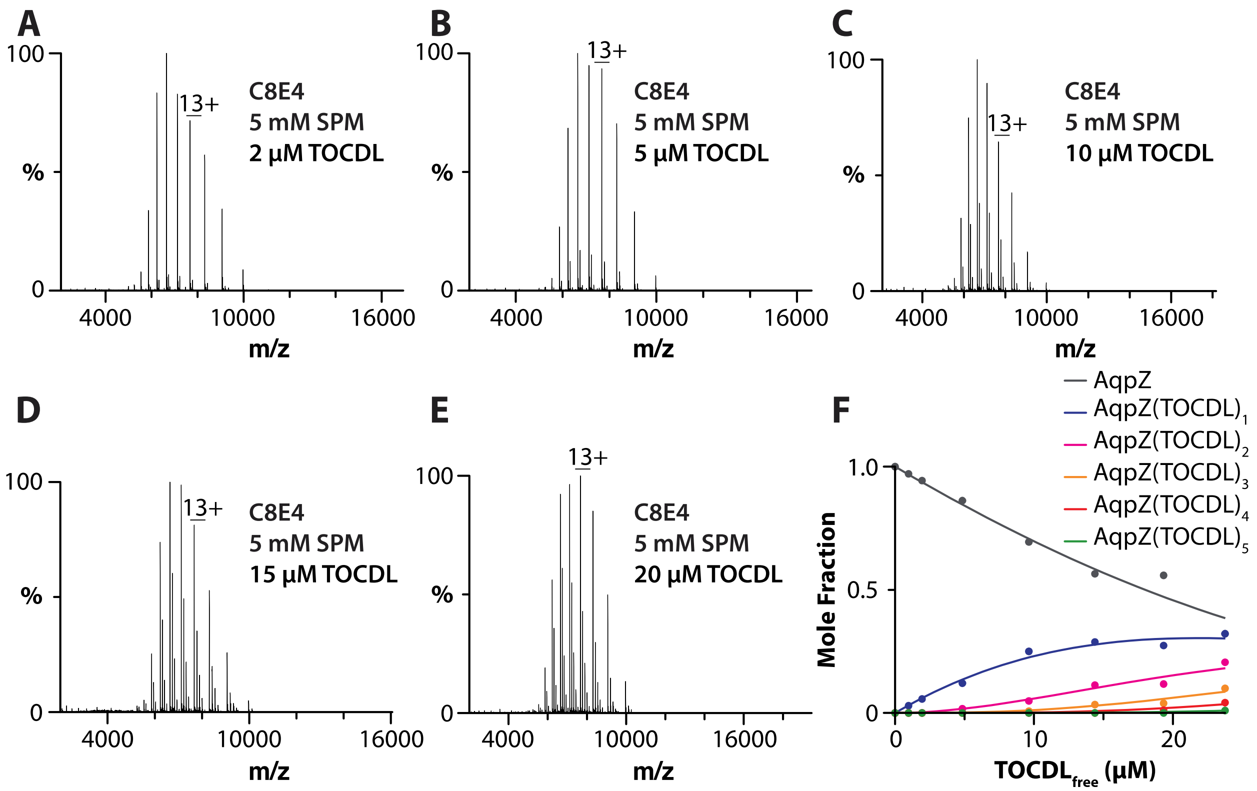


**Figure S15. Determination of AqpZ-TOCDL equilibrium binding constants.** A-E) AqpZ (1 μM) in C8E4 and 5 mM SPM mixed with different concentrations of TOCDL. F) Plot of mole fraction data (dots) determined from a titration series of TOCDL and subsequent fit of a sequential lipid binding model (lines).

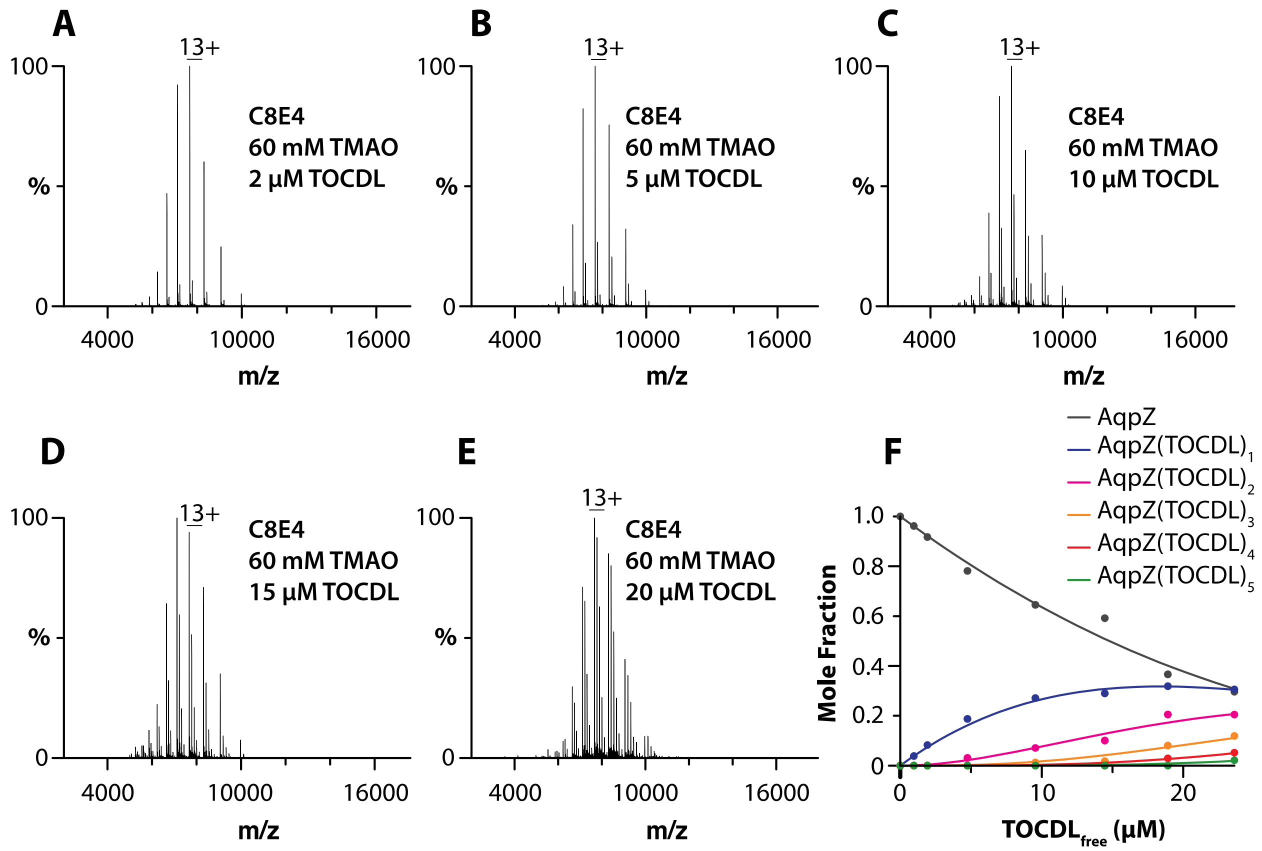


**Figure S16. Determination of AqpZ-TOCDL equilibrium binding constants.** A-E) AqpZ (1 µM) in C8E4 and 60 mM TMAO mixed with different concentrations of TOCDL. F) Plot of mole fraction data (dots) determined from a titration series of TOCDL and subsequent fit of a sequential lipid binding model (lines).

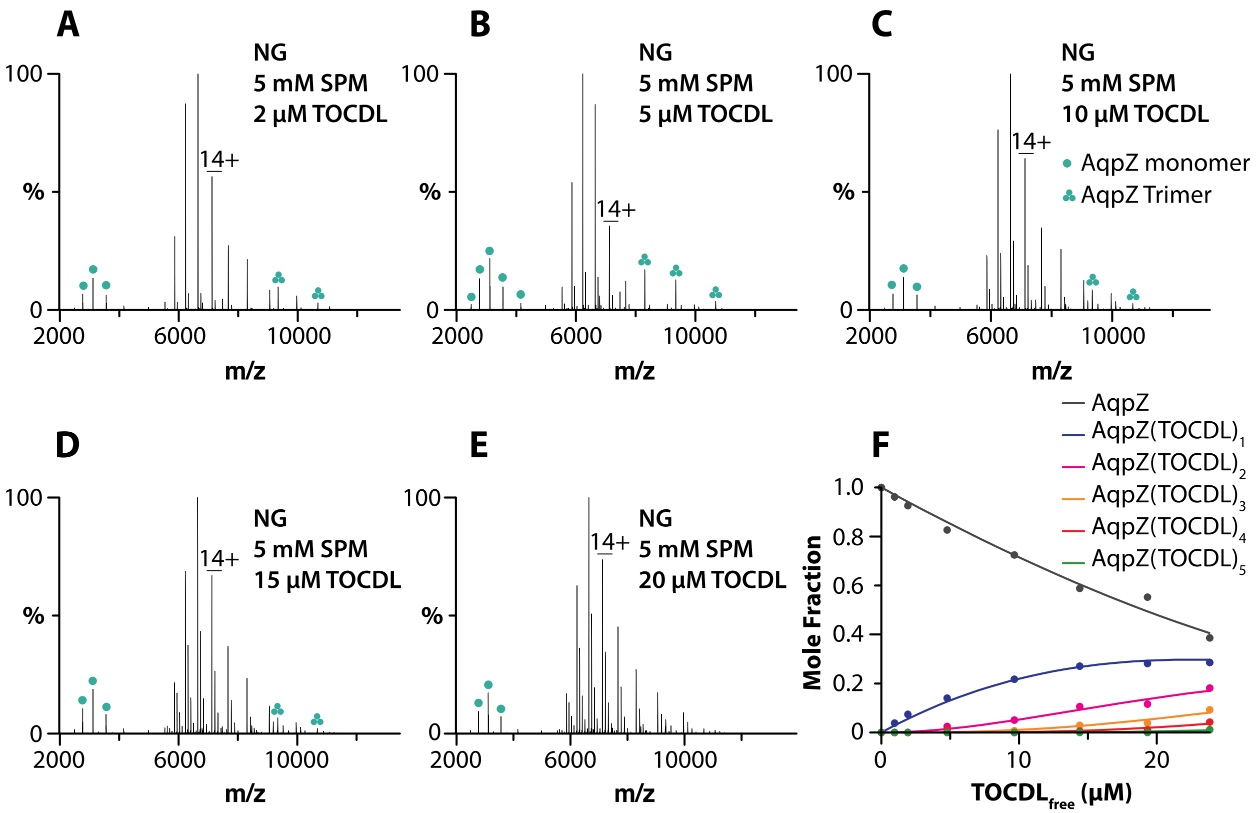


**Figure S17. Determination of AqpZ-TOCDL equilibrium binding constants.** A-E) AqpZ (1 µM) in NG and 5 mM SPM mixed with different concentrations of TOCDL. F) Plot of mole fraction data (dots) determined from a titration series of TOCDL and subsequent fit of a sequential lipid binding model (lines).

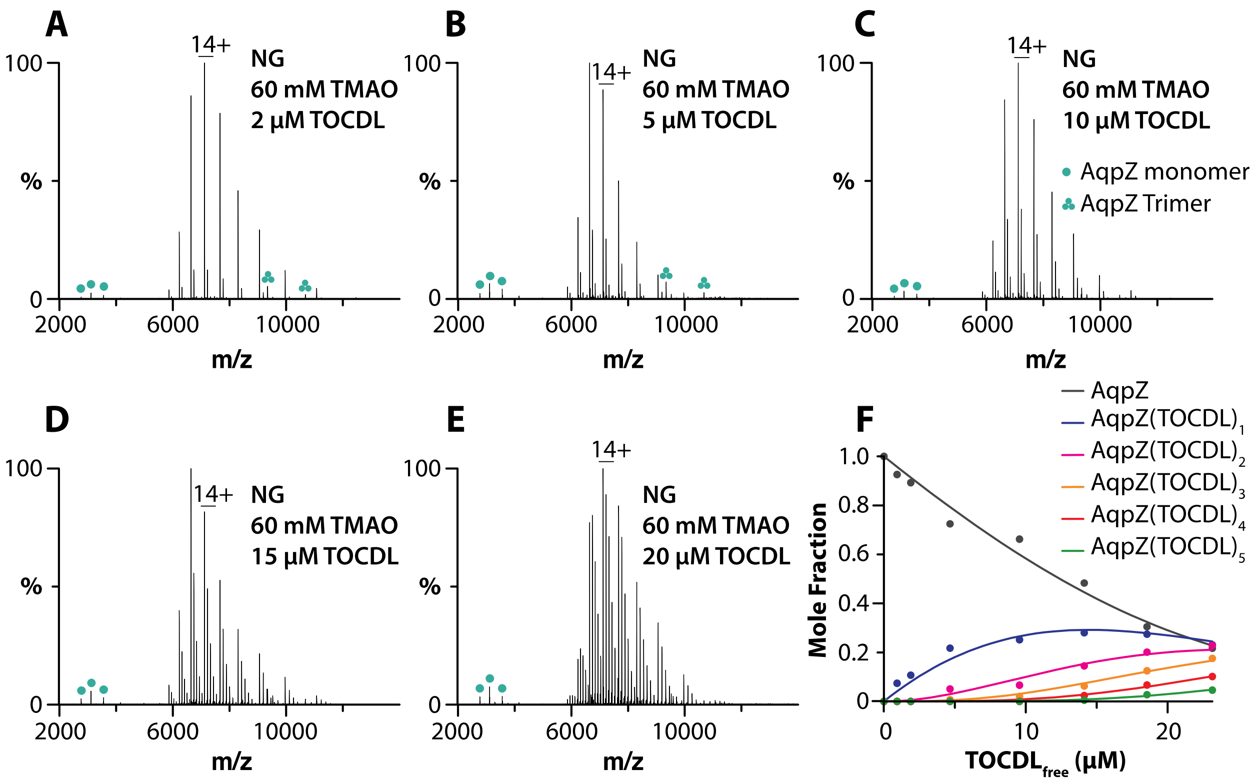


**Figure S18. Determination of AqpZ-TOCDL equilibrium binding constants.** A-E) AqpZ (1 µM) in NG and 60 mM TMAO mixed with different concentrations of TOCDL. F) Plot of mole fraction data (dots) determined from a titration series of TOCDL and subsequent fit of a sequential lipid binding model (lines).

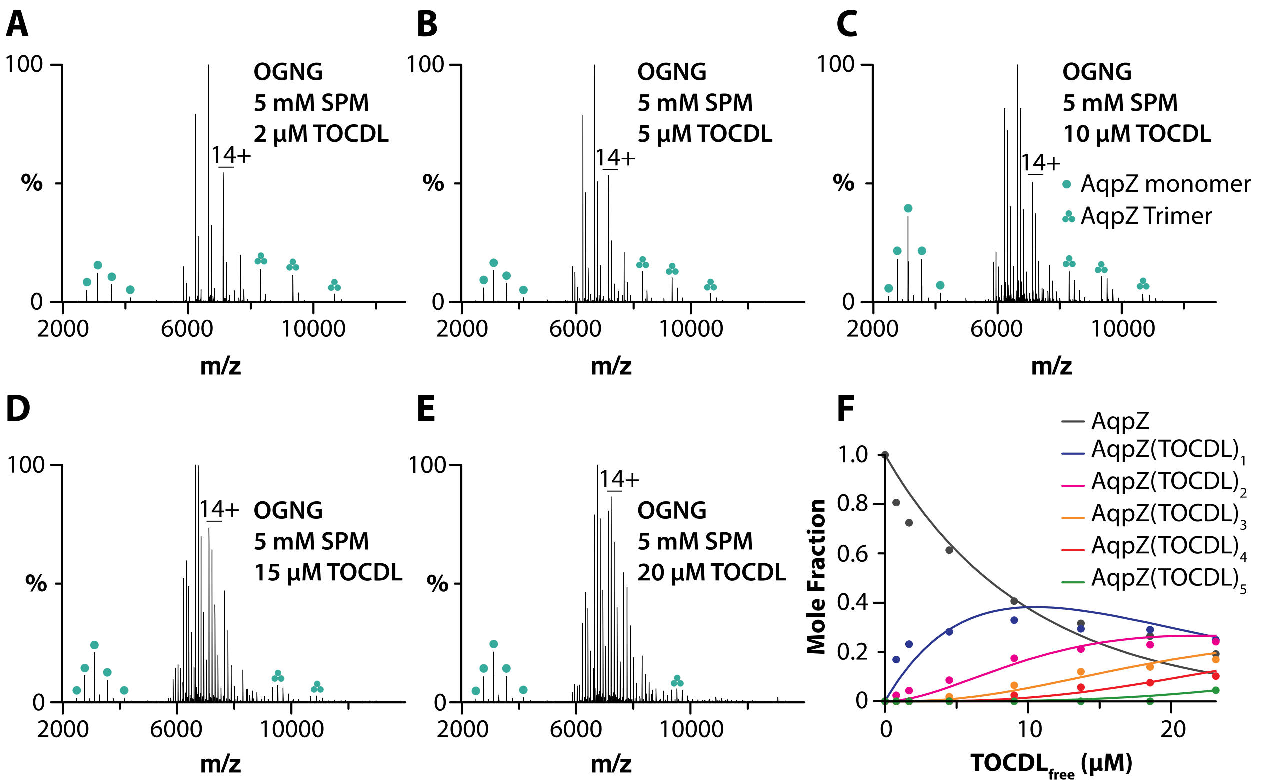


**Figure S19. Determination of AqpZ-TOCDL equilibrium binding constants.** A-E) AqpZ (1 µM) in OGNG and 5 mM SPM mixed with different concentrations of TOCDL. F) Plot of mole fraction data (dots) determined from a titration series of TOCDL and subsequent fit of a sequential lipid binding model (lines).

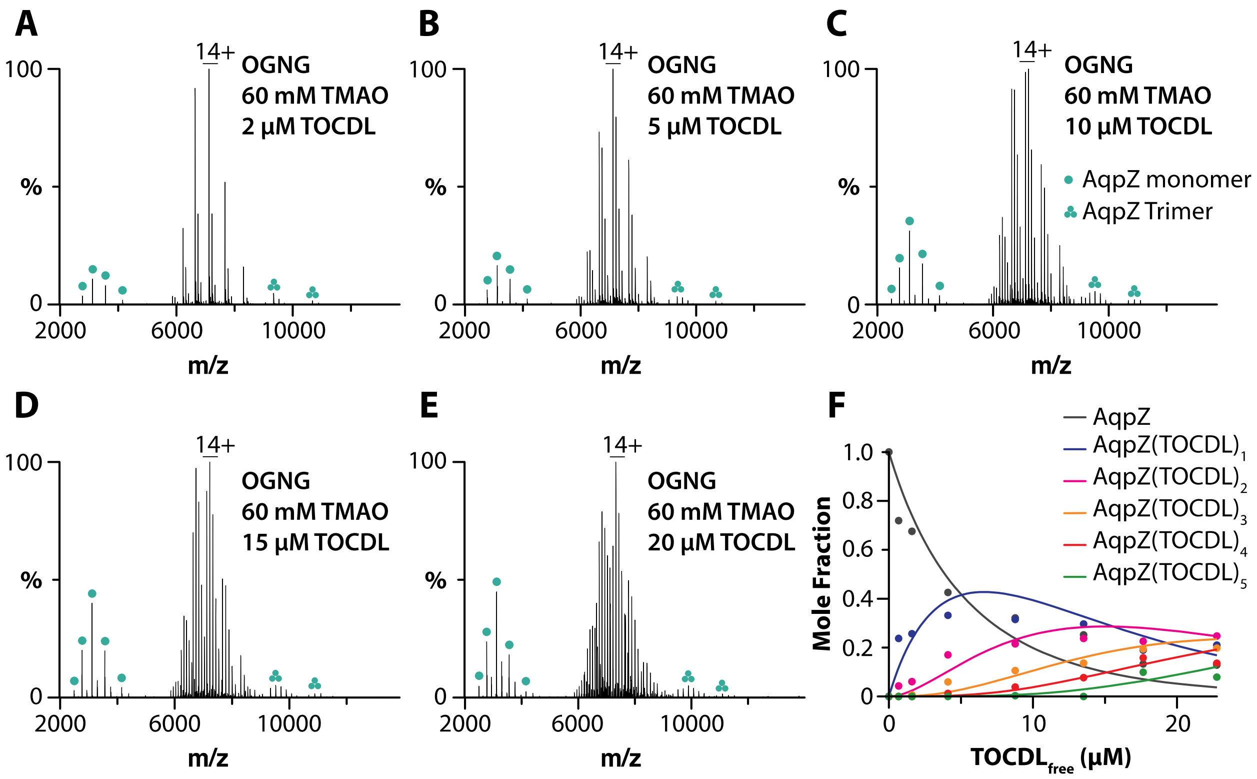


**Figure S20. Determination of AqpZ-TOCDL equilibrium binding constants.** A-E) AqpZ (1 μM) in OGNG and 60 mM TMAO mixed with different concentrations of TOCDL. F) Plot of mole fraction data (dots) determined from a titration series of TOCDL and subsequent fit of a sequential lipid binding model (lines).

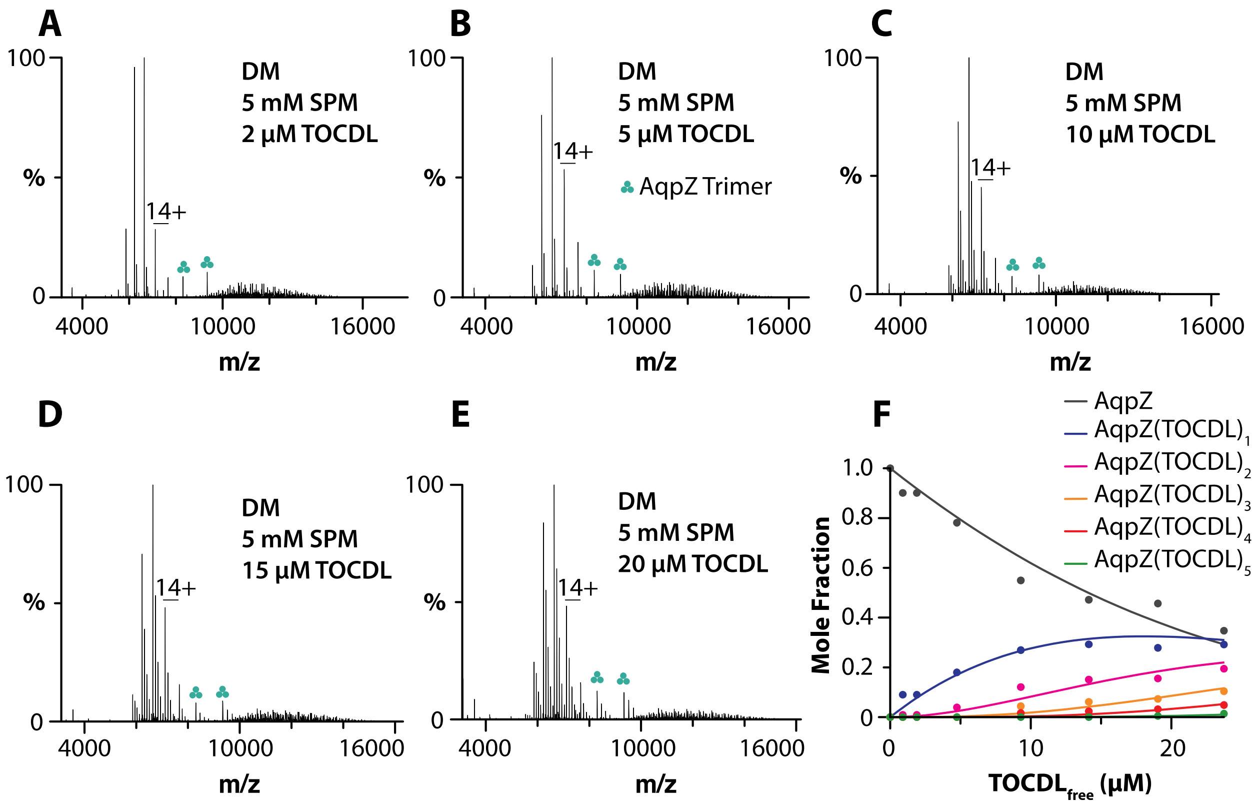


**Figure S21. Determination of AqpZ-TOCDL equilibrium binding constants.** A-E) AqpZ (1 µM) in DM and 5 mM SPM mixed with different concentrations of TOCDL. F) Plot of mole fraction data (dots) determined from a titration series of TOCDL and subsequent fit of a sequential lipid binding model (lines).

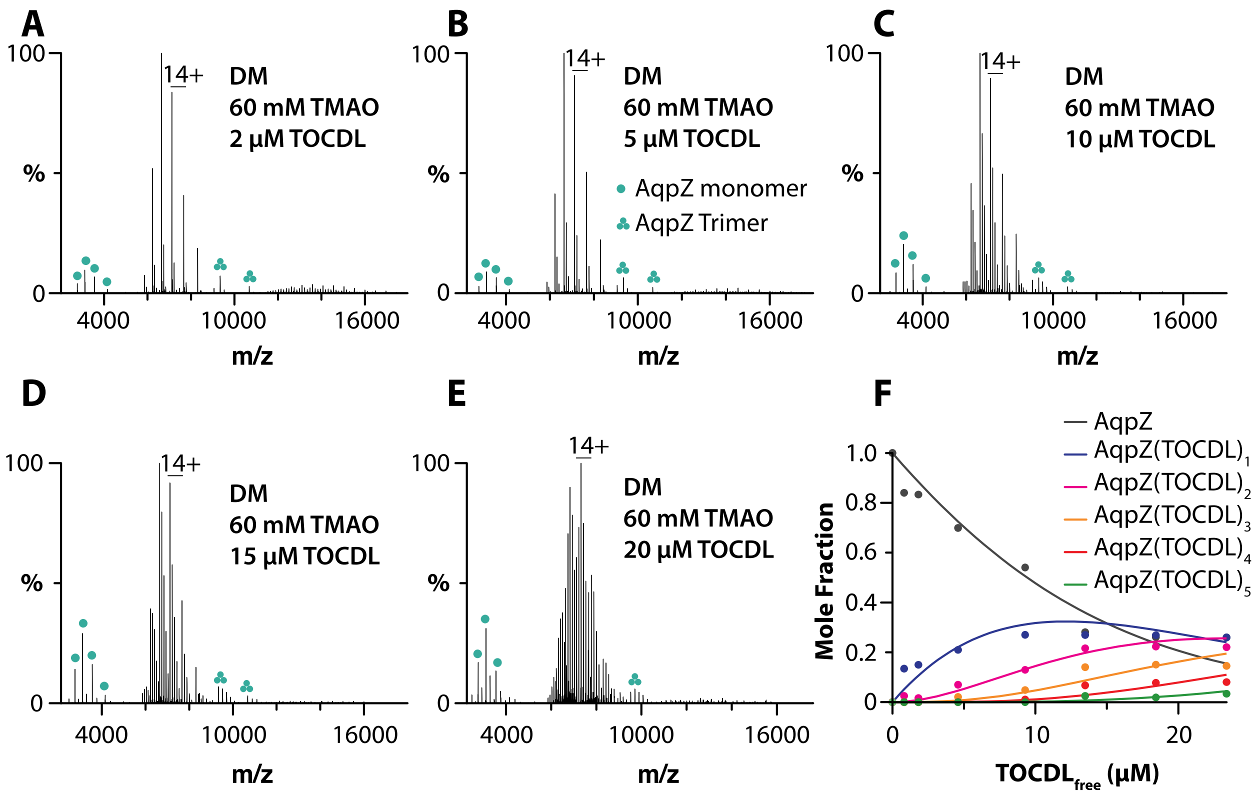


**Figure S22. Determination of AqpZ-TOCDL equilibrium binding constants.** A-E) AqpZ (1 μM) in DM and 60 mM TMAO mixed with different concentrations of TOCDL. F) Plot of mole fraction data (dots) determined from a titration series of TOCDL and subsequent fit of a sequential lipid binding model (lines).

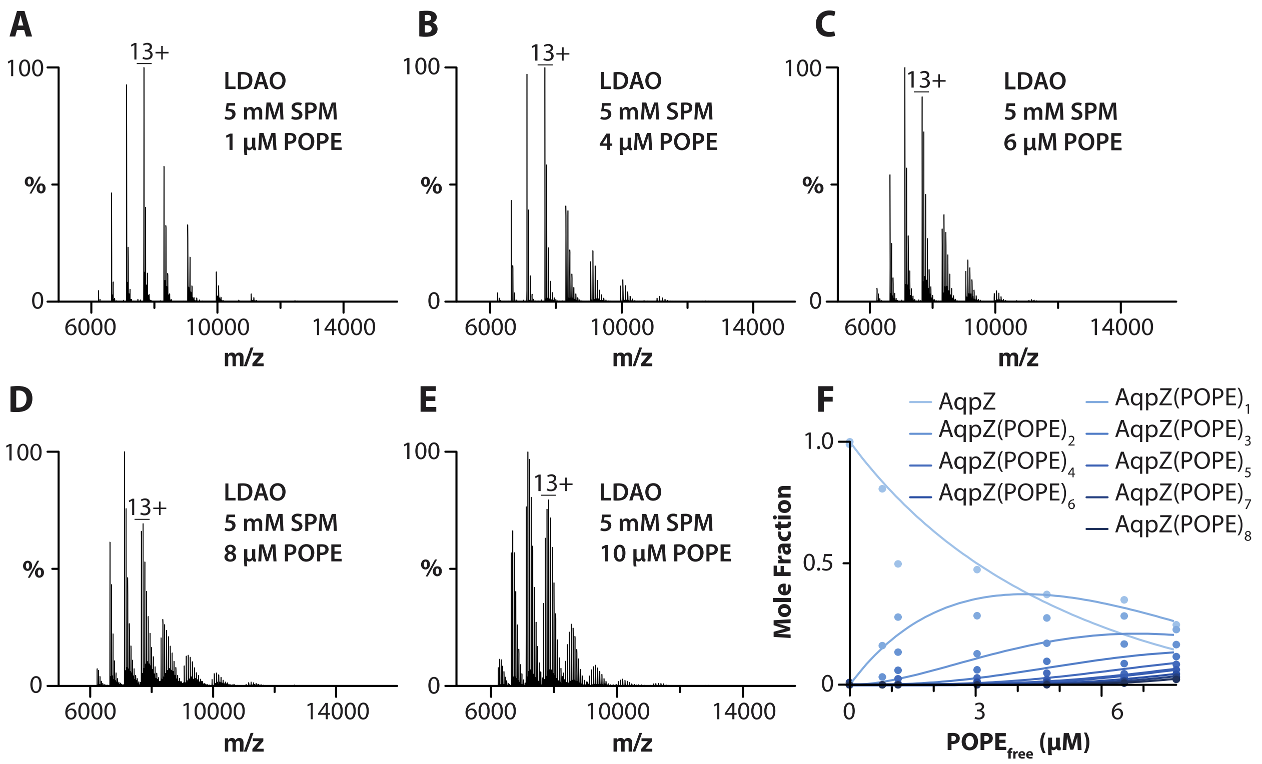


**Figure S23. Determination of AqpZ-POPE equilibrium binding constants.** A-E) AqpZ (1 µM) in LDAO and 5 mM SPM mixed with different concentrations of POPE. F) Plot of mole fraction data (dots) determined from a titration series of POPE and subsequent fit of a sequential lipid binding model (lines).

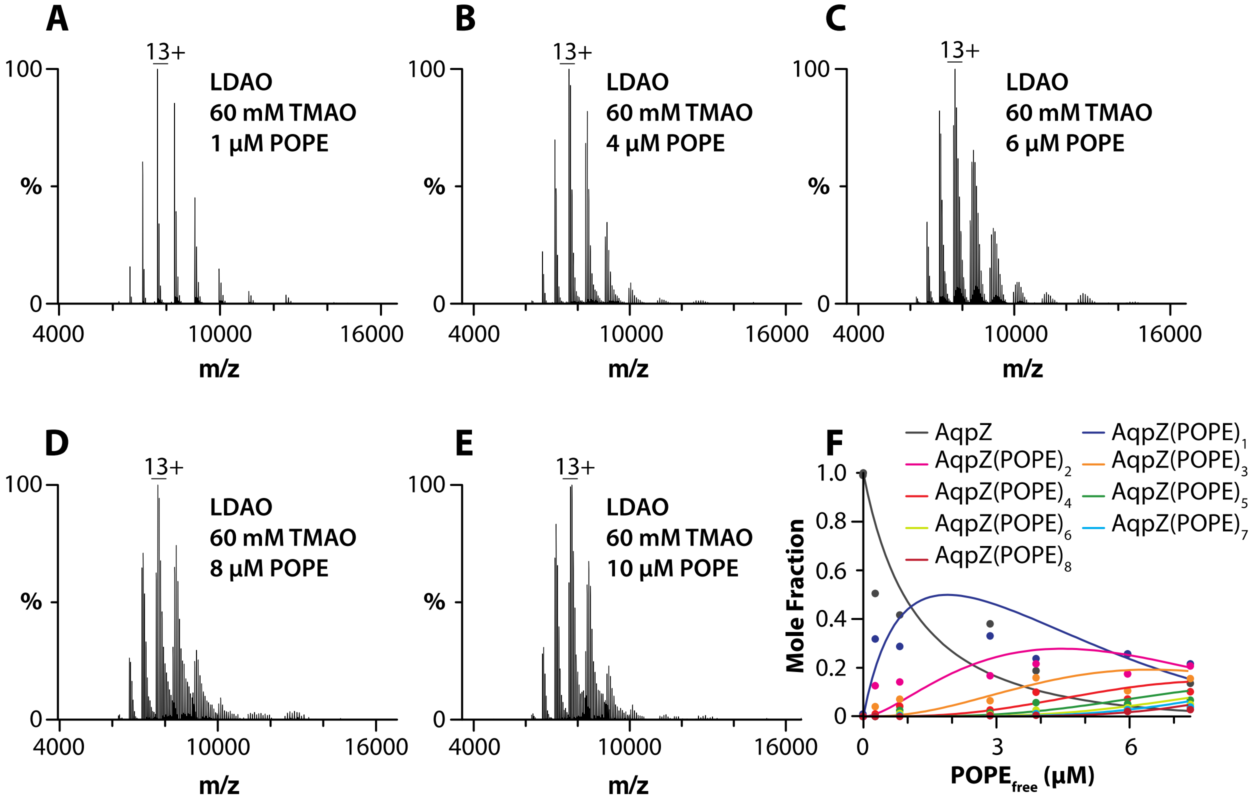


**Figure S24. Determination of AqpZ-POPE equilibrium binding constants.** A-E) AqpZ (1 μM) in LDAO and 60 mM TMAO mixed with different concentrations of POPE. F) Plot of mole fraction data (dots) determined from a titration series of POPE and subsequent fit of a sequential lipid binding model (lines).

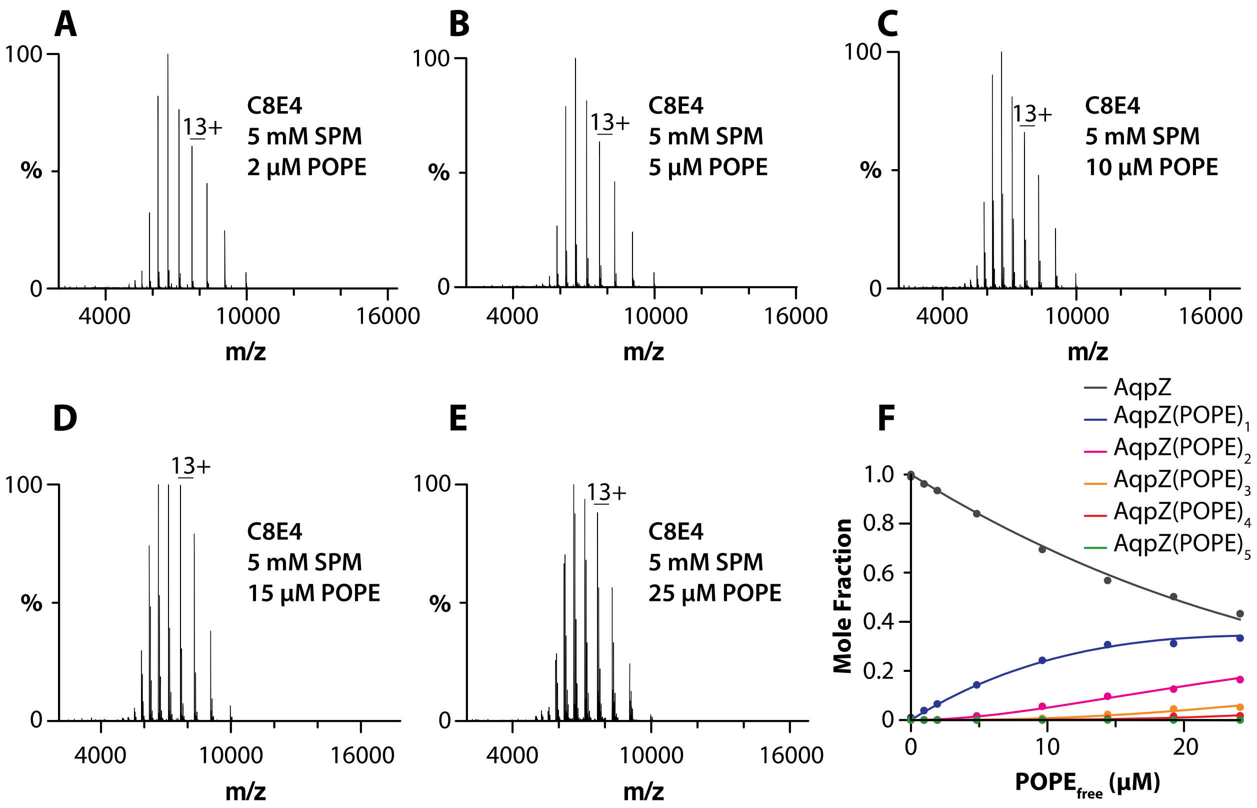


**Figure S25. Determination of AqpZ-POPE equilibrium binding constants.** A-E) AqpZ (1 μM) in C8E4 and 5 mM SPM mixed with different concentrations of POPE. F) Plot of mole fraction data (dots) determined from a titration series of POPE and subsequent fit of a sequential lipid binding model (lines).

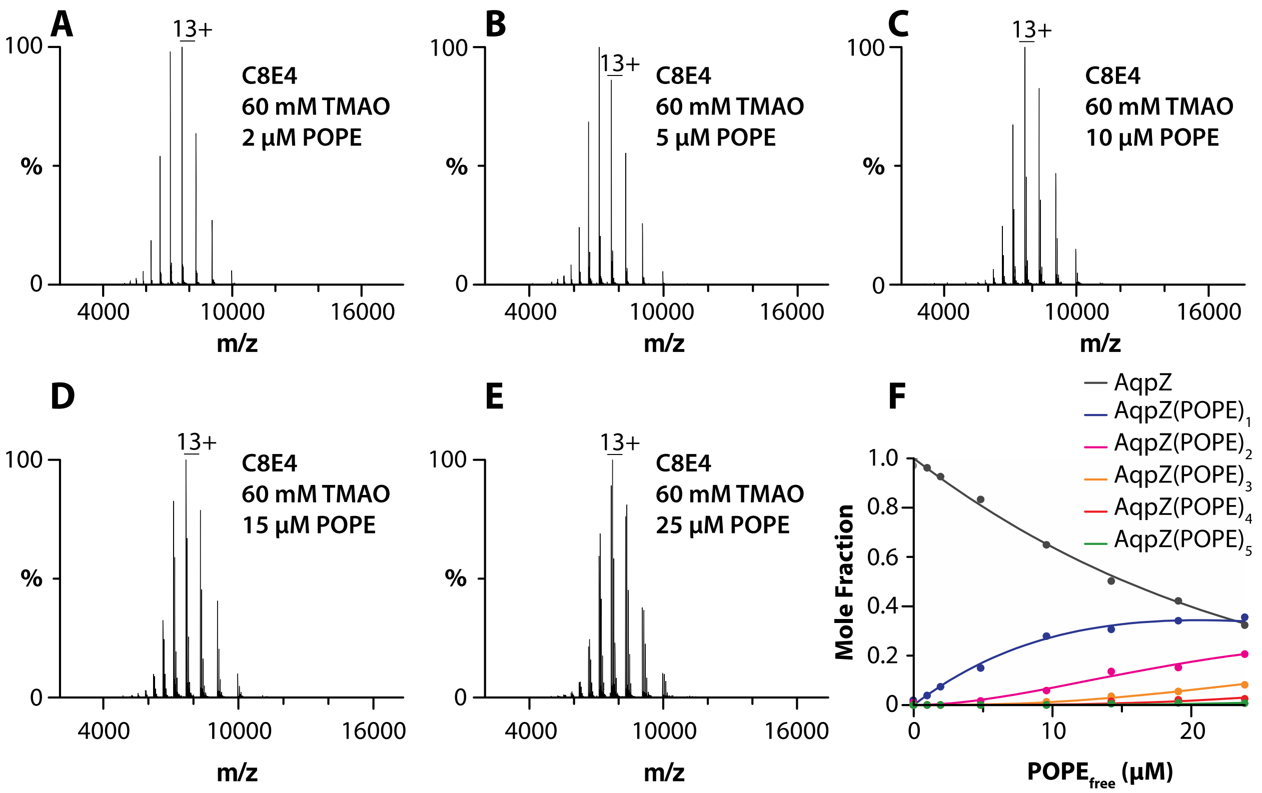


**Figure S26. Determination of AqpZ-POPE equilibrium binding constants.** A-E) AqpZ (1 μM) in C8E4 and 60 mM TMAO mixed with different concentrations of POPE. F) Plot of mole fraction data (dots) determined from a titration series of POPE and subsequent fit of a sequential lipid binding model (lines).

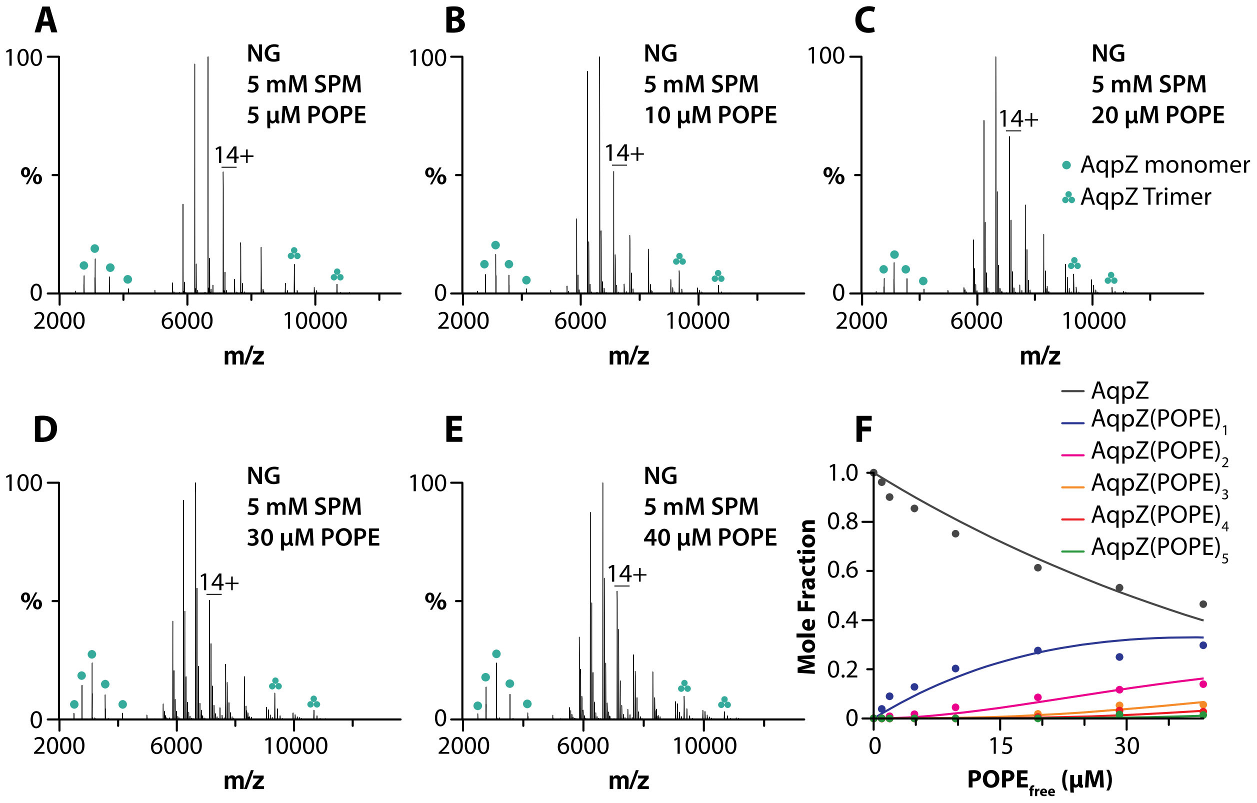


**Figure S27. Determination of AqpZ-POPE equilibrium binding constants.** A-E) AqpZ (1 µM) in NG and 5 mM SPM mixed with different concentrations of POPE. F) Plot of mole fraction data (dots) determined from a titration series of POPE and subsequent fit of a sequential lipid binding model (lines).

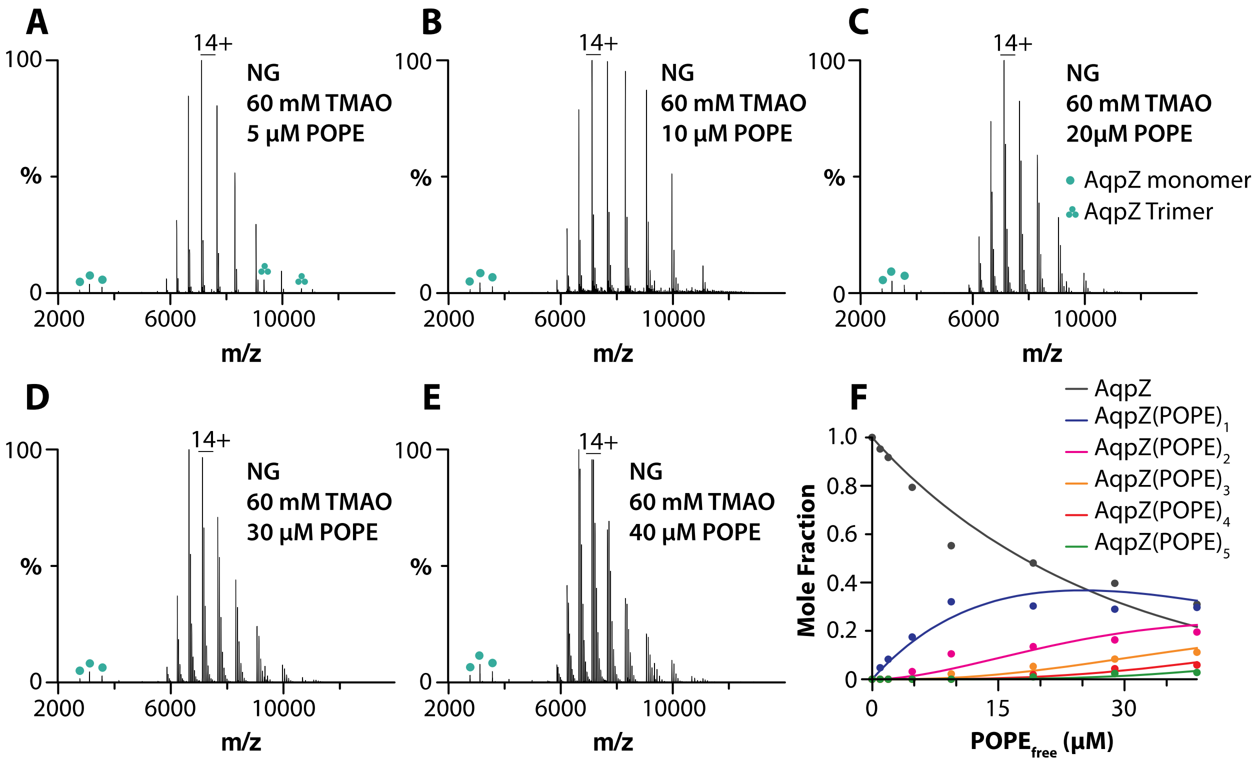


**Figure S28. Determination of AqpZ-POPE equilibrium binding constants.** A-E) AqpZ (1 µM) in NG and 60 mM TMAO mixed with different concentrations of POPE. F) Plot of mole fraction data (dots) determined from a titration series of POPE and subsequent fit of a sequential lipid binding model (lines).

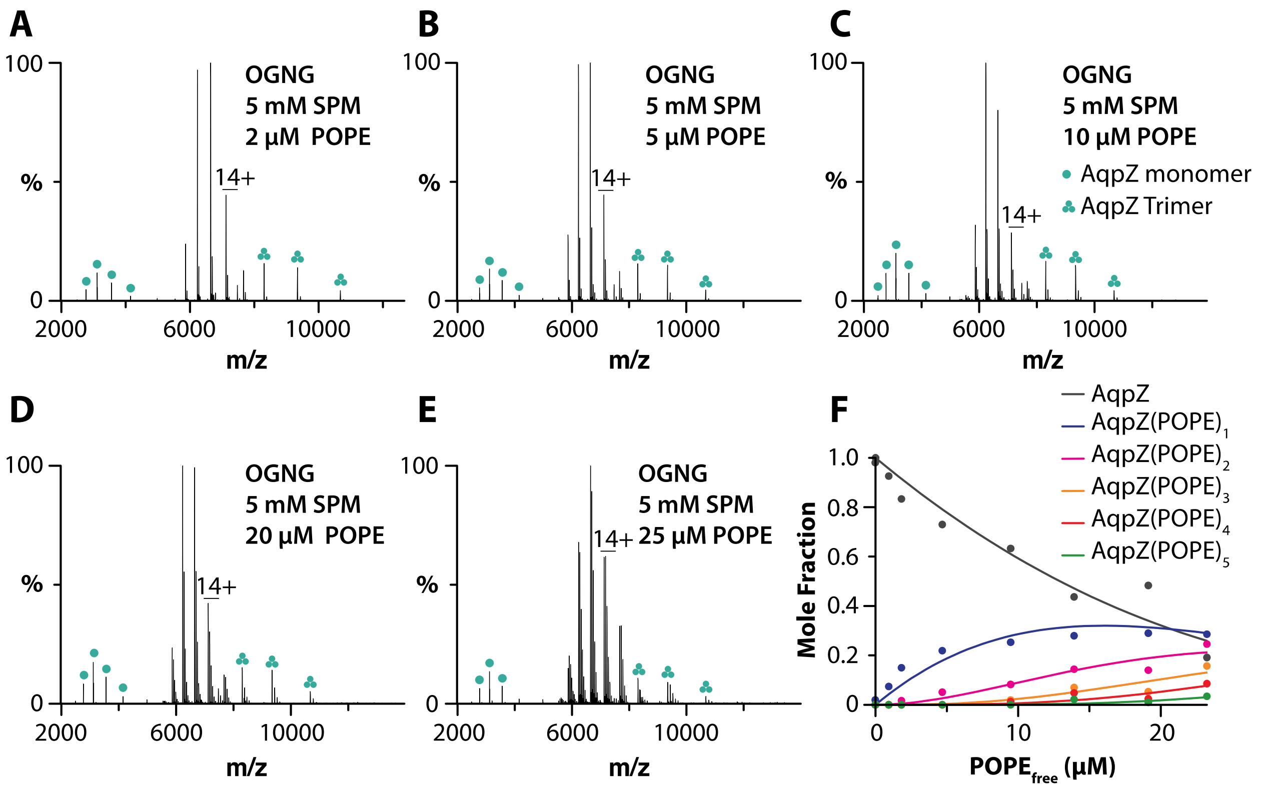


**Figure S29. Determination of AqpZ-POPE equilibrium binding constants.** A-E) AqpZ (1 µM) in OGNG and 5 mM SPM mixed with different concentrations of POPE. F) Plot of mole fraction data (dots) determined from a titration series of POPE and subsequent fit of a sequential lipid binding model (lines).

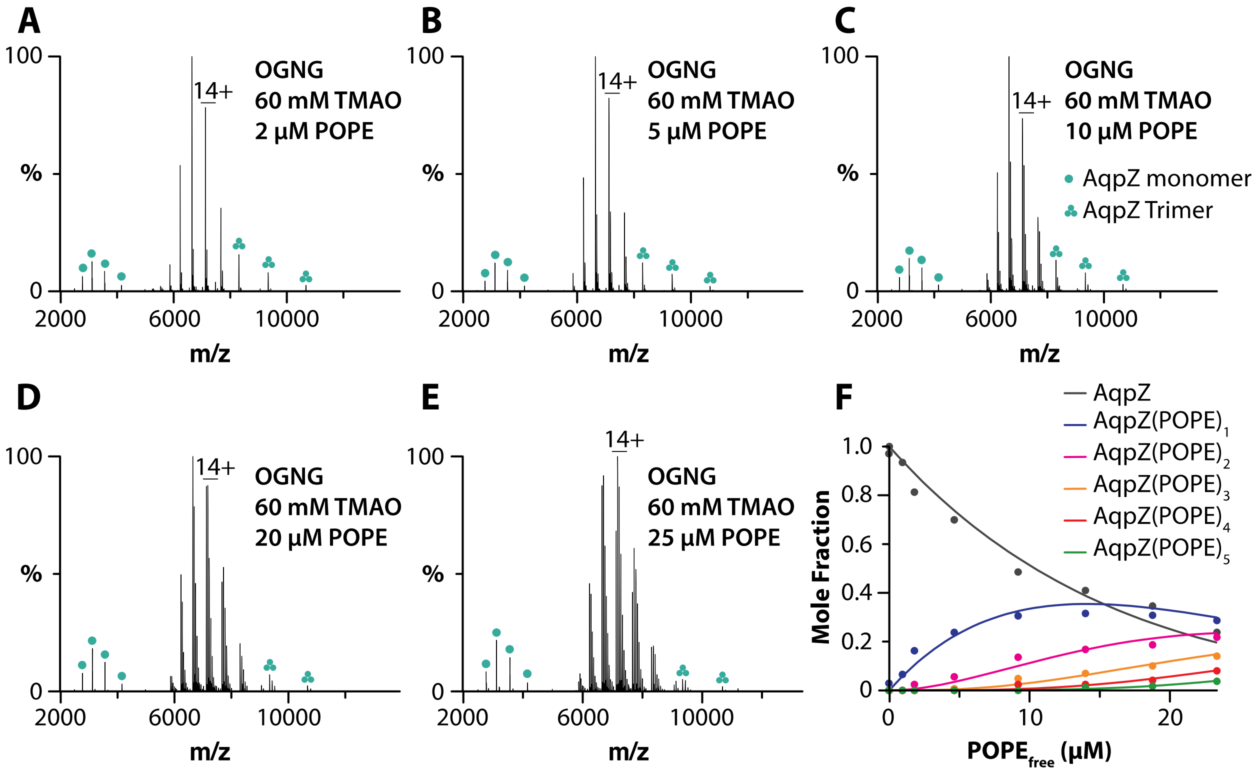


**Figure S30. Determination of AqpZ-POPE equilibrium binding constants.** A-E) AqpZ (1 µM) in OGNG and 60 mM TMAO mixed with different concentrations of POPE. F) Plot of mole fraction data (dots) determined from a titration series of POPE and subsequent fit of a sequential lipid binding model (lines).

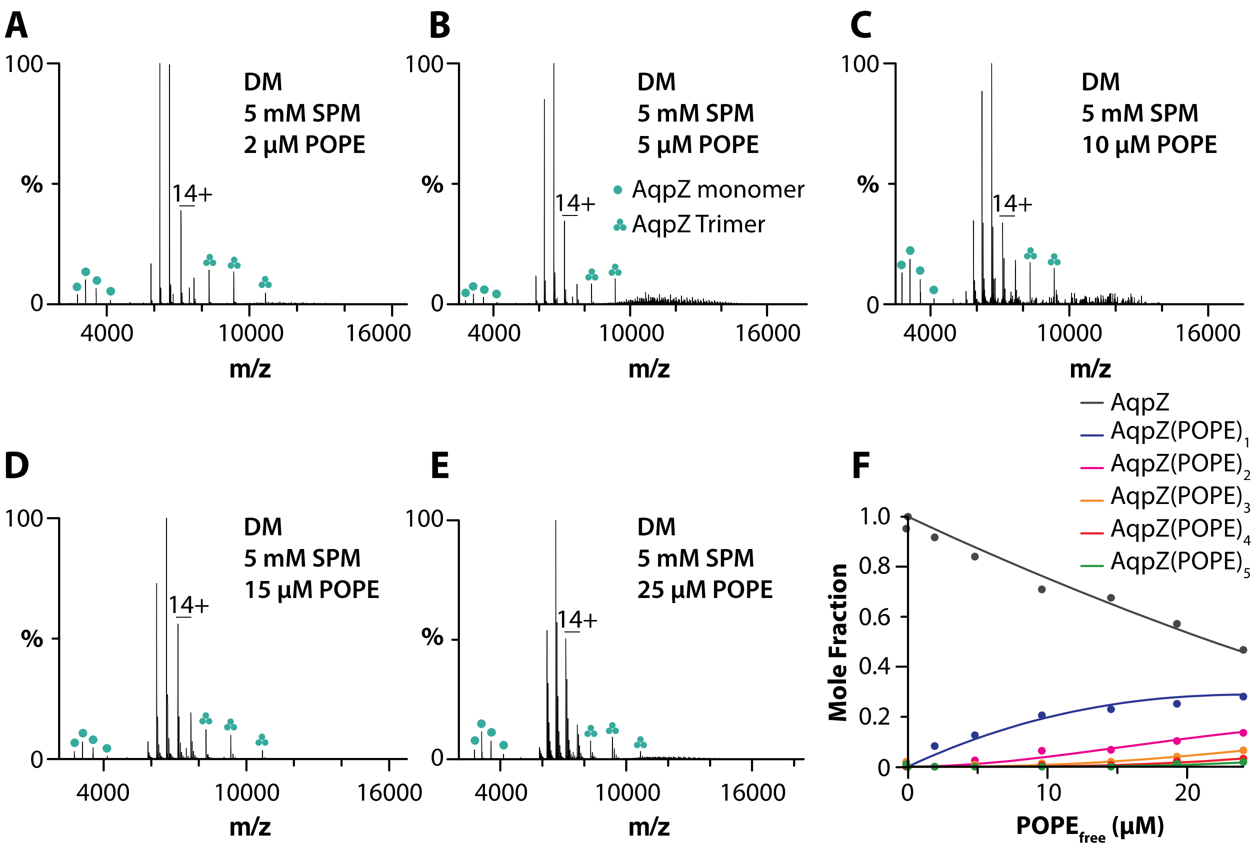


**Figure S31. Determination of AqpZ-POPE equilibrium binding constants.** A-E) AqpZ (1 µM) in DM and 5 mM SPM mixed with different concentrations of POPE. F) Plot of mole fraction data (dots) determined from a titration series of POPE and subsequent fit of a sequential lipid binding model (lines).

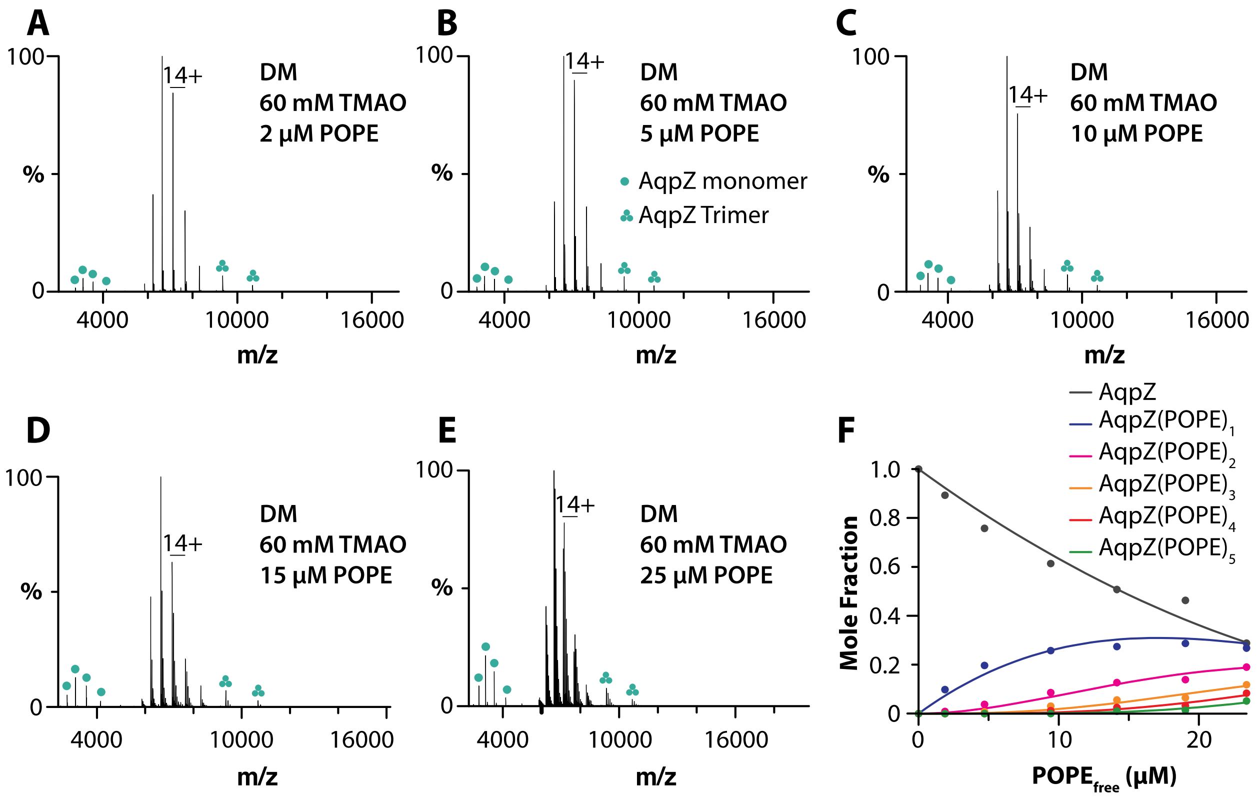


**Figure S32. Determination of AqpZ-POPE equilibrium binding constants.** A-E) AqpZ (1 µM) in DM and 60 mM TMAO mixed with different concentrations of POPE. F) Plot of mole fraction data (dots) determined from a titration series of POPE and subsequent fit of a sequential lipid binding model (lines).

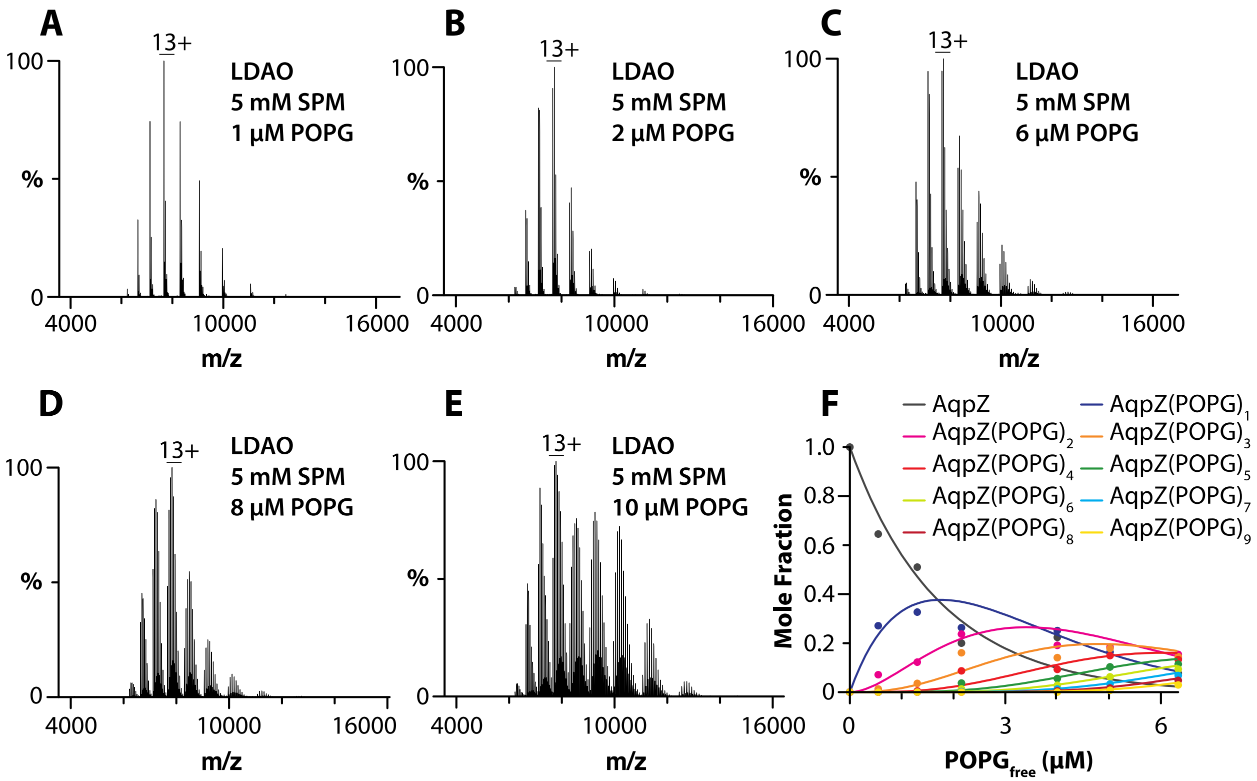


**Figure S33. Determination of AqpZ-POPG equilibrium binding constants.** A-E) AqpZ (1 μM) in LDAO and 5 mM SPM mixed with different concentrations of POPG. F) Plot of mole fraction data (dots) determined from a titration series of POPG and subsequent fit of a sequential lipid binding model (lines).

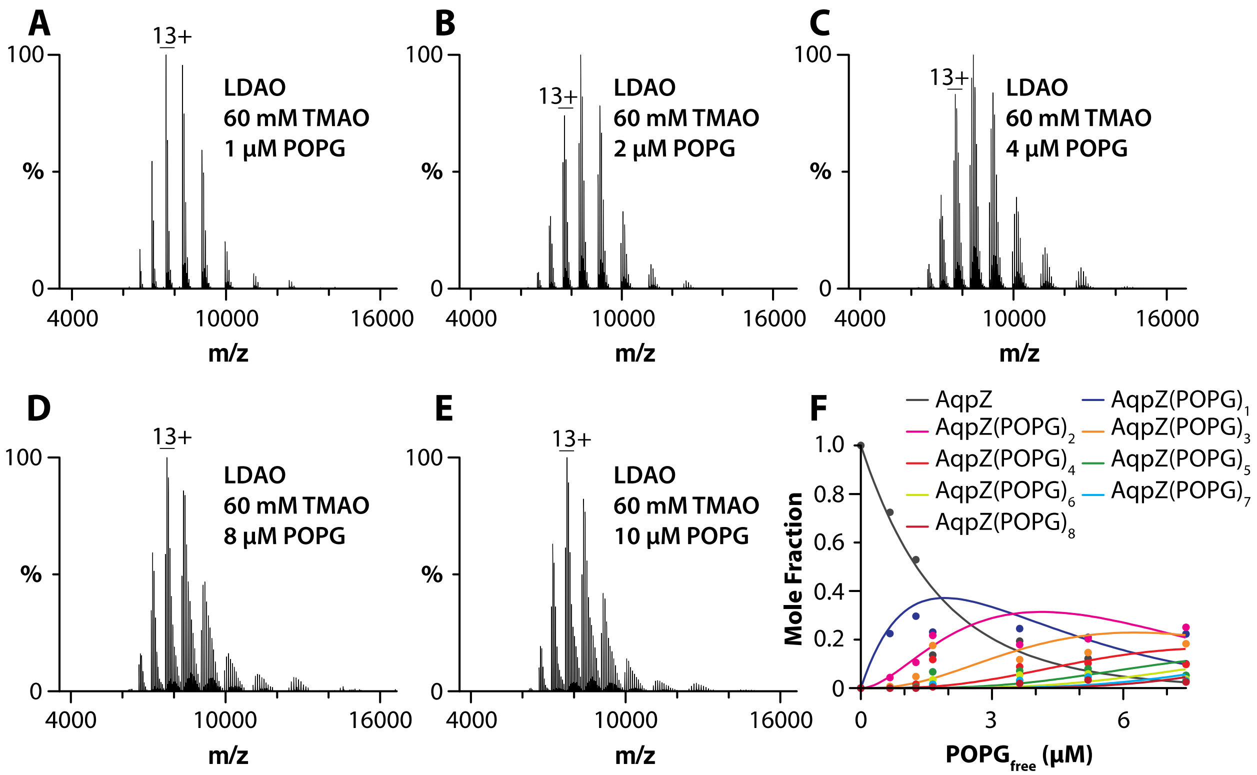


**Figure S34. Determination of AqpZ-POPG equilibrium binding constants.** A-E) AqpZ (1 μM) in LDAO and 60 mM TMAO mixed with different concentrations of POPG. F) Plot of mole fraction data (dots) determined from a titration series of POPG and subsequent fit of a sequential lipid binding model (lines).

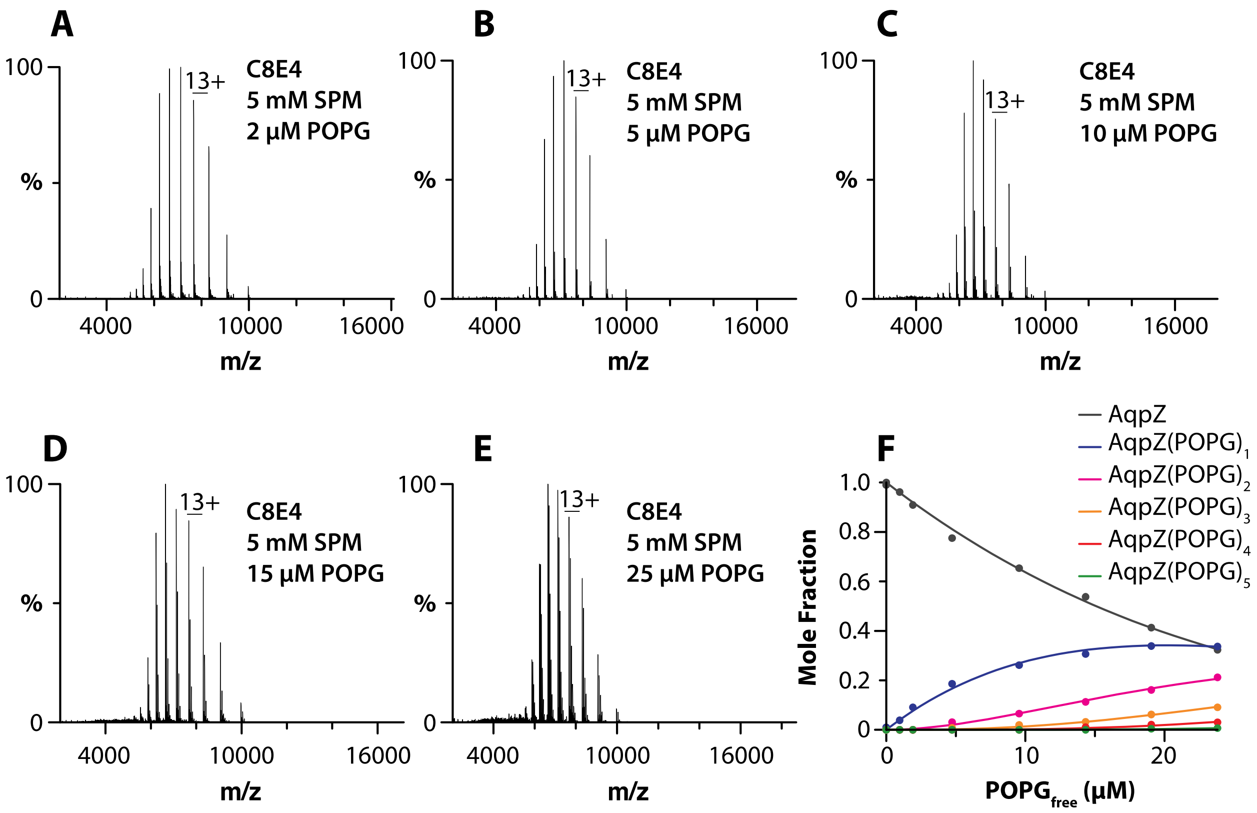


**Figure S35. Determination of AqpZ-POPG equilibrium binding constants.** A-E) AqpZ (1 μM) in C8E4 and 5 mM SPM mixed with different concentrations of POPG. F) Plot of mole fraction data (dots) determined from a titration series of POPG and subsequent fit of a sequential lipid binding model (lines).

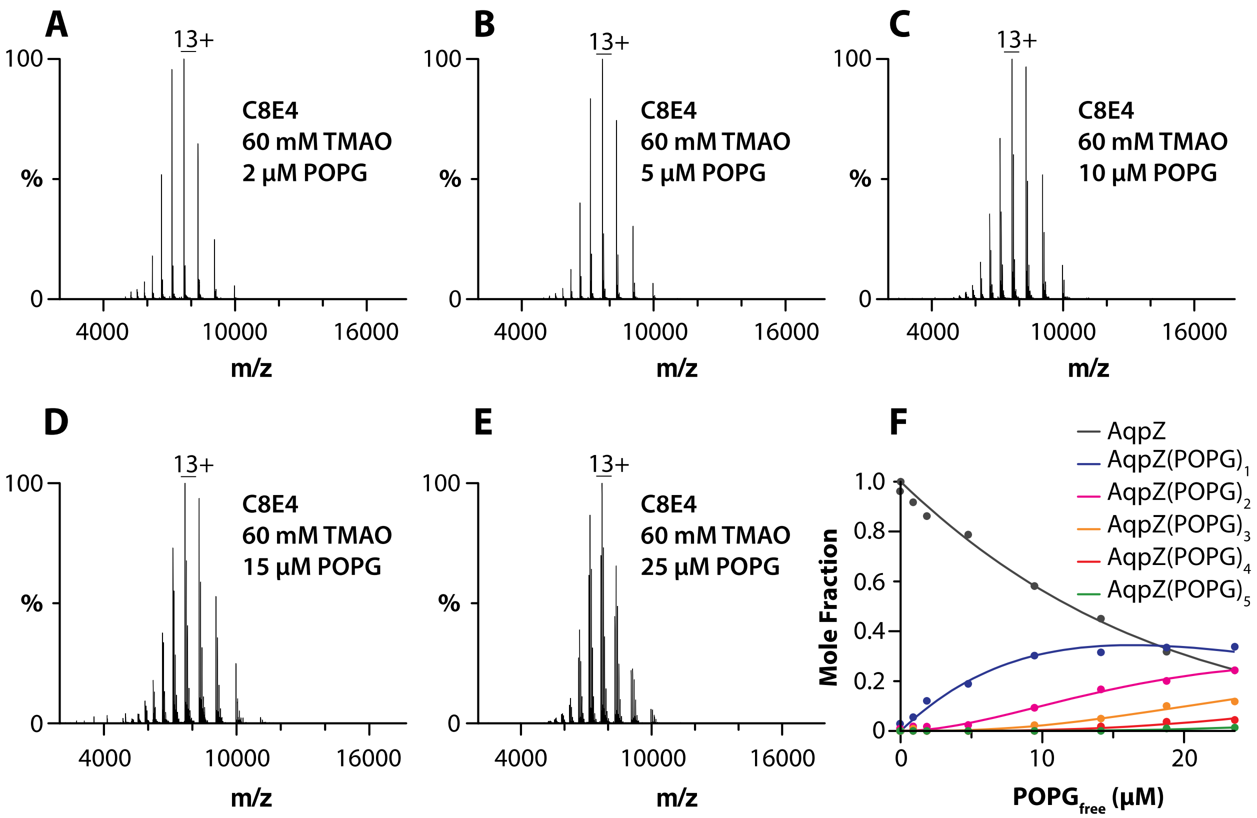


**Figure S36. Determination of AqpZ-POPG equilibrium binding constants.** A-E) AqpZ (1 µM) in C8E4 and 60 mM TMAO mixed with different concentrations of POPG. F) Plot of mole fraction data (dots) determined from a titration series of POPG and subsequent fit of a sequential lipid binding model (lines).

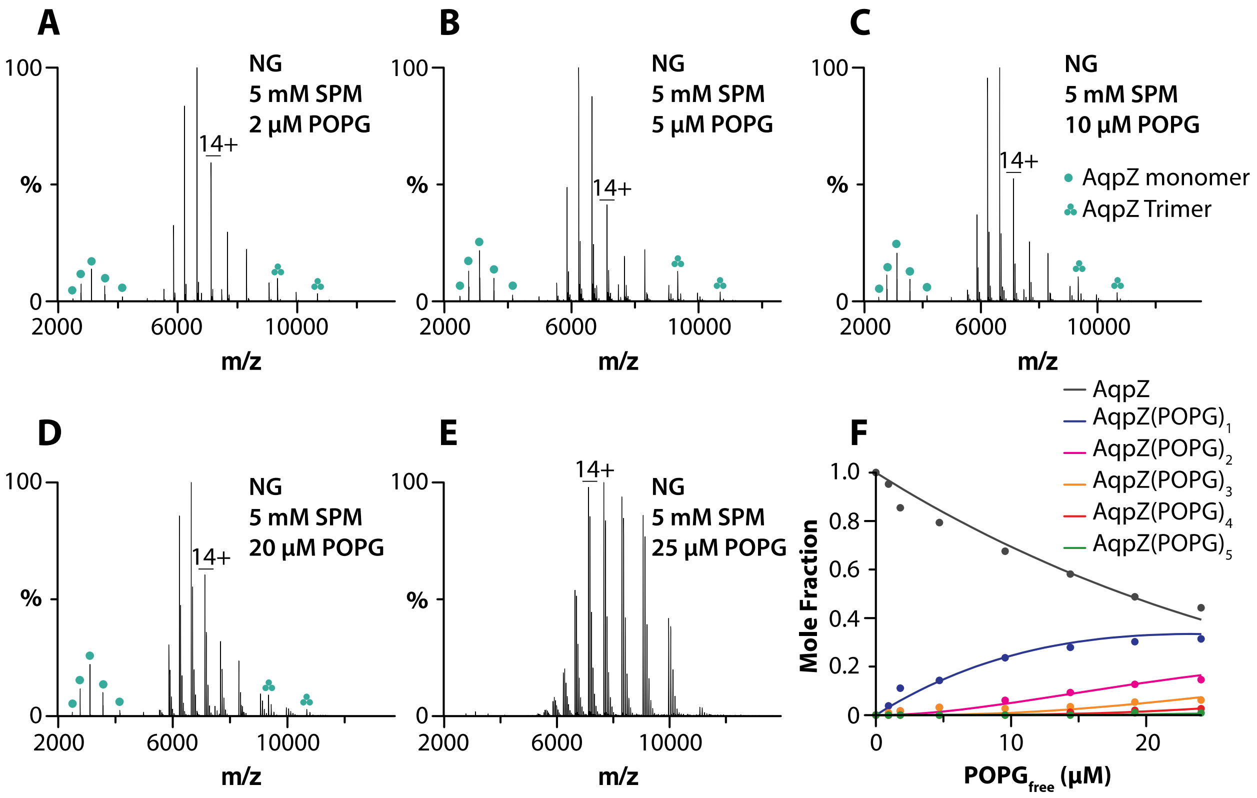


**Figure S37. Determination of AqpZ-POPG equilibrium binding constants.** A-E) AqpZ (1 µM) in NG and 5 mM SPM mixed with different concentrations of POPG. F) Plot of mole fraction data (dots) determined from a titration series of POPG and subsequent fit of a sequential lipid binding model (lines).

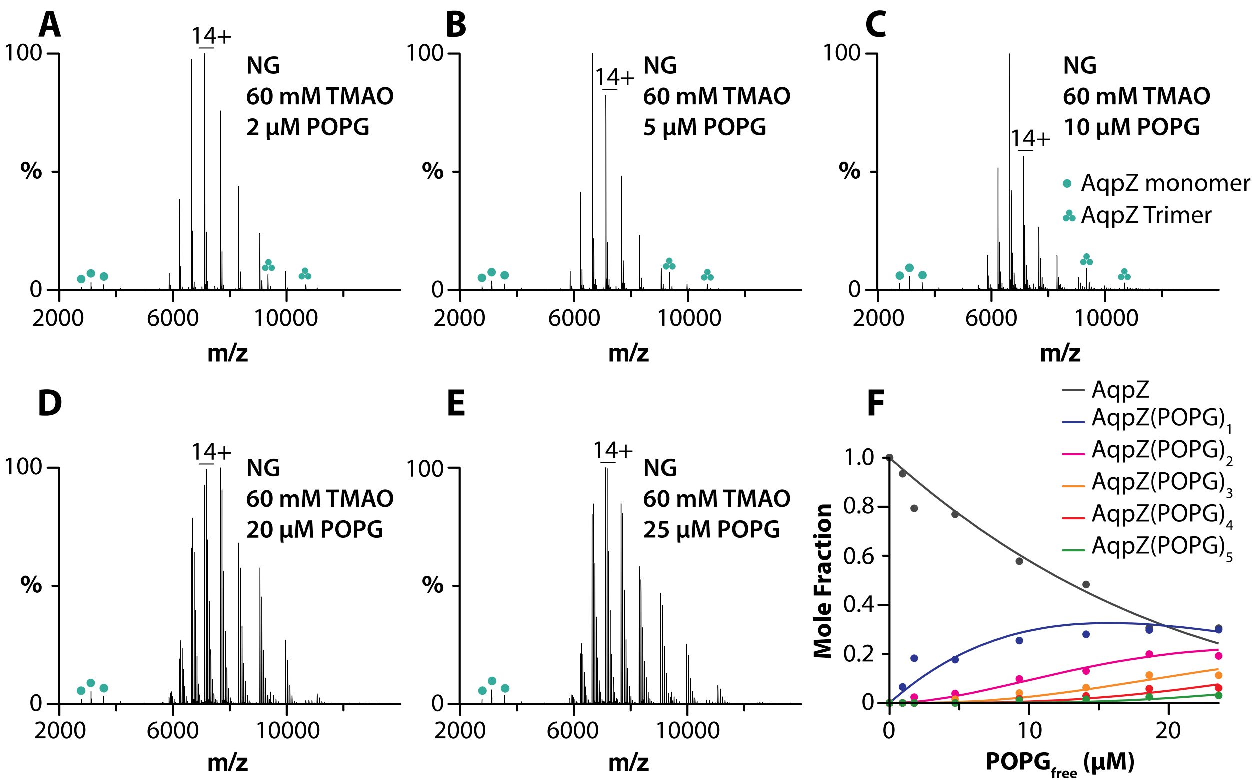


**Figure S38. Determination of AqpZ-POPG equilibrium binding constants.** A-E) AqpZ (1 µM) in NG and 60 mM TMAO mixed with different concentrations of POPG. F) Plot of mole fraction data (dots) determined from a titration series of POPG and subsequent fit of a sequential lipid binding model (lines).

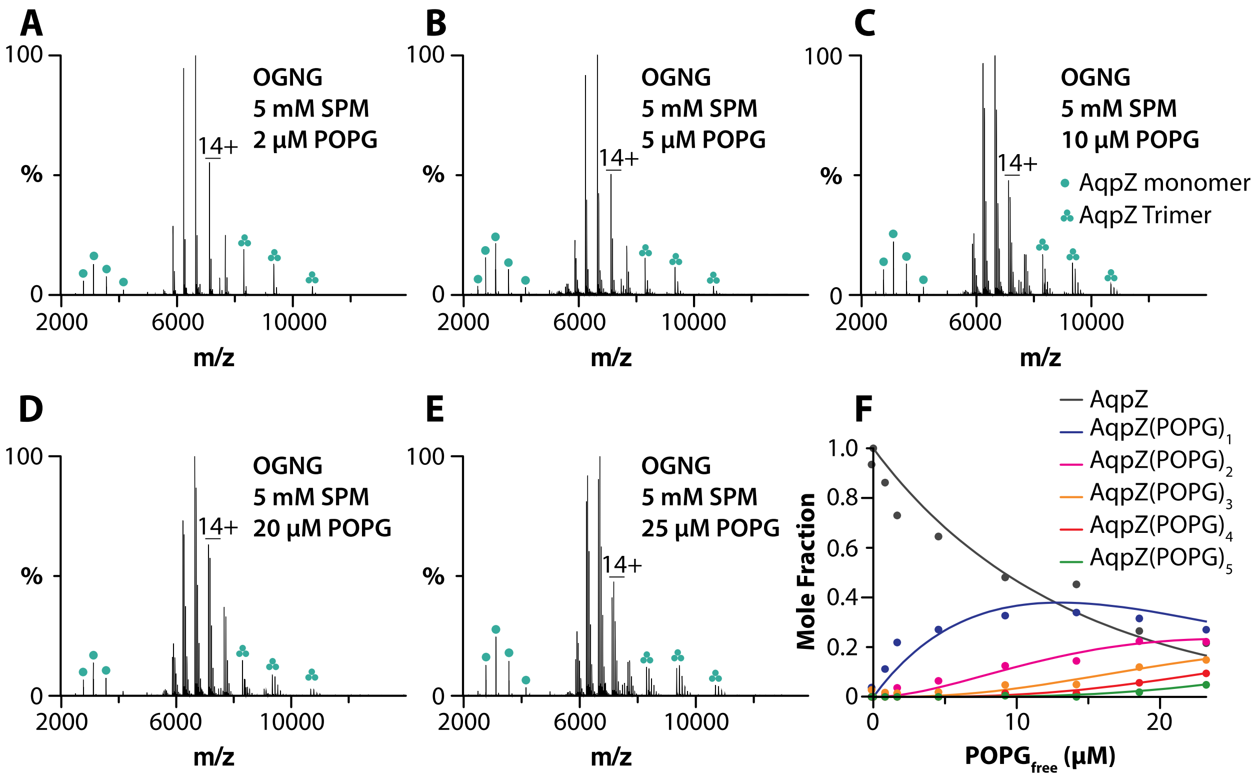


**Figure S39. Determination of AqpZ-POPG equilibrium binding constants.** A-E) AqpZ (1 µM) in OGNG and 5 mM SPM mixed with different concentrations of POPG. F) Plot of mole fraction data (dots) determined from a titration series of POPG and subsequent fit of a sequential lipid binding model (lines).

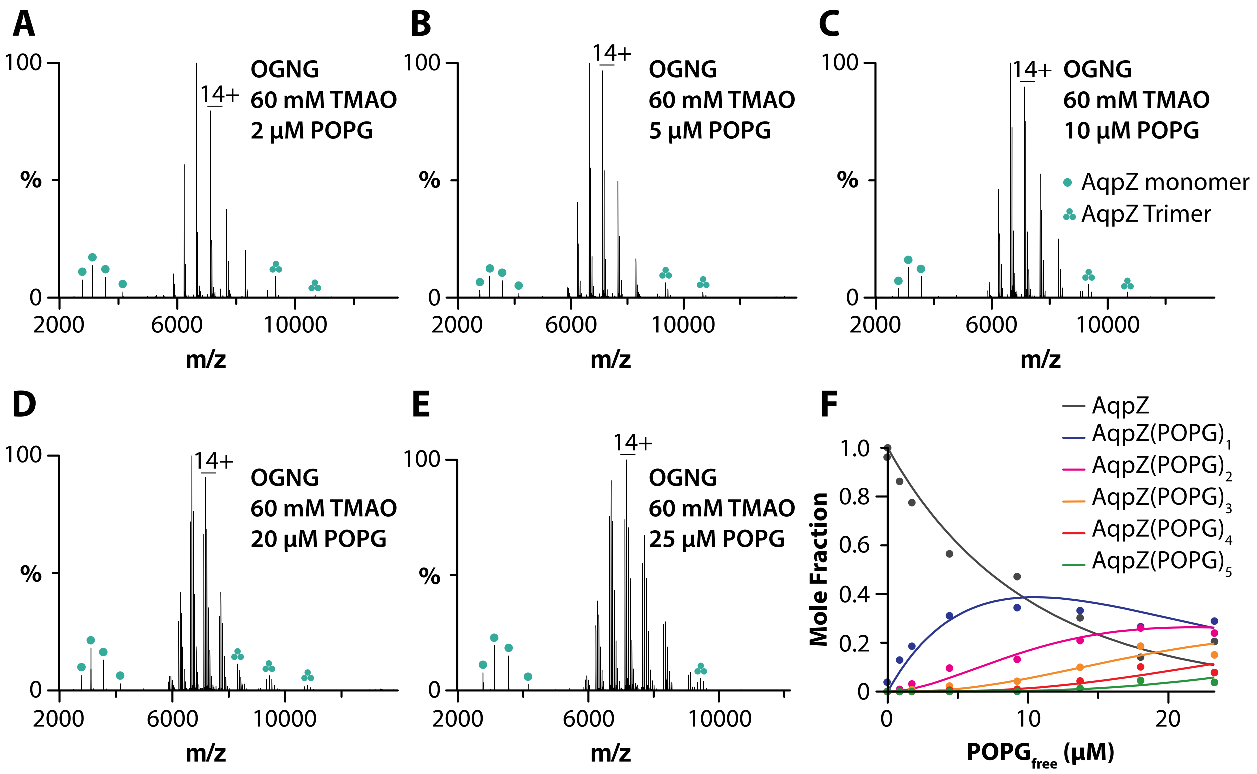


**Figure S40. Determination of AqpZ-POPG equilibrium binding constants.** A-E) AqpZ (1 µM) in OGNG and 60 mM TMAO mixed with different concentrations of POPG. F) Plot of mole fraction data (dots) determined from a titration series of POPG and subsequent fit of a sequential lipid binding model (lines).

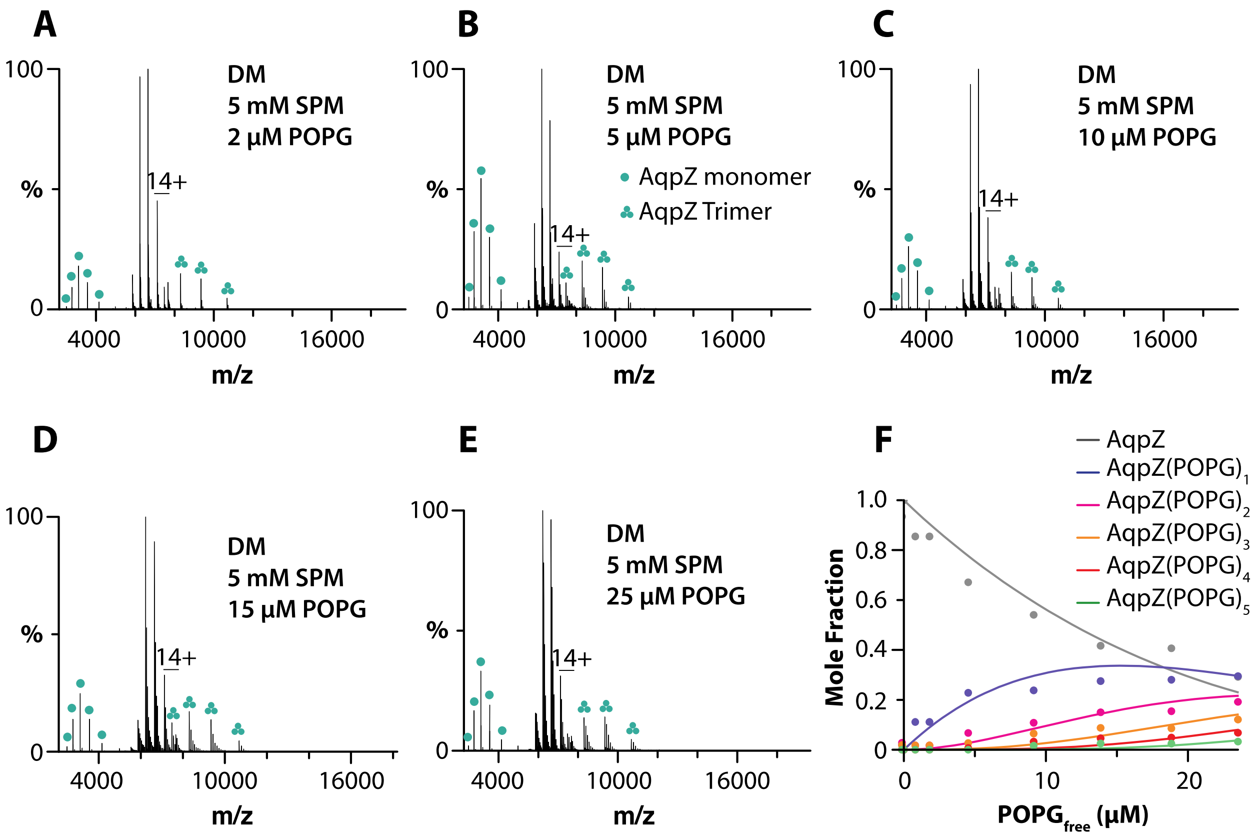


**Figure S41. Determination of AqpZ-POPG equilibrium binding constants.** A-E) AqpZ (1 µM) in DM and 5 mM SPM mixed with different concentrations of POPG. F) Plot of mole fraction data (dots) determined from a titration series of POPG and subsequent fit of a sequential lipid binding model (lines).

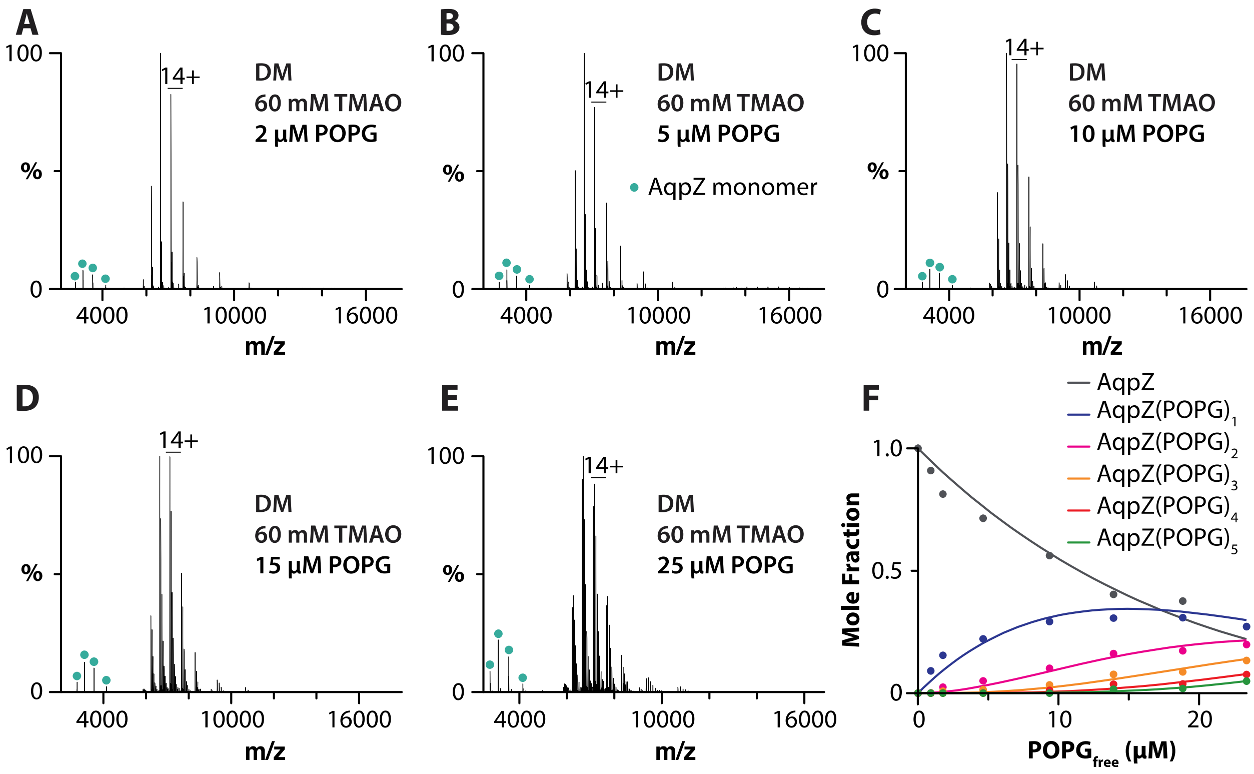


**Figure S42. Determination of AqpZ-POPG equilibrium binding constants.** A-E) AqpZ (1 µM) in DM and 60 mM TMAO mixed with different concentrations of POPG. F) Plot of mole fraction data (dots) determined from a titration series of POPG and subsequent fit of a sequential lipid binding model (lines).

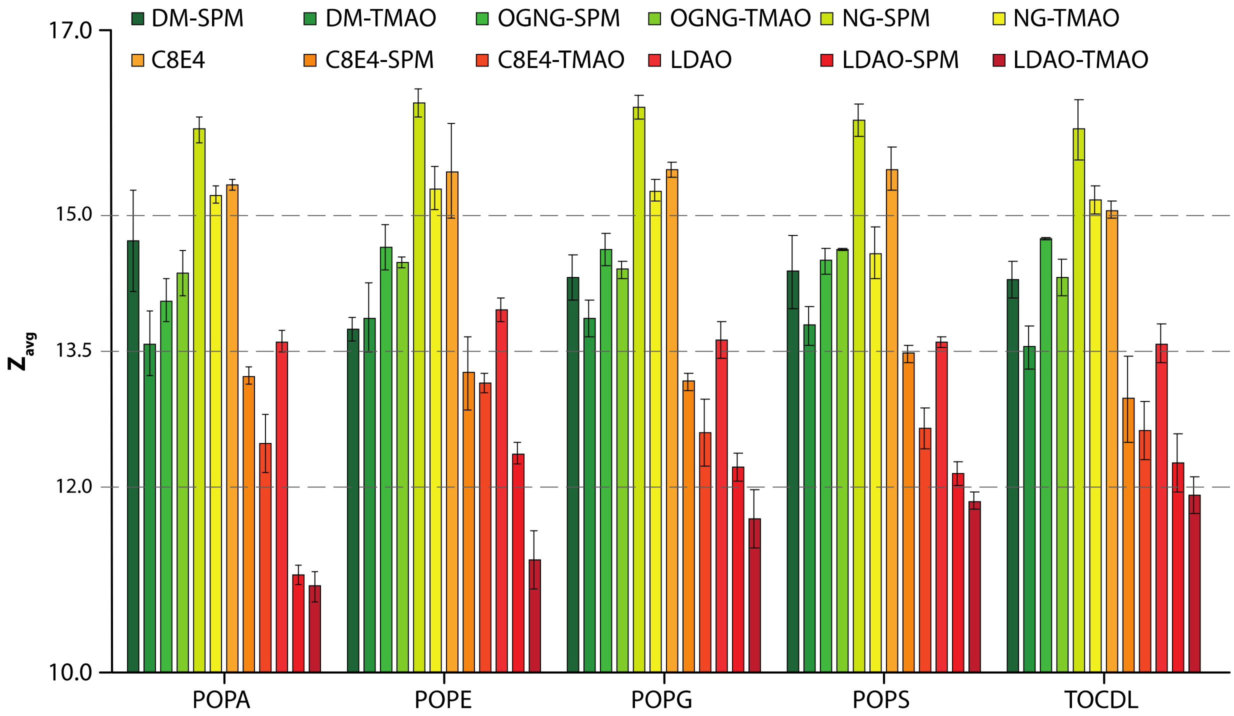


**Figure S43. $Z_{avg}$ for AqpZ in various detergents with different lipids.** Reported are the mean and standard deviation (*n*=3).

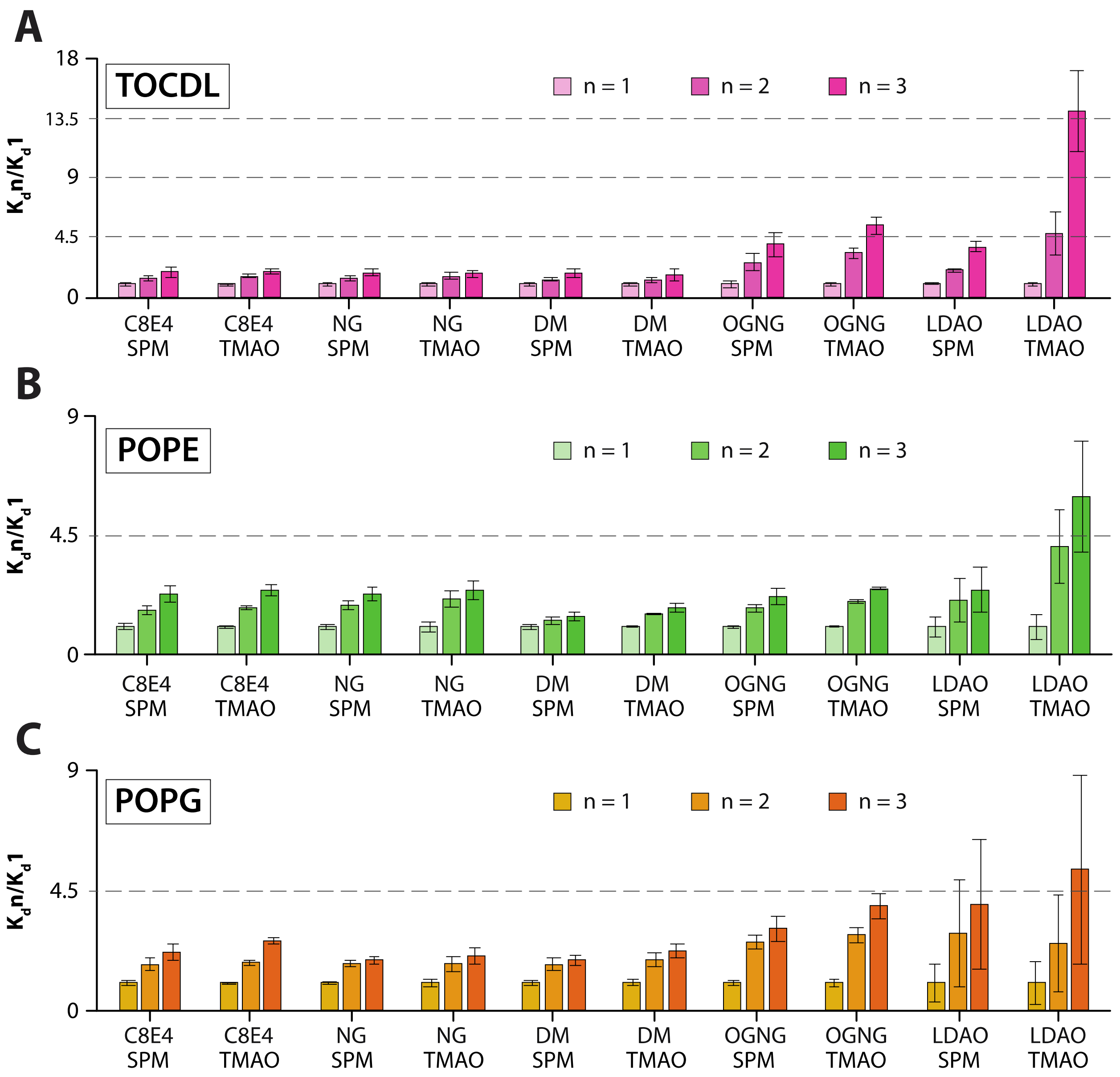


**Figure S44. Fold change in $K_d$s for subsequent lipid binding to AqpZ in different detergents.** Reported are the mean and standard deviation (*n*=3).

**Supporting Tables**

**Table S1. Instrument settings for analysis of AqpZ**.

| System | SID | HCD | Spray Voltage (kV) | Capillary Temperature (°C) | Trapping Pressure | Source DC Offset (V) | Injection Flatpole DC (V) | Inter Flatpole Lens (V) | Bent Flatpole DC (V) | Transfer Multipole DC (V) |
|---|---|---|---|---|---|---|---|---|---|---|
| **DM** | 100 | 100 | 1.50 | 300 | 6 | 40.0 | 8.0 | 4.0 | 3.0 | 3.0 |
| **DM-SPM** | 100 | 80 | 1.60 | 300 | 6 | 25.0 | 8.0 | 5.0 | 16.0 | 3.0 |
| **DM-TMAO** | 100 | 80 | 1.60 | 300 | 6 | 25.0 | 8.0 | 5.0 | 16.0 | 3.0 |
| **OGNG** | 80 | 100 | 1.50 | 300 | 7 | 60.0 | 4.0 | -20.0 | 10.0 | 6.0 |
| **OGNG-SPM** | 80 | 100 | 1.50 | 300 | 7 | 60.0 | 4.0 | -20.0 | 10.0 | 6.0 |
| **OGNG-TMAO** | 80 | 100 | 1.50 | 300 | 7 | 60.0 | 4.0 | -20.0 | 10.0 | 6.0 |
| **NG** | 60 | 70 | 1.70 | 200 | 5 | 25.0 | 8.0 | 7.0 | 6.0 | 2.0 |
| **NG-SPM** | 60 | 70 | 1.70 | 200 | 5 | 25.0 | 8.0 | 7.0 | 6.0 | 2.0 |
| **NG-TMAO** | 60 | 70 | 1.70 | 200 | 5 | 25.0 | 8.0 | 7.0 | 6.0 | 2.0 |
| **C8E4** | 60 | 70 | 1.70 | 200 | 5 | 25.0 | 8.0 | 7.0 | 6.0 | 2.0 |
| **C8E4-SPM** | 60 | 70 | 1.70 | 200 | 5 | 25.0 | 8.0 | 7.0 | 6.0 | 2.0 |
| **C8E4-TMAO** | 60 | 70 | 1.70 | 200 | 5 | 25.0 | 8.0 | 7.0 | 6.0 | 2.0 |
| **LDAO** | 60 | 70 | 1.70 | 200 | 5 | 25.0 | 8.0 | 7.0 | 6.0 | 2.0 |
| **LDAO-SPM** | 60 | 70 | 1.70 | 200 | 5 | 25.0 | 8.0 | 7.0 | 6.0 | 2.0 |
| **LDAO-TMAO** | 60 | 70 | 1.70 | 200 | 5 | 25.0 | 8.0 | 7.0 | 6.0 | 2.0 |

**Table S2. Average charge state ($Z_{avg}$) of AqpZ in different detergent environments**. Reported are mean and standard deviation (n=3).

| System | $Z_{avg}$ | # of charge states |
|---|---|---|
| **DM-SPM** | 14.5 ± 0.1 | 4.0 ± 0.0 |
| **DM-TMAO** | 13.5 ± 0.4 | 4.7 ± 1.7 |
| **OGNG** | 17.4 ± 0.5 | 6.0 ± 0.0 |
| **OGNG-SPM** | 14.7 ± 0.2 | 5.7 ± 0.5 |
| **OGNG-TMAO** | 14.9 ± 0.4 | 4.0 ± 0.0 |
| **NG** | 18.8 ± 0.1 | 5.3 ± 0.5 |
| **NG-SPM** | 16.5 ± 0.2 | 7.7 ± 0.5 |
| **NG-TMAO** | 14.5 ± 0.3 | 6.3 ± 0.5 |
| **C8E4** | 15.3 ± 0.0 | 5.0 ± 0.0 |
| **C8E4-SPM** | 13.2 ± 0.1 | 6.7 ± 0.5 |
| **C8E4-TMAO** | 13.0 ± 0.2 | 5.7 ± 0.5 |
| **LDAO** | 13.5 ± 0.1 | 6.7 ± 0.9 |
| **LDAO-SPM** | 11.5 ± 0.0 | 7.3 ± 0.5 |
| **LDAO-TMAO** | 10.3 ± 0.2 | 6.3 ± 0.5 |

**Table S3. Instrument settings for the analysis of AmtB-GlnK.** Reported are mean and standard deviation (n=3).

| System | SID | HCD | Spray Voltage (kV) | Capillary Temperature (°C) | Trapping Pressure | Source DC Offset (V) | Injection Flatpole DC (V) | Inter Flatpole Lens (V) | Bent Flatpole DC (V) | Transfer Multipole DC (V) |
|---|---|---|---|---|---|---|---|---|---|---|
| **DM** | 50 | 90 | 1.60 | 200 | 6 | 30.0 | 8.0 | 5.0 | 16.0 | 3.0 |
| **DM-SPM** | 50 | 120 | 1.60 | 200 | 6 | 35.0 | 8.0 | 5.0 | 16.0 | 3.0 |
| **DM-TMAO** | 50 | 120 | 1.60 | 200 | 6 | 35.0 | 8.0 | 5.0 | 16.0 | 3.0 |
| **OGNG** | 80 | 100 | 1.50 | 300 | 7 | 60.0 | 4.0 | -20.0 | 10.0 | 6.0 |
| **OGNG-SPM** | 80 | 100 | 1.50 | 300 | 7 | 60.0 | 4.0 | -20.0 | 10.0 | 6.0 |
| **OGNG-TMAO** | 60 | 60 | 1.60 | 200 | 5 | 60.0 | 4.0 | -20.0 | 10.0 | 6.0 |
| **NG** | 40 | 105 | 1.60 | 200 | 5 | 60.0 | 4.0 | -20.0 | 10.0 | 6.0 |
| **NG-SPM** | 40 | 100 | 1.60 | 200 | 5 | 60.0 | 4.0 | -20.0 | 10.0 | 6.0 |
| **NG-TMAO** | 70 | 80 | 1.60 | 200 | 5 | 60.0 | 4.0 | -20.0 | 10.0 | 6.0 |
| **C8E4** | 60 | 60 | 1.60 | 200 | 5 | 60.0 | 4.0 | -20.0 | 10.0 | 6.0 |
| **C8E4-SPM** | 60 | 60 | 1.60 | 200 | 5 | 60.0 | 4.0 | -20.0 | 10.0 | 6.0 |
| **C8E4-TMAO** | 60 | 60 | 1.60 | 200 | 5 | 60.0 | 4.0 | -20.0 | 10.0 | 6.0 |
| **LDAO** | 60 | 60 | 1.60 | 200 | 5 | 60.0 | 4.0 | -20.0 | 10.0 | 6.0 |
| **LDAO-SPM** | 60 | 60 | 1.60 | 200 | 5 | 60.0 | 4.0 | -20.0 | 10.0 | 6.0 |
| **LDAO-TMAO** | 60 | 60 | 1.60 | 200 | 5 | 60.0 | 4.0 | -20.0 | 10.0 | 6.0 |

**Table S4. $Z_{avg}$ of AmtB-GlnK in different detergent environments.**

| System | $Z_{avg}$ | # of Zs |
|---|---|---|
| **DM-SPM** | 21.9 ± 0.2 | 5.7 ± 0.5 |
| **NG** | 25.8 ± 0.2 | 5.0 ± 0.0 |
| **NG-SPM** | 22.6 ± 0.3 | 6.7 ± 0.5 |
| **C8E4** | 20.5 ± 0.1 | 4.0 ± 0.0 |
| **C8E4-SPM** | 18.8 ± 0.4 | 6.0 ± 0.8 |
| **C8E4-TMAO** | 17.3 ± 0.3 | 5.0 ± 0.0 |
| **LDAO** | 20.0 ± 0.3 | 8.7 ± 0.5 |
| **LDAO-SPM** | 17.7 ± 0.5 | 7.3 ± 0.5 |

**Table S5. Equilibrium dissociation constants ($K_D$s) for AqpZ-TOCDL in different detergents.** Reported are the mean and standard deviation (*n* = 3)

| Environment | $K_d$1 (μM) | $K_d$2 (μM) | $K_d$3 (μM) |
|---|---|---|---|
| **C8E4+5 mM SPM** | 30.3 ± 2.7 | 45.3 ± 4.0 | 58.3 ± 11.2 |
| **C8E4+60 mM TMAO** | 22.8 ± 0.9 | 37.0 ± 1.6 | 45.9 ± 4.0 |
| **NG+5 mM SPM** | 28.8 ± 3.3 | 42.3 ± 2.5 | 54.4 ± 4.6 |
| **NG+60 mM TMAO** | 19.7 ± 1.6 | 31.5 ± 4.1 | 35.5 ± 4.9 |
| **DM+5 mM SPM** | 21.7 ± 2.1 | 30.0 ± 2.8 | 39.5 ± 5.6 |
| **DM+60 mM TMAO** | 14.1 ± 1.8 | 18.9 ± 2.2 | 23.8 ± 5.1 |
| **OGNG+5 mM SPM** | 7.8 ± 1.7 | 20.9 ± 1.3 | 31.1 ± 0.9 |
| **OGNG+60 mM TMAO** | 4.4 ± 0.5 | 14.9 ± 0.6 | 23.9 ± 0.2 |
| **LDAO+5 mM SPM** | 1.4 ± 0.1 | 2.8 ± 0.1 | 5.2 ± 0.4 |
| **LDAO+60 mM TMAO** | 0.3 ± 0.0 | 1.6 ± 0.5 | 4.7 ± 0.9 |

**Table S6. $K_D$s for POPE binding to AqpZ in different detergents.** Reported are the mean and standard deviation (*n* = 3)

| Environment | $K_d$1 (μM) | $K_d$2 (μM) | $K_d$3 (μM) |
|---|---|---|---|
| **C8E4+5 mM SPM** | 28.2 ± 2.3 | 46.1 ± 2.4 | 63.2 ± 7.0 |
| **C8E4+60 mM TMAO** | 23.0 ± 0.5 | 39.3 ± 1.3 | 55.0 ± 5.0 |
| **NG+5 mM SPM** | 44.3 ± 3.0 | 80.2 ± 5.4 | 99.5 ± 9.7 |
| **NG+60 mM TMAO** | 30.7 ± 3.8 | 63.0 ± 5.2 | 73.4 ± 5.3 |
| **DM+5 mM SPM** | 37.8 ± 2.4 | 47.1 ± 4.5 | 52.4 ± 4.9 |
| **DM+60 mM TMAO** | 23.7 ± 0.3 | 35.3 ± 0.9 | 41.1 ± 3.6 |
| **OGNG+5 mM SPM** | 19.7 ± 0.7 | 33.6 ± 2.8 | 42.3 ± 6.1 |
| **OGNG+60 mM TMAO** | 15.1 ± 0.3 | 29.6 ± 0.7 | 37.0 ± 0.0 |
| **LDAO+5 mM SPM** | 4.2 ± 1.1 | 8.5 ± 2.5 | 10.1 ± 2.3 |
| **LDAO+60 mM TMAO** | 1.3 ± 0.4 | 5.2 ± 0.4 | 7.6 ± 0.7 |

**Table S7. $K_D$s for AqpZ-POPG in different detergents.** Reported are the mean and standard deviation (*n* = 3)

| Environment | $K_d$1 (μM) | $K_d$2 (μM) | $K_d$3 (μM) |
|---|---|---|---|
| **C8E4+5 mM SPM** | 25.8 ± 2.0 | 43.5 ± 4.9 | 55.5 ± 6.9 |
| **C8E4+60 mM TMAO** | 18.4 ± 0.1 | 32.2 ± 1.8 | 47.3 ± 2.3 |
| **NG+5 mM SPM** | 27.0 ± 1.2 | 46.3 ± 2.9 | 49.8 ± 2.7 |
| **NG+60 mM TMAO** | 19.0 ± 1.8 | 32.4 ± 4.6 | 38.3 ± 4.7 |
| **DM+5 mM SPM** | 19.9 ± 1.2 | 33.7 ± 4.0 | 36.8 ± 2.8 |
| **DM+60 mM TMAO** | 15.5 ± 1.3 | 28.8 ± 3.1 | 34.0 ± 2.5 |
| **OGNG+5 mM SPM** | 12.0 ± 0.6 | 30.1 ± 2.5 | 36.0 ± 5.1 |
| **OGNG+60 mM TMAO** | 8.8 ± 0.8 | 24.5 ± 1.1 | 34.0 ± 2.6 |
| **LDAO+5 mM SPM** | 2.0 ± 1.0 | 5.6 ± 2.8 | 7.8 ± 2.8 |
| **LDAO+60 mM TMAO** | 1.2 ± 0.7 | 2.8 ± 1.3 | 6.0 ± 2.1 |

**Table S8. CMC of different detergents used in this study.**

| Detergent | CMC (wt/wt) |
|---|---|
| **DM** | 0.087 |
| **OGNG** | 0.058 |
| **NG** | 0.2 |
| **C8E4** | 0.25 |
| **LDAO** | 0.023 |
